# DeMoDa: A global open access database for comparative research in human dental morphological variation

**DOI:** 10.64898/2026.05.30.728952

**Authors:** Marina Martínez de Pinillos González, Ana Álvarez Fernández, Beatriz Delgado Esteban, Miguel Delgado, Laresa L. Dern, Joel D. Irish, Osamu Kondo, María Martinón Torres, Mario Modesto-Mata, G. Richard Scott, Arthur Thiebaut, Kathleen S. Paul, Marin A. Pilloud, Hannes Rathmann, Hugo Reyes-Centeno, Tatiana Vlemincq-Mendieta, Leslea J. Hlusko

## Abstract

Human dental morphology is diverse and varies both within and between populations worldwide. Common variants include different numbers of cusps and roots, as well as different configurations in the fissures, ridges, and grooves on tooth crowns. Because teeth preserve well in taphonomic contexts and retain strong genetic signatures in their morphology, researchers across disciplines use dental form for research ranging from population affinity identification in forensic cases to the reconstruction of population history in archaeological and paleontological studies, and to explore the genetic underpinnings of dental development.

However, these analyses are limited, as no publicly available comprehensive database of human dental morphological variation currently exists. Data are typically shared only within professional networks, excluding scientists outside those circles. Information is dispersed across repositories and the supplementary materials that accompany peer-reviewed publications, creating a fragmented and difficult-to-navigate data landscape. Although numberous dental anthropologists devoted their careers to collecting extensive datasets from thousands of individuals worldwide, their data have not yet been published in raw form nor made compatible.

Here, we introduce the Dental Morphological Database (DeMoDa), an open access repository comprising 246 dental traits for 17,308 individuals across 32 major geographic regions worldwide spanning the past several thousand years. These data are from the legacy datasets of Turner, Hanihara, Scott, and Irish. Because these researchers used different scoring systems, we developed a scoring system with two steps that enables the three most-common scoring systems to be combined. We provide two versions of the data that correspond to these two steps: one with more nuanced trait scores at the cost of a smaller sample size, and another with less nuanced trait scores but broader sample coverage. We discuss the implications of publishing these legacy data and outline our decision-making process that guided their release in accordance with both the FAIR and CARE principles of open science.

## I. Introduction

The practice of science in the 21^st^ century is distinct from previous praxis in two ways. First, thanks to the increased analytical and storage capacities of computers and the ubiquitous use of the internet as a scientific infrastructure, scientists can develop, access, and analyze large datasets in ways that were not previously feasible. Second, scientists now generally share their raw data, facilitating larger, “big data” analyses. Anthropology is no exception to this data revolution (e.g., Douglas-Jones et al., 2021; Green, 2023; Paff, 2022).

Within anthropology, the subdiscipline of dental anthropology was founded on the idea that large datasets documenting human dental variation could yield new insight into origins and patterns of human migration (Scott and Turner II, 1997). However, this dental variation was difficult to capture using simple methodologies because the features are continuously variable and 3-dimensional. Several different researchers defined traits and developed scoring systems to turn them into quasi-quantitative data (Hanihara, 2008; Scott and Irish, 2017; Turner et al., 1991). With tens of thousands of individual dentitions representing people from all around the world, anthropologists have used the variation observed for these traits to reconstruct population history and migration, evaluate biological affinities among ancient and modern groups, examine hominin evolutionary relationships, and investigate developmental and genetic mechanisms underlying human variation (Scott et al., 2018). Consequently, dental morphology represents one of the most informative skeletal systems available for studying both microevolutionary and macroevolutionary processes in our species.

Until fairly recently, sharing data in the anthropological sciences was logistically complicated. The different scoring systems and peculiarities of a heterodont dentition lead to datasets that are unusual and quirky (as they are a mix of categorical, nominal, discrete, and continuous values across 32 teeth that fall into 8 tooth type categories in two jaws). Additionally, until recently, there was no official repository to affirm and ensure data comparability and integrity during the sharing process. When data were shared, a file was passed directly between individuals. This manner of sharing is less than ideal because the more times a file passes hands, the greater the chance that inconsistencies, errors, and misunderstandings are introduced. Additionally, there was no clear mechanism for guaranteeing that the data were authentic, and no way to formally credit the person who collected the original data. And of course, data were only shared within professional networks, excluding scientists from outside. As previous merit systems rewarded data-sharing primarily through peer-reviewed research publication, the large datasets developed in dental anthropology were presented in scientific publications through summary tables and appendices that were feasible to present, maintained accuracy, and could be easily cited (e.g. Scott and Turner II, 1997).

Simultaneous to this research in anthropology over the last two decades, we have begun to understand the biological underpinnings of dental variation (Novacescu *et al*., 2025). It is increasingly evident that some of the dental morphological variants studied in anthropology are influenced by variation in at least one gene pathway that effects a wider spectrum of our anatomy and physiology far beyond the dentition (the ectodysplasin pathway, e.g., (Adhikari et al., 2016; Coletta et al., 2021a; Hlusko et al., 2018; Kataoka et al., 2021; Kimura et al., 2009; Li et al., 2025; Morandini et al., 2025; Wu et al., 2026). These discoveries demonstrate that the variation recorded by dental anthropologists provides a unique window into our species-level variation of broader physiological phenomena. However, research probing variation in the ectodysplasin pathway using dental morphological scores is limited by the inaccessibility of the individual-level data, and a lack of access to datasets large enough to provide statistical robusticity. In order to fulfill the scientific potential of these data, we need to make available the individual-level scores and combine the efforts of different researchers despite their different data collection methodologies and scoring systems.

There have been efforts to compile data from across these published studies for meta-analyses. For example, Rathmann *et al*. (2017) used the dental trait dataset compiled by Tsunehiko Hanihara to test the assumption that these traits are selectively neutral, while Irish *et al*., (2020) conducted a broader study using data collected by Christy G. Turner II, G. Richard Scott, and Joel D. Irish. In follow-up research, Rathmann and Reyes-Centeno, (2020) analyzed millions of combinations of dental traits derived from datasets compiled by Turner and Scott, identifying a set of highly diagnostic features that most effectively preserve underlying neutral genetic signals that permit the tracking of population movement in the deep past. In a subsequent study, this time using a combined dataset collected by Turner, Scott, and Irish, Rathmann *et al*., (2023), demonstrated that dental morphological traits are among the most informative markers in the human skeleton for inferring neutral variation, significantly outperforming other data types commonly employed in anthropology, such as cranial traits and dental linear measurements.

More recently, Irish et al. (2026) used dental morphological trait frequencies, matched genomic data, and minimum-slope geographic distances to show that these traits parallel neutral genetic structure in both recent populations and Early through Late Holocene groups. These results support their use for reconstructing population history within Africa and during the Late Pleistocene Out-of-Africa II dispersal. Qaq et al. (2026) surveyed the literature and compiled dental morphology scored using the Arizona State University Dental Anthropology System (ASUDAS) (Scott and Irish, 2017; Turner et al., 1991), combining data from 43 studies for a dataset of 36,919 individuals (the authors note that there may be duplication of individuals in their dataset given their reliance on summary tables). Qaq et al. (2026) organized the data into five major populations and focused their analyses on testing the robusticity of these divisions: Sub-Saharan Africa, Western Eurasia, Sino-Americas, Sunda-Pacific, and Sahul-Pacific. Although their results somewhat support this continent-level categorization, the authors point to the fact that the significant amount of overlap between the samples undermines the value of studying human variation in this manner. Ultimately, they call for “research into the biological and developmental influences on dental morphology” (2026: 8).

Clearly, there is significant untapped potential for uncovering the biological underpinnings of dental variation in these published data. However, the challenges are to combine data from across researchers and make the data available in their original form. While the original dental morphological scores were recorded as quasi-continuous traits, they typically were published as dichotomized presence-absence traits (i.e. binary data), thus losing information about variation in trait expression among samples. Additionally, no one has yet combined these original quasi-continuous data from Hanihara with those collected by Turner, Scott, and Irish. That said, it is important to acknowledge that dichotomized data and publications using them do have several advantages: they can further reduce inter- and intra-observer error, provide higher average heritability estimates, i.e., almost twice those reported when traits are treated as continuous variables (Stojanowski et al., 2019), and avoid weighting bias from different numbers of grades across ASUDAS traits, which range from two to eight (Scott and Irish, 2017). These strengths make dichotomized datasets valuable, but they also underscore the need for a resource that preserves original scores while allowing different recording systems to be compared.

Here, we present a major advancement to the study of human dental variation: the Dental Morphological Database (DeMoDa). We developed DeMoDa to host original historical dental anthropological data collected by a few prolific researchers. In addition to developing a repository for the original data that was published in various formats and configurations, we created a scoring system that enables data collected via different scoring systems to be combined. The DeMoDa scoring system consists of two steps that translate three of the most widely used scoring systems into one. These two innovations enable the publication of the largest dataset of individual-level human dental variation to date: 247 traits scored for 17,308 individuals from 32 geographic regions that cover the world and span several thousand years. The database, the two scoring systems, the dataset, and the thought processes that guided decisions about their presentation format are described herein.

## II. The Dental Morphological Database (DeMoDa)

The Dental Morphological Database (DeMoDa) is an online repository for individual dental morphological trait scores. It is designed to accommodate the original data collected by four prolific dental anthropologists but can be expanded to include data collected by other researchers in the future. DeMoDa was developed to preserve these historical data, provide a framework to integrate them, and to make them open access, thereby advancing our understanding of world-wide human dental variation. The DeMoDa was explicitly designed with a commitment to Open Science, in accordance with the FAIR Principles (https://www.go-fair.org/fair-principles/) and the Open Science initiative of the ERC Horizon Europe program that provided the funding through Grant agreement ID: 101054659, Tied2Teeth. The FAIR Principles call for digital assets and research data to be <u>F</u>indable, <u>A</u>ccessible, Interoperable, and <u>R</u>eusable (FAIR).

To ensure that the data in DeMoDa are thoughtfully managed in the immediate- and long-term, two entities have been tasked with responsibility and stewardship. The most proximal is the Scientific Advisory Board (SAB) with twelve members (all are included as coauthors) charged with managing the details and logistics of DeMoDa. The SAB members represent eight countries across four continents, bringing to DeMoDa a range of scientific and societal perspectives. The principal investigator of the European Research Council’s grant supporting this project, L.J.H. initiated the design of the SAB and invited the members of the inaugural configuration. The first meeting of the SAB was held in Burgos, Spain, on June 2-4, 2025. During this meeting, the SAB developed the policies and procedures for DeMoDa (see **Appendix 1**). Going forward, the SAB membership will be determined in accordance with those procedures.

To ensure the higher-level, long-term stewardship of the database, the DeMoDa SAB leadership is developing a Memorandum of Understanding with the Dental Anthropology Association (DAA). The project was presented to the DAA membership during the 2026 annual business meeting in Denver, Colorado. There will be an upcoming vote on the inclusion of the DAA President as a permanent member of the SAB moving forward and enshrined in the DAA constitution.

## III. Overview of the historical data compiled in DeMoDa

DeMoDa hosts dental morphological scores from permanent teeth of adults collected with the three most-used dental anthropology scoring systems (Hanihara, 2008; Scott and Irish, 2017; Turner et al., 1991) and data collected by the scientists who developed those systems (Tsunehiko Hanihara, Joel Irish, Christy G. Turner II, and G. Richard Scott). These data were collected specifically to explore human migration. Consequently, the researchers prioritized observations of the dentitions of people who had been long-term residents in their geographic region. These data fall under the category of “legacy data” as they were collected under past standard practices and have been extensively published in the scientific literature.

Standards and protocols for data collection from human remains in biological anthropology are constantly shifting. A detailed consideration of these shifts in practice guided the decisions we made for the presentation of the legacy data in DeMoDa. We first present an overview of the benefit to society that can be derived from these legacy data and the general pros and cons to open science. We then provide an overview of the legal and ethical contexts that we considered in our decision-making process for how to make these dental morphological data open access, the decision to create datasets rather than give free access to the main database itself (DeMoDa Dataset Version 1), the data included in the first dataset created, and a description of the DeMoDa scoring system that made it possible.

## IV. Balancing the societal benefit of the data in DeMoDa against its potential harm

The SAB set two guiding principles for the Dental Morphological Database (DeMoDa):

> **1. These data have value in known and as-yet-unknown scientific research areas that provide significant benefit to society. The data should be made freely-available in a manner that facilitates this research.**

Although DeMoDa is not a presentation of new data, we are presenting these legacy data in a new manner –the original scores for each individual rather than in summary tables by population. This reformatting will enable a much more nuanced exploration of variance and covariance within human dental variation. These historical data, presented in this updated format, provide a unique and important opportunity to dramatically improve studies of human dental variation at the global scale, with relevance to biological, biomedical, and evolutionary research, and as well as contributions to pedagogy and improving the public’s understanding of human biological variation.

The potential benefits to society include **new insights to human physiology**, as the genetic influences on dental variation also influence other aspects of our anatomies. For example, Hlusko et al. (2018) leaned on dental morphological data to identify the episode of intense evolutionary selection on the V370A allele of the *EDAR* gene that occurred in the Arctic region during the last ice age. With this geographic region identified, it was proposed that the very low levels of ultraviolet radiation exposure likely led to challenges in vitamin D sufficiency, especially for infants. Consequently, *EDAR V370A*’s effects on mammary glands could well have been the target of selection (Hlusko *et al*., 2018). This insight has contributed to the development of new hypotheses about human biology that range from lactational biology (e.g., (Zhang et al., 2022), to fatty acids (e.g., (Mitina et al., 2026), to inflammation response (Hlusko and McNelis, 2022), metabolic disease (Coletta et al., 2021), and possibly even risk of breast cancer (Tikhonov, 2022), not to mention the potential for selection of genetic variation of *EDAR* in other mammals (Font-Porterias et al., 2022).

Another likely benefit relates to the development of **new methods in forensic anthropology** and the tracing of the origins of isolated and marginalized populations, such as Afro-Colombians. Delgado-Burbano, (2007) investigated both the origins and the level of gene flow in an isolated Afro-Colombian community on the Colombian Pacific coast. By investigating dental nonmetric traits, this study identified African ancestors unrecognized in historical records, later confirmed by DNA studies, and provided evidence of low levels of gene flow with neighboring Native American and mestizo populations. Delgado *et al*., (2019) and (Yang *et al*., (2023) investigated dental metric and nonmetric diversity in living Latin American populations and, pairing these dental data with genomic data, developed methods for reconstructing biological ancestry and patterns of gene flow. Likewise, these studies provided a more detailed analysis of how dental metric and nonmetric traits can be used to investigate age and sexual dimorphism within the same population samples. Notably, the ability to estimate sex and biological affinity using alternative approaches is especially important in regions with histories of violence. Similarly, the reconstruction of the population history of isolated communities has also contributed to the self-recognition of their ancestors and promoted processes of identity reconstruction.

The information preserved in DeMoDa also **counters historical typological studies** that may have contributed to misconceptions and, in some cases, social injustices. In many anthropological investigations of skeletal variation based on large datasets, the true variability within populations was traditionally often collapsed into a single summary statistic of central tendency, and these group centroid estimates were then treated as representative of entire populations for comparative purposes. For example, in **Figure 1**, we demonstrate how this approach artificially emphasizes the differences between samples by comparing the dichotomized population frequencies next to the non-dichotomized trait data for individuals collected by Tsunehiko Hanihara. The individual-level data provide a much more accurate presentation of the immense overlap in phenotypic variation that exists within our species, variation that is continuous and beautifully complex. The individual-level data provided by DeMoDa will serve as a foundational comparative resource for a growing body of research that emphasizes individual-level rather than group-level approaches (e.g., Rathmann *et al*., 2019; Rathmann, Stoyanov y Posamentir, 2022; Mardini *et al*., 2023; Rathmann, Lismann, *et al*., 2023; Smith-Guzmán *et al*., 2025), which will facilitate a move away from race-based typological research.

**Figure 1.**
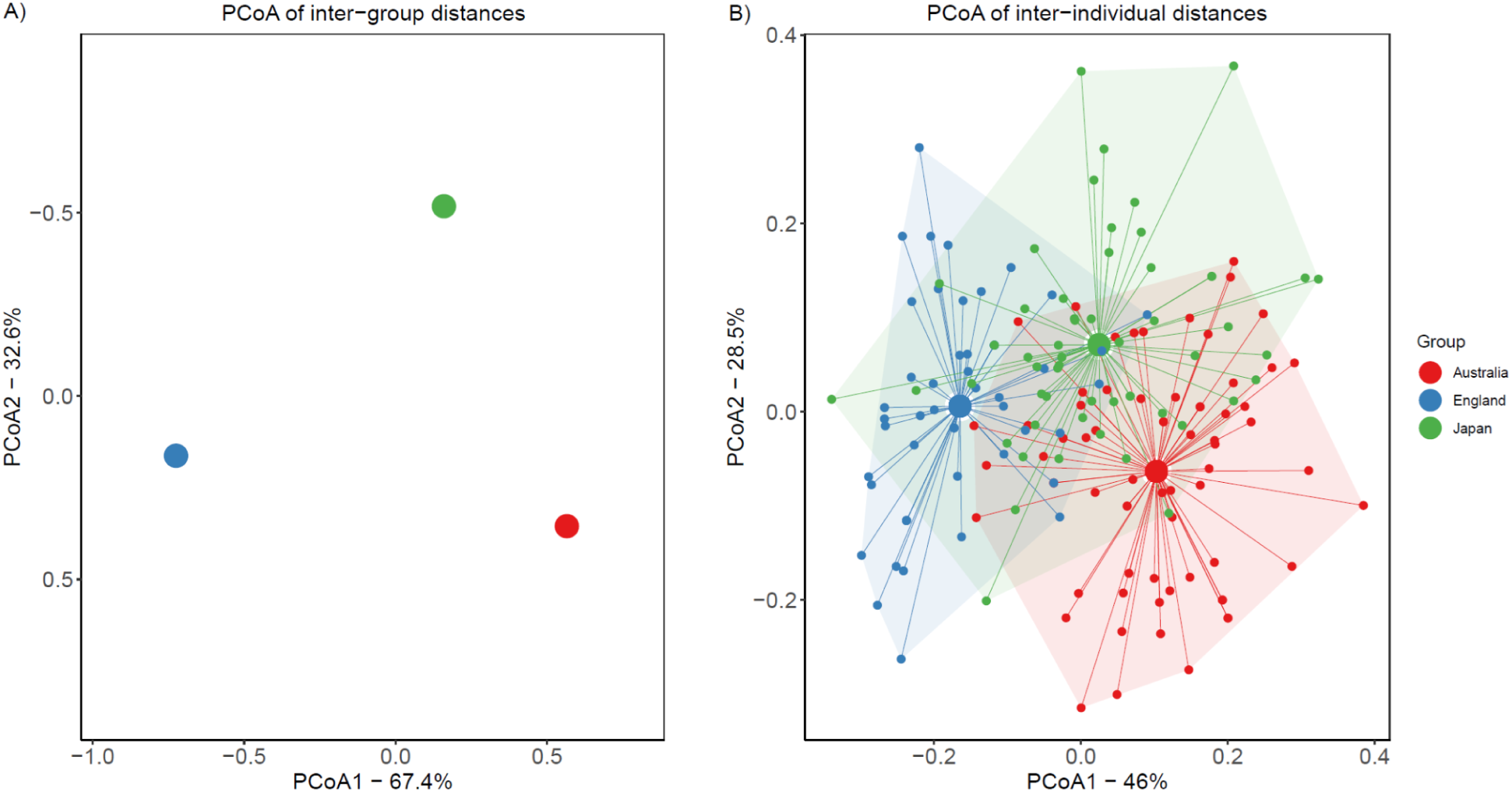
Exemplary contrast in the information content obtained through conventional group-level analyses routinely employed in dental anthropology versus individual-level analyses. (A) Principal Coordinates Analysis (PCoA) of inter-group distances calculated from dichotomized dental trait frequencies. (B) PCoA of inter-individual distances estimated from non-dichotomized raw trait data. For demonstration purposes, three geographically distinct population samples from the Tsunehiko Hanihara dataset were selected: Australia (red), England (blue), and Japan (green).

The more accurate demonstration of the complexity of human dental variation provided by the individual-level data in DeMoDa is particularly powerful when applied to pedagogical settings, where global-scale data help **materialize and visualize complex microevolutionary processes for students, dentists, and other health care professionals.** DeMoDa’s online mapping function (see Section XIX) represents a critical teaching tool that can illustrate clinical trends as the product of long-term genetic drift and gene flow in humans. On the one hand, geographically structured patterns of dental morphological traits can inform decisions in orthodontic practice (e.g.,Fasale, Rao y Panwar, 2024; Adel *et al*., 2025). At the same time, the mapping output illustrates *variation within* populations in the form of expression pie charts (as opposed to just a single frequency value) helps students move past problematic categorization of human groups, to instead grasp the complexities of variation in *Homo sapiens*, as well as the processes driving that variation. As demonstrated in an initial pedagogical study, the addition of DeMoDa to classroom activities has the potential to increase students’ confidence in collecting and, perhaps more importantly, *interpreting*, dental morphological data (Paul et al., in prep).

> 2. **The foreseeable potential harm from making these data open access is in the pursuit of research on specific, vulnerable communities. Therefore, these data should be made available with enough contextual information to facilitate large-scale scientific research questions into human variation, but not include details that would enable research at localized geographic or cultural levels.**

These anonymous non-biological data by themselves do not cause harm (unlike DNA that can be connected to specific individuals), rather, it is the narratives and stories that may be told with them about specific communities (Borgelt et al., 2025). Therefore, the SAB decided that the most benefit DeMoDa can provide to society is through an open access/FAIR presentation of the individual level-data that enables innovative research and, at the same time, presents these data with an additional layer of anonymity so that research on any specific marginalized community is not possible. This means that the data are presented more specifically at the individual level, but with very little contextual information about each of the individuals. By de-sensitizing the contextual information, the presentation of the data protects descendants’ right to control the data derived from the ancestral communities to which they are connected, as more specific population-focused projects are not possible.

These two guiding principles follow the ethical balance between beneficence and justice, maximizing possible societal benefits and minimizing possible harms, while ensuring that both the benefits and burdens of research are distributed fairly (e.g., as established long ago in The Belmont Report, (Department of Health, Education, and Welfare, 1979). The details of how we settled on the specific aspects of this balance for the dental morphological data in DeMoDa, and the logistics required to achieve it within the framework of open science are discussed in the subsequent sections.

## V. The pros and cons of publicly-funded open science

Governments invest heavily in scientific research recognizing that knowledge is a public good (Boulton, 2025; Taylor, 2016), and consequently, public investment in science continues to grow. For example, the United States increased investment in research and development by more than 57% between 2000 and 2009; China did so by almost 740% (Wang et al., 2012). When taxes fund research, the products produced through and by that investment belong to the public and should be in the public domain (i.e., publicly-funded science should be open science, (Suber, 2003).

A suite of three papers recently surveyed the economic, societal, and academic impacts of open science (Cole et al., 2024; Klebel et al., 2025; Tsipouri et al., 2025). Open science (OS) is commonly defined as,

> “*[E]fforts by researchers, governments, research funding agencies or the scientific community itself to make the primary outputs of publicly funded research results–publications and the research data–publicly accessible in digital format with no or minimal restriction as a means for accelerating research*” (Organisation for Economic Co-operation and Development (OECD), 2015): 7).

UNESCO has stated that OS “has transformative potential…for reducing the existing inequalities in [science, technology and innovation] and accelerating progress towards the implementation of the…Sustainable Development Goals and beyond” (UNESCO, 2021: 2). In a review of the data to assess movement towards these goals, Tsipouri et al. (2025) found strong empirical evidence that open/FAIR data do provide efficiency gains, enhance innovation, and foster economic growth, especially in the life sciences. The positive societal impacts include education, science awareness, environmental and climate awareness, and social engagement, but have led to only limited improvements in public trust in academic research (Cole *et al*., 2024). The effects of open/FAIR data on the academy appear to be more mixed for a variety of reasons (Klebel et al., 2025). Data were re-used for reproducibility studies mostly when authors also shared their analytical coding in addition to their data. There are distinct differences between academic disciplines regarding the value placed on sharing and reusing data. Additionally, there are significant differences in the resources available to utilize those data between the global North and global South. Klebel et al., (2025) also found that the slow rate of academic publishing leads scientists in low- and middle-income countries to hesitate sharing data out of fear of being scooped by better-resourced investigators. And, of course, there are drawbacks. In terms of consent, research participants generally provide consent for very specific uses of their personal data, complicating the ability to reuse those data to address other research questions. Additionally, the people whose data is being shared are not always the people who benefit (Serwadda et al., 2018). We also recognize that institutions that fund this research do not always have the best interest of the research participants in mind, particularly when they are marginalized groups.

As we developed the DeMoDa open science resource for dental morphological data, we took under advisement these benefits and concerns about open science generally. In the sections below, we focus in on these pros and cons within the specific context of reusing dental morphological data collected from anonymous human skeletal remains.

## VI. Changing practices in biological anthropology

The data in DeMoDa were collected from human skeletal remains in museum and other institutional collections. The different histories of how human remains were incorporated into these collections have increasingly raised concerns surrounding the lack of consent inherent to most institutional skeletal collections (Robbins Schug et al., 2025; Sholts, 2025; Stantis, et al. 2025) and the push to decolonize museums more broadly (Wali and Collins, 2023). Some groups of people in these collections were incorporated as a result of serious social inequities and injustices, ranging from societal neglect to racism and, in some instances, events that could fairly be described as genocide (Fforde et al., 2020). Forensic anthropologists, anatomists, biological anthropologists, archaeologists, and dental anthropologists are investing considerable effort to re-envision our science going forward to ensure ethical practices that allow for trust to be built with descendant communities who carry trauma from these histories.

A recent shift in practice stems from discussions about the treatment of human skeletal remains (Auerbach et al., 2026; Nash and Colwell, 2020), responses to DNA and ancient DNA research (Bader et al., 2023; Claw et al., 2018; Tallbear, 2013), conversations about the museum display of human remains (Francis et al., 2025) and the need for better collection management (Lewis et al., 2026), the use of unprofessional or derogatory academic presentation titles (Passalacqua et al., 2014), and discussion about data-sharing and the reuse/retrospective analysis of published data (Boyer et al., 2020; Mulligan et al., 2022; Tsosie et al., 2021; Turner and Mulligan, 2019). Recognizing that community ethics and cultural attitudes vary across the world and between professional contexts, several scholars and practitioners have documented country- and region- level shifts in the standards developed for working with human remains, pointing to historical sociopolitical contingencies and recurring hegemony. For example, Grupe and Wahl (2018) review shifts in Germany linked to its history of imperialism and (de)colonization, National Socialism, geopolitical division, and reunification. Considering the hegemony of practices developed by English-speaking communities, Haque et al. (2026) review the diversity of traditions surrounding human remains in Southeast Asia and problematize the utility and basis of universal standards, favoring instead practices that are localized and contextualized in time.

Within this international and multi-cultural context is the increasing requirement from funding agencies and publishers that data be shared as openly as possible (ERC Scientific Council, 2022; Jones, 2012), and the demonstrated societal benefits that derive from open science (Cole et al., 2024; Klebel et al., 2025; Tsipouri et al., 2025; Van Vaerenbergh et al., 2026). Within biological anthropology, researchers have been lauded for compiling, organizing, and facilitating free access to their skeletal data. For example, hundreds of research papers have been published using W.W. Howell’s dataset of over 60 measurements from the skulls of 50 males and 50 females from 28 populations from around the world that was posted on the internet in 1996 (Howells, 1996; accessible through this link: https://web.utk.edu/∼auerbach/HOWL.htm), a dataset credited as instrumental in demonstrating the lack of distinct races in our species (McHenry and Delson, 2008). However, datasets like this are starting to come under question as anthropologists reconsider what community-consent looks like for retrospective research with data that do not fall under specific laws and institutional regulations. As of now, there are no universally-accepted guidelines that specifically address research with legacy skeletal data.

As the DeMoDa SAB considered how to manage and share dental morphological legacy data so that the research questions can diversify, we guided our decision-making with principles derived from a survey of the legal context of research on human remains, and the ethical guidance provided for the repatriation of Indigenous remains, human remains more generally, health databases and biobanks, and the study of Indigenous and ancient DNA. An overview of the stakeholders and influences on our decision-making are illustrated schematically in **Figure 2**. We summarize our deliberations in the sections that follow.

**Figure 2.**
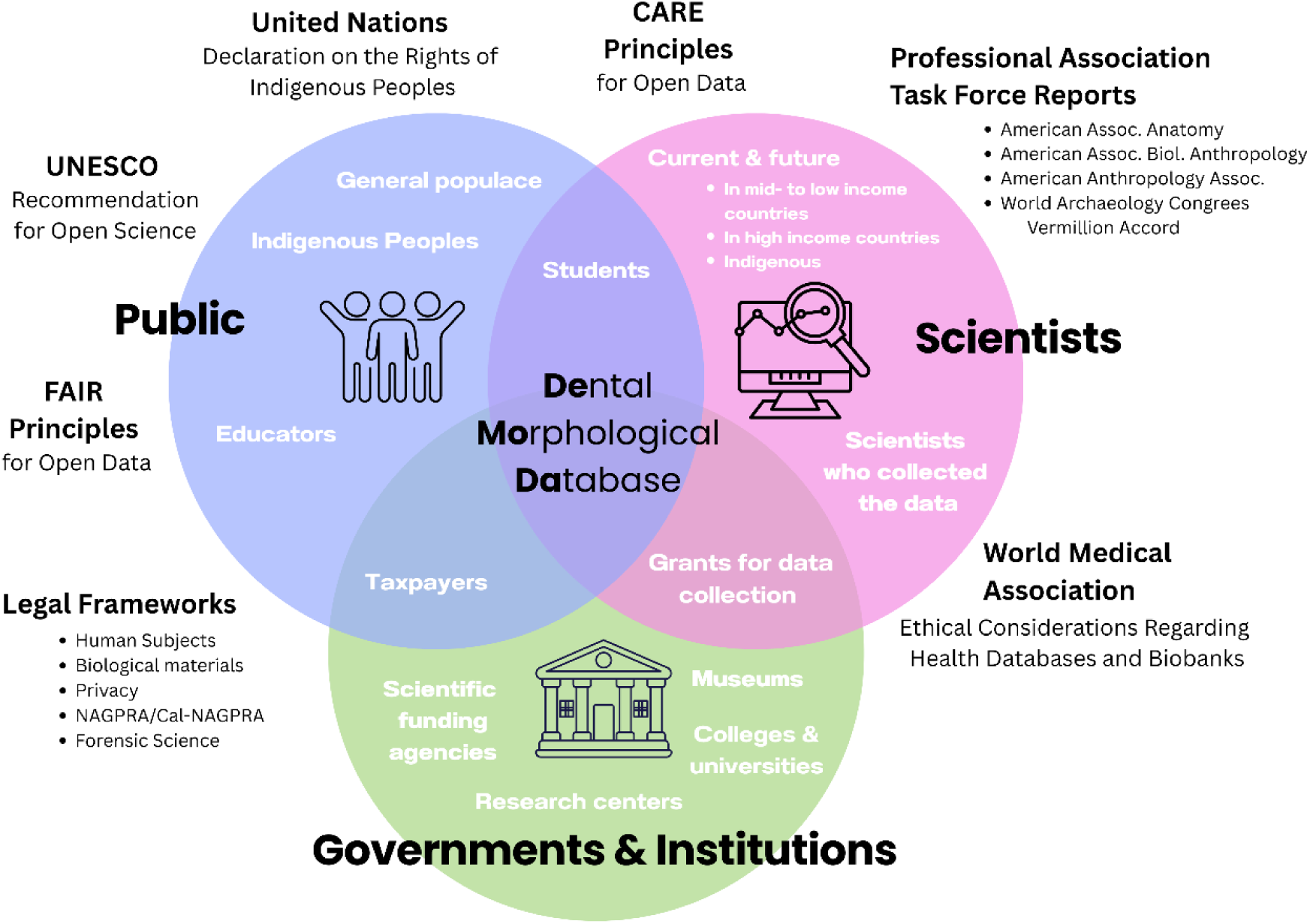
Schematic overview of the stakeholders (Public, Scientists, Governments & Institutions) and the legal frameworks, guiding principles, declarations, and task force reports that were considered in the design of DeMoDa.

## VII. Are there applicable legal frameworks for dental anthropological data?

The data in DeMoDa are based on observations of human skeletal material. Variations in tooth shape were assessed visually and assigned numerical scores based on comparison with published standards (Hanihara, 2008; Scott and Irish, 2017; Turner et al., 1991). All data incorporated into DeMoDa are anonymous and were obtained in accordance with the applicable governmental laws and institutional policies and procedures at the time. Given the nature of these data, they do not fall under legal frameworks for human subjects, privacy, or biological materials. To provide the reader with a clear understanding as to why, we briefly summarize these legal contexts below.

“Human subjects” rules do not apply to skeletal remains according to the Common Rule (45 CFR 46) in the United States, as a skeleton is not a living individual. Although we cannot be certain as to whether or not any of the dental remains scored in DeMoDa were replicas, Institutional Review Boards that consider research on human subjects do not consider anonymized casts as “human subjects” (Capili and Anastasi, 2024).

Privacy laws are not applicable because these data were collected from people who have been dead for centuries or even millennia and are not linked to a known, specific individual. The data in DeMoDa are from the study of institutional/museum skeletal collections, based on dental remains of people identified only by a museum identification code. In the European Union, the General Data Protection Regulation does not apply to deceased individuals, even if they are identifiable (Regulation (EU) 2016/679 of the European Parliament and of the Council of 27 April 2016 on the protection of natural persons with regard to the processing of personal data and on the free movement of such data [2016] OJ L119/1) (https://eur-lex.europa.eu/eli/reg/2016/679/oj, last accessed 20 April 2026). And in the United States, the health privacy law HIPAA protects a deceased person’s health records only for 50 years after death (https://www.hhs.gov/hipaa/for-professionals/privacy/laws-regulations/index.html, last accessed 20 April 2026).

Dental morphological data do not fall under the legal definition of “biological material,” a term that refers to materials derived from living organisms that have the ability to reproduce because they contain DNA or RNA (Petrini, 2012). Consequently, the data in DeMoDa do not fall under governmental legal frameworks for biological materials collected for biomedical research and held in biobanks, such as the Council of Europe Recommendation CM/Rec(2016)6 (https://www.coe.int/en/web/human-rights-and-biomedicine/biobanks, last accessed 20 April 2026).

In some countries, such as Spain, archaeological or historical objects discovered via excavation or chance are considered public property (public domain) and state domain (Spanish Historical Heritage Law 16/1985, https://www.boe.es/eli/es/l/1985/06/25/16/con, last accessed 28 May 2026). In contrast, at least two countries do have specific laws governing the human remains of Indigenous Peoples. For example, the Japanese government implemented “The Ainu Policy Promotion Act” in 2019 that explicitly recognizes the Ainu people as an Indigenous People, and set the guidelines for Ainu human remains and the associated burial goods to be repatriated to the local communities where they were excavated (https://www.cas.go.jp/jp/seisakukaigi/ainusuishin/policy.html#policy; https://www.cas.go.jp/jp/seisakukaigi/ainusuishin/return.html, last accessed 28 May 2026). Similarly, in the United States, remains of Indigenous groups are subject to the Native American Graves and Repatriation Act (NAGPRA; Native American Graves Protection and Repatriation Act, Pub. L. No. 101-601, 104 Stat. 3048 (1990); (Nash and Colwell, 2020; Office of the Secretary, Interior (United States), 2023) and in California, the State of California’s NAGPRA law AB 275 (State of California’s NAGPRA law AB 275 (California Health and Safety Code §§ 8010–8030, 2024). These laws specifically refer to human remains and artifacts related to Indian Tribes and Native Hawaiian organizations in the United States (including biological samples; Bader et al., 2023). Additionally, the United States passed legislation in 2022 under the African American Burial Grounds Preservation Act to identify and protect historic African American cemeteries. However, this Act only provides funding for preservation and does not discuss human remains within those cemeteries (Barber and Brock, 2024). In conclusion, the Japanese, United States, nor the California laws make reference to non-genetic data collected from human remains (such as dental morphological scores). For more detailed review of country-specific legislation regarding human remains in archaeological contexts as of 2011, we refer the reader to Marquez-Grant and Fibiger (2011).

Although we find no guidance for the data in DeMoDa from these legal frameworks, we did find guidance from the various documents developed for the implementation of repatriation efforts that aim to provide Indigenous communities sovereignty over their Ancestors and items of cultural patrimony. We discuss this in the next section.

## VIII. Guidance on the repatriation of Indigenous human remains and artifacts

When the NAGPRA federal law in the United States went into effect in 1990, it motivated a huge shift in the relationship between curatorial institutions, researchers, and Indigenous communities that bears on how we think about legacy data (Nash and Colwell, 2020). The United Nations Declaration on the Rights of Indigenous People in 2007 further shaped repatriation efforts around the world (United Nations Declaration of the Rights of Indigenous Peoples 2007, https://www.un.org/development/desa/indigenouspeoples/wp-content/uploads/sites/19/2018/11/UNDRIP_E_web.pdf last accessed 5 May 2026). Numerous other countries have followed in spirit and guidance on the repatriation of Indigenous remains and cultural artifacts. As mentioned in the previous section, in Japan, building on “The Ainu Policy Promotion Act” of 2019, research ethics concerning Ainu human remains and cultural artefacts are currently being formulated under the auspices of domestic relevant academic societies and the Hokkaido Ainu Association, and are likely to address the handling of legacy data, although they have not yet been published. While there is no specific federal law in Canada guiding how the skeletal remains of Indigenous/First Peoples are treated and repatriated, these practices are guided by local laws and statements developed by various Assemblies, Commissions, Task Forces, and Associations (Hanna, 2003; University of Toronto Department of Anthropology, 2020, https://www.anthropology.utoronto.ca/resources/policies/repatriation-policy last accessed 24 April 2026); Canadian Museum of History 2026 (https://www.historymuseum.ca/indigenous-heritage/repatriation last accessed 24 April 2026 and https://www.historymuseum.ca/wp-content/uploads/2026/04/Repatriation-Policy-2026-EN.pdf last accessed 24 April 2026). Australia is similar in lacking a federal law, but the repatriation of Indigenous remains has also transformed collections management in that country (Pickering, 2020). In the edited volume by Fforde et al. (2020), numerous chapters discuss repatriation efforts around the world, ranging from former colonial powers in Europe, to Russia and the Pacific, missionaries, and within former colonies such as Australia and Chile, and of course, the United States.

Within the policies and guidelines for repatriation that we reviewed, the disposition of non-genetic data is not specifically addressed, although we found a few statements that are potentially relevant. For example, the National Museum of Australia’s repatriation handbook advocates the return of “associated notes and data” to the descendant community (Pickering, 2020: 13). The Canadian Museum of History’s Repatriation Policy from 2026 says that casts and other replicas of Indigenous ancestral remains are eligible for return, but makes no mention of data. The University of California’s 2021 policy on Native American Cultural Affiliation and Repatriation in response to the passage of Cal-NAGPRA addresses the disposition of items that are not human remains or cultural items, encouraging campuses to voluntarily deaccession them as part of the repatriation process (2021, https://nahc.ca.gov/wp-content/uploads/2021/12/Final-UC-NAGPRA-Policy.pdf, last accessed 24 April 2026) This has led to the recent deaccessioning of a range of non-NAGPRA defined items, including photographic negatives of petroglyphs in California held by the University of California Los Angeles’ Fowler Museum (National Park Service Federal Register 2026 (https://www.federalregister.gov/documents/2026/01/07/2026-00073/notice-of-intended-repatriation-fowler-museum-university-of-california-los-angeles-los-angeles-ca#p-14, last accessed 26 April 2026).

Aside from the National Museum of Australia’s recommendation to repatriate data, all other guidance on repatriation that we surveyed refers specifically to physical items. The focus on physical, tangible items brings up the contrary, the idea of “virtual repatriation,” a concept that is potentially applicable to DeMoDa. In response to the suggestion that a museum’s sharing with an Indigenous Tribe images and scans of cultural patrimony constitutes a form of repatriation, (Boast and Enote, 2013) note that,

> *“[T]he objects denoted by repatriation in law, in convention, and in practice are those autochthonal objects created by the source community or culture of origin…These digital objects and data originate both from the collecting institutions and from the academy. Therefore, though data sharing is taking place, there is no* restitution *or* repatriation*”* (2013: 109, emphasis in original).

For example, Boast and Enote describe how, for the Zuni, “the word *repatriation* finally became understood as the bodily, material, and actual return of a Zuni item or items back to Zuni land and people” (Boast and Enote, 2013: 110, italics original). This concept for what should be repatriated raises the question of what dental morphological data are – something created by members of the academic community and/or something that inherently belongs to the individuals from whom these trait scores were observed? We do not have a specific answer to this question, but we hold it in mind.

In the context of repatriation, it is evident that the data in DeMoDa do not represent skeletal remains or replicas/images of skeletal remains, biological materials, tangible items created by the Indigenous group, nor do they carry personally identifiable data. Rather, the data in DeMoDa are already published in a wide range of presentations and can be found in digital format online and/or published in books in libraries around the world (e.g., Qaq et al., 2026). Consequently, these data are not something that can be “returned” in the same way that human remains or an object or even “notes” can be, even though the collection of these data caused harm to some descendent communities similar to the collection of the remains and objects that are clearly subject to repatriation laws.

## IX. Guidance for the ethical treatment of human remains within archaeology, anatomy, and biological anthropology

The World Archaeology Congress’ Vermillion Accord (1989; The Vermillion Accord on Human Remains, https://worldarchaeologicalcongress.com/code-of-ethics/ last accessed 5 May 2026) marked the first time professional archaeologists and Indigenous groups collaborated to develop a set of principles to guide practices around the human remains recovered through archaeological work (Fforde, 2020). The six principles of the Vermillion Accord hold respect as the focal point, calling for a balance between both the societal concerns and the scientific value. When the United States passed NAGPRA in 1990 and updated regulations in 2023 (https://www.federalregister.gov/documents/2023/12/13/2023-27040/native-american-graves-protection-and-repatriation-act-systematic-processes-for-disposition-or last accessed 5 May 2026), and Japan passed “The Ainu Policy Promotion Act” in 2019, this balance was made explicit in legal form for Indigenous human remains and cultural artifacts in those countries.

As academic disciplines began to consider the implications of non-consent for the remains of other unknown (and some known) people in institutional anatomical collections more broadly, task forces formed to develop guidelines that would be relevant for the remains of people who are not included in the laws focused on Indigenous Peoples (Agarwal, 2024; Jackson and Auerbach, 2026; Passalacqua et al., 2025). The American Association for Anatomy developed three guiding principles for legacy anatomical collections (Cornwall et al., 2024), defined as “historical collections containing human tissue, including human remains where provenance is unknown or unclear or the informed consent of the individual has not been determined” (https://www.anatomy.org/ANATOMY/About-Us/Policies/Legacy-Collections-Policy.aspx retrieved 27 February 2026). These guiding principles require that the custodian be aware of and adhere to the relevant and applicable legal frameworks, convey respect and acknowledge the manifest wishes of the deceased, and lastly, be aware of potential harm to the legacy of the deceased or others as a result of the custodian’s actions. Forensic anthropologists also developed guidelines and recommendations (Pilloud et al., 2025), leading to suggestions for standards by the Forensic Anthropology subcommittee of the Organization of Scientific Area Committees for Forensic Science, National Institute of Standards and Technology in the United States (Passalacqua et al., 2025).

Most recently, the American Association of Biological Anthropologists published a report from the Presidential Task Force on the Ethical Treatment of African American Human Remains (Auerbach et al., 2026; Jackson and Auerbach, 2026). This is the first analysis that paired perspectives from two communities: the priorities and concerns of both the descendant community and the community of biological anthropologists. Through surveys of those self-identified as part of an African American descendant community, concerns raised included the privacy of genomic and other biological data, the selling of data to “big pharma”, and the need for researchers to follow the laws on data protection and privacy. Given the atrocities inflicted on the African American community by governmental and healthcare institutions, these are not trivial points of concern (Washington, 2006). The Task Force advises that no research on African American remains occur without the explicit consent of relevant descendant communities(Auerbach et al., 2026).

Although these task force reports provide no guidance on the management of published skeletal data, they are relevant to DeMoDa in that they advocate that people from all communities be treated with the same high level of respect and consideration, be they Indigenous, former enslaved, unclaimed burials, impoverished, war dead, etc. We especially note the guidance that researchers be aware of potential harm to the legacy of the deceased or to other people close to them.

## X. Guidance on the ethical considerations for health databases and biobanks

Although the data in DeMoDa are not biological materials nor human remains, they are data about biological phenomena. Consequently, we also considered the ethical discussions and policies around health databases and biobanks. A survey of 117 open-access resources with DNA, RNA, protein, and phenotype data was recently published, providing a useful entry-point to these issues (Aguado and Hernández, 2026). A critical determinant in weighing the risks and potential benefits of human-derived data in biobanks is the degree to which the data can be linked to its specific source (Godard et al., 2003). The World Medical Association’s (WMA) Ethical Considerations Regarding Health Databases and Biobanks (Aicardi et al., 2016),

> “*lays down ethical principles for medical research involving human subjects, including the importance of protecting the dignity, autonomy, privacy and confidentiality of research subjects, and obtaining informed consent for using identifiable human biological material and data*” (adopted by the 53rd WMA General Assembly in 2002 and revised by the 67th WMA General Assembly in 2016; https://www.wma.net/policies-post/wma-declaration-of-taipei-on-ethical-considerations-regarding-health-databases-and-biobanks/, last accessed 24 April 2026).

The main focus of the WMA’s Declaration for health databases and biobanks revolves around privacy, protecting the identity and respecting the wishes of living individuals. This builds on the landmark WMA 1964 Declaration of Helsinki providing ethical principles for medical research involving human participants.

The data in DeMoDa, as already described, do not contain information about specific, known individuals. However, the primary DeMoDa data do make reference to specific cultural groups. There are numerous examples of laws and guidelines calling for specific protections for the genetic data from vulnerable populations, which has led to the concept of “group consent” in combination with, or in lieu of individual consent (Godard et al., 2003). As the anthropological community begins to turn to descendant communities for a type of group-consent for the study of archaeological populations, we found the Icelandic Healthcare Database and proposed deCODE genetic database an interesting comparison (Gulcher and Stefánsson, 2000).

The Icelandic Healthcare Database (IHD) is part of the nationalized healthcare system that holds excellent medical records for all of its citizens since World War I (Annas, 2000; Gulcher and Stefánsson, 2000). In the late 1990s, a genetic database called deCODE was proposed to link with the IHD in order to facilitate the identification of genetic risks for disease, and to do so in collaboration with a for-profit company from the United States (Árnason and Andersen, 2013). Aspects of the deCODE project were criticized heavily, including by the Icelandic Medical Association. Two of the criticisms that caught the attention of the DeMoDa SAB were that: 1) deCODE was to be established through “community consent”, meaning that the larger community would give approval for the endeavor and that an individual opposed to it would have to opt-out, and 2) the biobank was established in collaboration with a United States private company to use these DNA data for commercial purposes. As one author pointed out at the time, “it seems unlikely that the citizens of this welfare state would willingly make a large gift of their DNA to a for-profit U.S. corporation” (Annas, 2000: 1831). Annas further stated the view that “research should not be conducted on a population, even research related to migration patterns or the evolution of a genome, unless the benefit to the population is likely to outweigh the risks” (2000: 1831), with citation to the Committee on Genome Diversity (National Research Council (US), 1997) and Glantz et al. (1998). The development of the community-consent genetic database was ultimately found unconstitutional because genetic data, by definition, can never be fully anonymized (Árnason and Andersen, 2013). The for-profit corporation did go on to build a similar genetic database through individual consent and eventually sold it to another company for $415 million. The Icelandic people never received any financial compensation for sharing their genomic data (Árnason and Andersen, 2013).

For DeMoDa, there are no data (genetic or otherwise) linked to specific, known individuals, nor could they be linked to related individuals through identity-by-descent (as can be done with genome sequences). But, there are important take-aways for DeMoDa from our consideration of genomic databases and biobanks. Following the guidance of the WMA, the SAB holds that DeMoDa’s activities should benefit society, including the facilitation of public health objectives, and in so doing, not contribute to for-profit endeavors. Considering, and expanding on the concept of vulnerable communities, DeMoDa should provide additional anonymity (going above the individual level) to prohibit research on a specific population/cultural group to minimize potential harm to that group.

## XI. Critiques of research using Indigenous Peoples’ DNA

Although we have already clarified that dental morphological data are not genetic data, it is useful to consider how the use of genomic data has caused harm to some populations. The harm that research on human genetic data can inflict is not uniform across all populations, as historical and contemporary contexts dramatically shape the implications. The world’s Indigenous peoples have borne comparatively greater harm from some uses of these data than have other communities.

The concept of who is Indigenous can be academic (Corntassel, 2003), but from a practical perspective we borrow from the United Nations State of the World’s Indigenous Peoples (2009) (2009):

> *“Indigenous communities, peoples and nations are those which, having a historical continuity with pre-invasion and pre-colonial societies that developed on their territories, consider themselves distinct from other sectors of the societies now prevailing on those territories, or parts of them. They form at present non-dominant sectors of society and are determined to preserve, develop and transmit to future generations their ancestral territories, and their ethnic identity, as the basis of their continued existence as peoples, in accordance with their own cultural patterns, social institutions and legal system”* (2009: 4).

We referred previously to the 2007 United Nations Declaration on the Rights of Indigenous Peoples (UNDRIP) in reference to the treatment of human remains. This Declaration was a milestone on many levels, as it set in motion on-going formal efforts within the UN to advocate for and monitor Indigenous people’s rights and wellbeing. Among these rights, Article 31 of the Declaration states that,

> “*Indigenous peoples have the right to maintain, control, protect and develop their cultural heritage… including human and genetic resources… [and to] maintain, control, protect and develop their intellectual property of such cultural heritage*” (Department of Economic and Social Affairs, United Nations Secretariat, 2009): 67).

As genetic and genomic studies became possible and grew at an exponential rate over the last thirty years, there has been an interest in the DNA of Indigenous populations. From the biomedical perspective, communities seen as more isolated than others are more powerful for the genetic analysis of complex traits (Arcos-Burgos and Muenke, 2002). While not all of these isolated populations are Indigenous (Aguado and Hernández, 2026; Boycott et al., 2008; Capocasa et al., 2013), quite a few of them are (Fumagalli et al., 2015; Pontzer et al., 2018). From an evolutionary perspective, the genetic data from Indigenous populations provide deeper insight to our understanding of the breadth, depth, and structure of human genetic variation (Mulligan et al., 2004; Tishkoff and Verrelli, 2003). The relevance to human genetic variation to understanding and improving health has long been appreciated (Cavalli-Sforza, 2005; Templeton, 1999), and there is also a societal benefit as analyses of global genetic variation can help to break down misconceptions of race (Benn Torres, 2020).

The re-purposing/reuse of DNA sequence data from Indigenous peoples has raised a plethora of ethical issues about consent, control, and authority. A recent review article provides a helpful introduction (Halmai et al., 2025). Building on the UNDRIP Declaration from 2007, the movement for Indigenous Data Sovereignty gained momentum over the subsequent decades. More than a dozen resources now provide guidance on the use and reuse of Indigenous genomic data and biospecimens (e.g., Claw et al., 2018; Hudson et al., 2020). A framework developed by the Research Data Alliance’s International Indigenous Data Sovereignty Interest Group and the Global Indigenous Data Alliance is called the CARE Principles for Indigenous Data Governance (Carroll et al., 2020). The four CARE Principles are envisioned to work alongside the four FAIR Principles of open data by drawing attention to the people from which data were collected and the purpose for which the data are being analyzed (Carroll et al., 2021).

The CARE Principles are seen as the minimum that researchers should employ when contemplating the reuse of data derived from Indigenous Peoples (Carroll et al., 2021, 2020). The first principle is to consider the **collective benefit** for Indigenous Peoples, ensuring that data are being reused for inclusive development and innovation, for improved government and citizen engagement, and for equitable outcomes. Second, researchers recognize the rights and interests of Indigenous Peoples’ **authority** to control and govern these data, and be actively involved in stewardship decisions for Indigenous data held by other entities. Third, researchers have the **responsibility** to nurture respectful relationships with Indigenous Peoples from whom data originate. And fourth, researchers have an **ethical obligation** to ensure that Indigenous Peoples’ rights and wellbeing are centered in the planning for how data are reused so as to minimize harm, maximize benefits, promote justice, and allow for future use. Central to this last point is that Indigenous Peoples need to be the ones assessing benefits, harm, and the potential future uses. Today, best practice for the collection and analysis of data from an Indigenous community is to consult with that community prior to beginning the project, and to include the Indigenous community perspectives and priorities in the research design and implementation through a process called community consultation (e.g., (Claw et al., 2018; Dadich et al., 2019; Fitzpatrick et al., 2016).

## XII. Our decision not to pursue community consultation at this time

The data in DeMoDa were collected to probe research questions at a much broader scale of human variation compared to the research questions derived from today’s ground-up consultation with descendant communities (e.g., (Blakey, 2010; Clinton and Jackson, 2021; Lindo et al., 2018). Due to the broad scope of the research question, the researchers who collected the DeMoDa data typically did so from across multiple small populations and combined them to form larger populations. Along the way, in the process of concatenation, information on specific institutional repositories in which the human remains were held, the numbers of individuals from specific subgroups, and individual museum or other institutional identification numbers or codes were often not retained or were noted only cryptically. However, we have evidence that well over 500 populations/groups/communities are affiliated in one way or another with these data (**Appendix 2**).

There is not a formal list of Indigenous Peoples, as the UN State of the World’s Indigenous Peoples (2009) determined that this identity should be flexible and self-described rather than assigned by an outside entity. Therefore, following on the guidance from the professional society task forces described in Section IX, we decided to apply the same respect for all of the communities and populations represented in these data in accordance with the CARE Principles, be those communities considered Indigenous or not.

Following the guidance that any research on a specific community should be done in consultation with that community (e.g., Auerbach et al., 2026; Carroll et al., 2021), the SAB decided that the data in DeMoDa should be made public without any of the information that would enable community-specific research. This higher-level of de-identification (removing community information) significantly reduces the potential harm to any of the included communities, while also enabling the societal benefit that comes from the ability to analyze individual-level data at a much broader scale. Future iterations of DeMoDa could be designed to accommodate and balance access to subsets of the data in different ways. The SAB welcomes those discussions and collaborations.

## XIII. Considerations on the reuse of legacy data from skeletal remains

The study of human variation is historically rooted in efforts to identify similarities and differences between human populations, sometimes with the intent of using those differences to justify mistreatment built into colonial practices, often referred to as scientific racism (Comas, 1961). Although the academic discipline has largely shifted to more diverse and biologically grounded motivations for studying human variation (Fuentes, 2010; Washburn, 1951), the legacy of harm is pervasive. For some communities, the wounds of scientific racism remain quite raw. As anthropologists continue to explore how our discipline can help to redress the academy’s contributions to this history, legacy data from skeletal remains have become a point of conversation, and for some, a point of contention.

In a report from the American Anthropological Association’s Commission for the Ethical Treatment of Human Remains, the authors describe a listening session held at the annual meeting of the American Association of Biological Anthropology in 2024 that touched on the reuse of legacy data (American Anthropological Association, 2024). During this session, the suggestion was raised that legacy data only be reused if contemporary standards of consent for the study of human remains have been met (2024: 26). The authors of the report claim that,

> “*Nearly all major anthropological and biological anthropology journals no longer accept publication of ‘legacy data’ and have policies in place that ask for contemporary standards of consent*” (2024: 26).

We are unable to find examples of this shift in publication policy. The *American Anthropologist* does not have a section on data availability or data ethics on their Submitting to the Journal webpage (https://www.americananthropologist.org/how-to-submit/, last accessed 24 April 2026). The *American Journal of Biological Anthropology’s* Author Guidelines require authors to include a data availability statement, and for studies reporting on data from human remains, a statement describing community consultation and other permissions (https://onlinelibrary-wiley-com/page/journal/26927691/homepage/forauthors.html, last accessed 24 April 2026). However, the Author Guidelines are explicit that “This requirement refers specifically to new data; legacy data is not included.” The *Journal of Human Evolution*’s Guide for Authors (https://www.sciencedirect.com/journal/journal-of-human-evolution/publish/guide-for-authors, last accessed 24 April 2026) points to Elsevier’s general guidelines on human research (https://www.elsevier.com/about/policies-and-standards/research-ethics#0-human-research, last accessed 24 April 2026), a statement that leans heavily on the WMA’s principles discussed previously. There is no specific mention of legacy skeletal data or community consultation. Consequently, we conclude that there is presently no wide-spread evidence that academic journals will not accept the reuse of published data from skeletal populations without new consent from modern descendant communities. However, this claim by the American Anthropological Association’s Commission indicates that some anthropologists are pushing the discipline in that direction.

Through our consideration of the laws and the range of guiding principles presented in the preceding sections, we have come to appreciate that the new standards of practice for human remains are an imperfect fit for legacy dental anthropological data (and non-biological skeletal data more generally). Human remains are objects that require handling and disposition. Published data are intangible, and permanently and pervasively present in libraries and on the internet. We have concluded with the available guidance at the moment that the best way forward is to consider how we can shape the future use of these data so as to encourage the societal benefit they offer while minimizing the potential harm they could facilitate to specific communities.

Our conclusion is in line with the discussions unfolding within biological anthropology. In 2025, the American Association of Biological Anthropologists’ hosted the symposium *An Open Debate on the Analysis of Legacy Data from Human Skeletal Remains* at their annual conference organized by the AABA Chair of the Science Policy Committee, Lumila Menéndez. This symposium facilitated a broad discussion that explored positives, negatives, and ways to pursue the former while avoiding the latter (Beguelin and Menéndez, 2025; Edgar and Berry, 2025; Klafehn, et al., 2025; L’Abbe et al., 2025; Quinto Sanchez, 2025; Rivera et al., 2025; Seguchi and Hayashi, 2025; Wilson, 2025). For example, Beguelin & Menéndez described how the craniometric dataset created by Héctor Pucciarelli, while problematic, facilitates a considerable amount of science for researchers with limited access to funding (Beguelin and Menéndez, 2025; Menéndez and Urban, 2025; Paschetta et al., 2017; Pucciarelli et al., 2006). L’Abbe et al. (2025) from the University of Pretoria, South Africa, proposed that the osteometric and morphological data from 20th century cadaveric skeletons from the Apartheid era be available to researchers, but that approval from an oversight committee be granted before the data are employed. Wilson (2025) noted that original datasheets and field notes also need to be curated, preserved, and made available alongside legacy data, as these contain important information that may well have scientific value for decades. Seguchi & Hayashi (2025) describe the important role that historical data play in characterizing ancestral populations so that Japanese and Filipino war dead from World War II can be properly distinguished and repatriated to their descendants. Klafehn et al. (2025) noted that the use of existing legacy datasets documenting human variation could reduce the pressure to excavate more archaeological sites, assisting with preservation of sites and the human remains enterred within them. Alternatively, Klafehn et al. (2025) suggest that newly-excavated sites could enable legacy datasets to be replaced through more modern frameworks of consultation and descendant community consent. This symposium also raised points of concern from the next-of-kin to the individuals who are represented in a large database of CT scans. Edgar & Berry (2025) conducted interviews with next-of-kin focus groups for the New Mexico Decedent Image Database, finding that while there was strong support for the use of these data for forensic investigations and health research, there were negative reactions to artistic uses. This echoes the point made by forensic anthropologists – data derived from human remains are not harmful in and of themselves, the harm comes from the stories that may be told with them (Borgelt et al., 2025b).

A key component of the scientific process is to have the ability to revisit data and the conclusions drawn from their analysis. Re-analysis of legacy data can enable those stories to be rewritten when flaws are found in the previous research. The most well-known example of how access to, and reanalysis of legacy data helped to redress scientific racism is Gould’s critique of several disciplines within American race science (Gould, 1981, 1978). Particularly relevant to this discussion of legacy skeletal data, he revisited one of the more notorious examples of typological and racist investigations on human cranial variation, Samuel G. Morton’s study of head size (Morton, 1844, 1839). Because Morton published his data and described in detail his methods, Gould was able to replicate Morton’s research step-by-step. In so doing, Gould concluded that Morton’s methods introduced a bias that likely resulted from subconscious racist expectations (Gould, 1981). Perhaps not surprisingly, as is the way science is supposed to unfold, Gould’s critique itself has also been reassessed (Lewis et al., 2011). This episode in the history of science has been taught in many, many anthropology and biology courses over the years, helping students understand that science is not apolitical.

This well-known example is a useful demonstration of the scientific process at work: a scientist facilitates access to their original data and carefully describes their methods so that other scientists can challenge the data, the methodologies, and/or the conclusions. Replicability is an essential part of best-practice in science. If the original data from past studies are no longer considered appropriate for re-analysis or new analysis, the growth of the study of human skeletal variation to move beyond typology is halted. This lack of progression should be concerning to anthropologists as the potential for science to benefit society may be lost.

## XIV. Making these legacy data open-access through a curated dataset

With all of these various considerations in place, we developed a list of the best practices for sharing and reusing legacy skeletal/dental data (non-biological, non-genetic), and how DeMoDa addresses them in order to advance the mission of open science (**Table 1)**.

**Table 1.** Best-Practices for the Sharing and Reuse of Legacy Skeletal Data.

| Guideline/Recommendation |  | The Responsive Design Decisions for DeMoDa |
| --- | --- | --- |
| **FAIR Principles** | Findability | DeMoDa Dataset v1 is published on a public repository so that it is findable through an internet search and via DOI. |
|  | Accessibility | DeMoDa Dataset v1 is published in an established repository (Dryad) that ensures integrity of the data and metadata. |
|  | Interoperability | Extensive metadata are provided with DeMoDa Dataset v1, facilitating interoperability. |
|  | Reuse | DeMoDa Dataset v1 is published with CCO copyright permission, so data can be reused freely. |
| **CARE Principles** | Collective benefit for Indigenous Peoples | Structure of the published DeMoDa Dataset v1 facilitates biological research, large climatological/geographic patterns, and reassessments of the pattern of global variation. This research benefits Indigenous Peoples through the collective health insights that may derive from improved understanding of human biology, and the social justice provided by reassessments of variation that undermine racist concepts. |
|  | Authority: Researchers recognize the rights of Indigenous Peoples to control and govern data. | DeMoDa Dataset v1 protects the rights of Indigenous Peoples to control their data by providing an attractive, scientifically flexible configuration of the data (conferring a benefit) that lacks the contextual information needed to investigate population-specific questions (minimizing harm, as individuals are pooled into large geographic regions). For the future, the main database of DeMoDa has the design structure to enable population-specific access settings according to Indigenous communities' decisions. (Although most of the data are not associated with much detail.) |
|  | Responsibility to nurture respectful relationships with Indigenous Peoples from whom the data originate. | The published DeMoDa Dataset v1 does not include identifying information that could be used to investigate questions about specific Indigenous (or other) communities. The metadata associated with DeMoDa Dataset v1 directs researchers interested in those types of questions to pursue them in collaboration with the Indigenous community(ies). |
|  | Ethics: Indigenous Peoples' rights and wellbeing are centered in the plan for data reuse to minimize harm, maximize benefits, promote justice, and allow future use. | The DeMoDa Scientific Advisory Board is not providing full access to the original data in order to protect the rights and wellbeing of Indigenous Peoples. This same level of protection is applied to all of the more than 500 populations included in DeMoDa Dataset v1 (as per the advice from the American Association for Anatomy and the American Association of Biological Anthropologists). The names of cultural groups included in the 32 geographic regions are listed in Appendix 2 so that communities can reach out with concerns and requests for future dataset releases. |
| **World Medical Association** | Data should benefit society, including public health objectives. | DeMoDa Dataset v1 presents the data in original form so that researchers have access to the more descriptive scoring system and individual-level data. This will enable more biologically-oriented questions that require covariance analytical methods. |
|  | Data should be open access. | DeMoDa Dataset v1 is published open access with no copyright restrictions. |
| **Forensic Anthropology** | Encourage researchers to not use the data to study questions that harm marginalized communities. | DeMoDa Dataset v1 does not contain the level of detail about populations that would enable these types of research questions. We also provide guidance on how to determine if a research question should be pursued through community consultation. |

Recognizing that once data are released to the public, there is no way to pull them back, the DeMoDa SAB decided to release a curated dataset rather than enable full access to the main database: DeMoDa Dataset version 1. The aim of the curated dataset is to facilitate the scientific investigation of human species-level variation without the detailed contextual information that would enable research on narrowly focused questions about the migration, origins, and/or gene flow of specific cultural groups. The general consensus among the professional community of biological anthropologists today is that these latter, more specific questions are more appropriately explored in consultation with the communities that would be directly impacted by the results. We also chose not to classify populations by scientifically out-dated major racial/ethnic categories but rather geographic regions, as we prefer to encourage new approaches to human biological variation, as has been repeatedly called for by previous scholars (e.g., (Clever and Jong, 2025; Echo-Hawk, 2010; Graves, 2021).

Two specific steps were taken to desensitize the contextual information for the open/FAIR/CARE publication of the data through DeMoDa Dataset version 1. Rather than using population names or detailed geographical locations, we organized the published dataset into large geographical regions so that no single population can be identified. We also removed all other information (when provided by the data-collector), as this might enable specific museums or other repositories to be identified, and thereby the more detailed source population(s) revealed.

## XV. DeMoDa Dataset Version 1

Each individual in the dataset has three pieces of contextual information. The first is **a unique DeMoDa identification number** that carries no information on the repository museum or institution but enables us to link it to the main database in case a question or concern arises. When skeletal biological **sex** was indicated by the researcher, we also include this. And third, we indicate the geographic region of the individual. This curated dataset is organized into **32 geographic regions (Figure 3)**. In an effort towards transparency, we provide a list of the populations included within each of these 32 large geographic regions for descendant communities and communities of care to evaluate (**Appendix 2**). The list includes a mix of cultural names and geographic locations, as we often worked from cryptic references preserved in original data collection notes. There were numerous instances when it was unclear if the geographic location listed on a data sheet was meant to clarify a population name or to indicate a different group. Therefore, we list everything.

**Figure 3.**
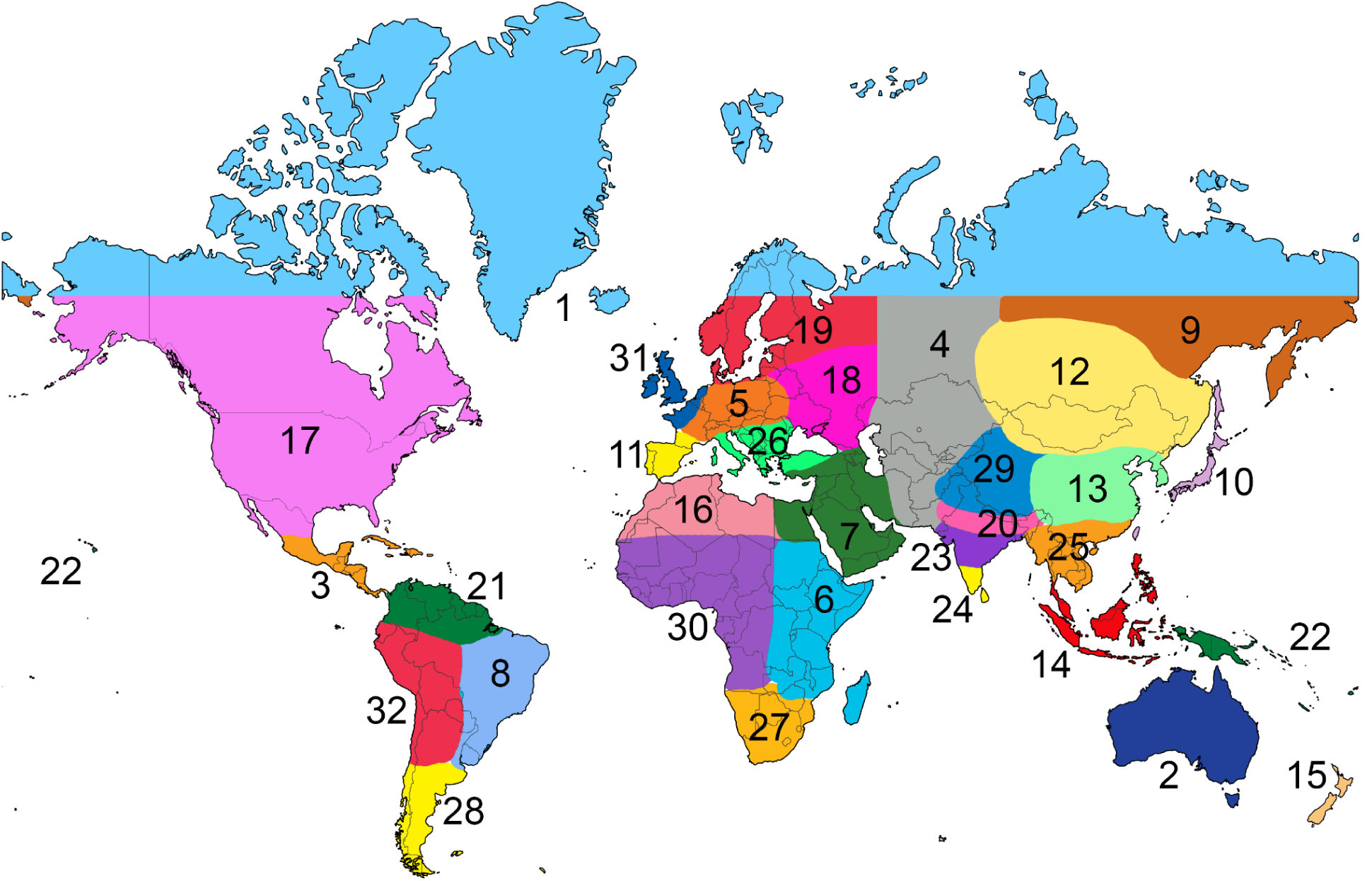
Populations in DeMoDa Dataset v1 were categorized into the 32 major geographic regions approximately illustrated on this world map: 1-Arctic (N66° and higher), 2-Australia, 3-Central America, 4-Central Asia, 5-Central Europe, 6-Eastern Africa, 7-Eastern Mediterranean/Levant, 8-Eastern South America, 9-Eastern Siberia, 10-First Island Chain, 11-Iberian Peninsula, 12-Inner Asia, 13- Mainland East Asia, 14- Malay Archipelago, 15-New Zealand, 16-North Africa, 17-North America, 18-North of Black Sea, 19-Northern and Northeastern Europe, 20-Northern India, 21-Northern South America, 22-Pacific Islands, 23-Peninsular India, 24-South India, 25-Southeast Asia Mainland, 26-Southeastern Europe, 27-Southern Africa, 28- Southern South America, 29-Tibetan Plateau and Siwalik Hills, 30-Western Africa, 31-Western Europe, 32-Western South America. Basemap by Ultimaps.com.

In addition to these three contextual fields, the dataset contains scores for 246 dental traits for each individual (as applicable). These data are presented via the two scoring systems we developed to enable the combination of data from different researchers.

## XVI. The DeMoDa Scoring Systems

In order to maximize the societal and scientific value of dental anthropological data, we need to be able to combine data from different researchers and facilitate the “big data” questions that can yield new insight into human biology. As the discipline of dental anthropology was developing, several morphological scoring systems were developed and tweaked. Consequently, one of the major hurdles to the creation of this global dataset was to address the differences across scoring systems used to capture observed variation.

We focused on the scoring systems developed by Turner et al. (1991) with modifications in Scott & Irish (2017) and Hanihara (2008), as these researchers also collected the largest individual datasets. All three of these scoring systems characterize morphological variation in the adult human permanent dentition.

We developed two scoring systems that enable data from these three systems to be combined in a step-wise manner. DeMoDa Scoring System 1 collapses Turner et al. (1991) and Scott & Irish (2017). DeMoDa Scoring System 2 collapses Hanihara (2008) with DeMoDa Scoring System 1. The translation details between scoring systems are presented in **Appendix 3**. In total, these two datasets present world-wide dental variation for 33 morphological features, summarized in **Table 2**. Eight of these features are observed on individual maxillary and mandibular teeth, and most are observed on more than one tooth. As scores were made for the right and left sides of the dentition, the final dataset includes scores for 246 individual traits.

**Table 2.**
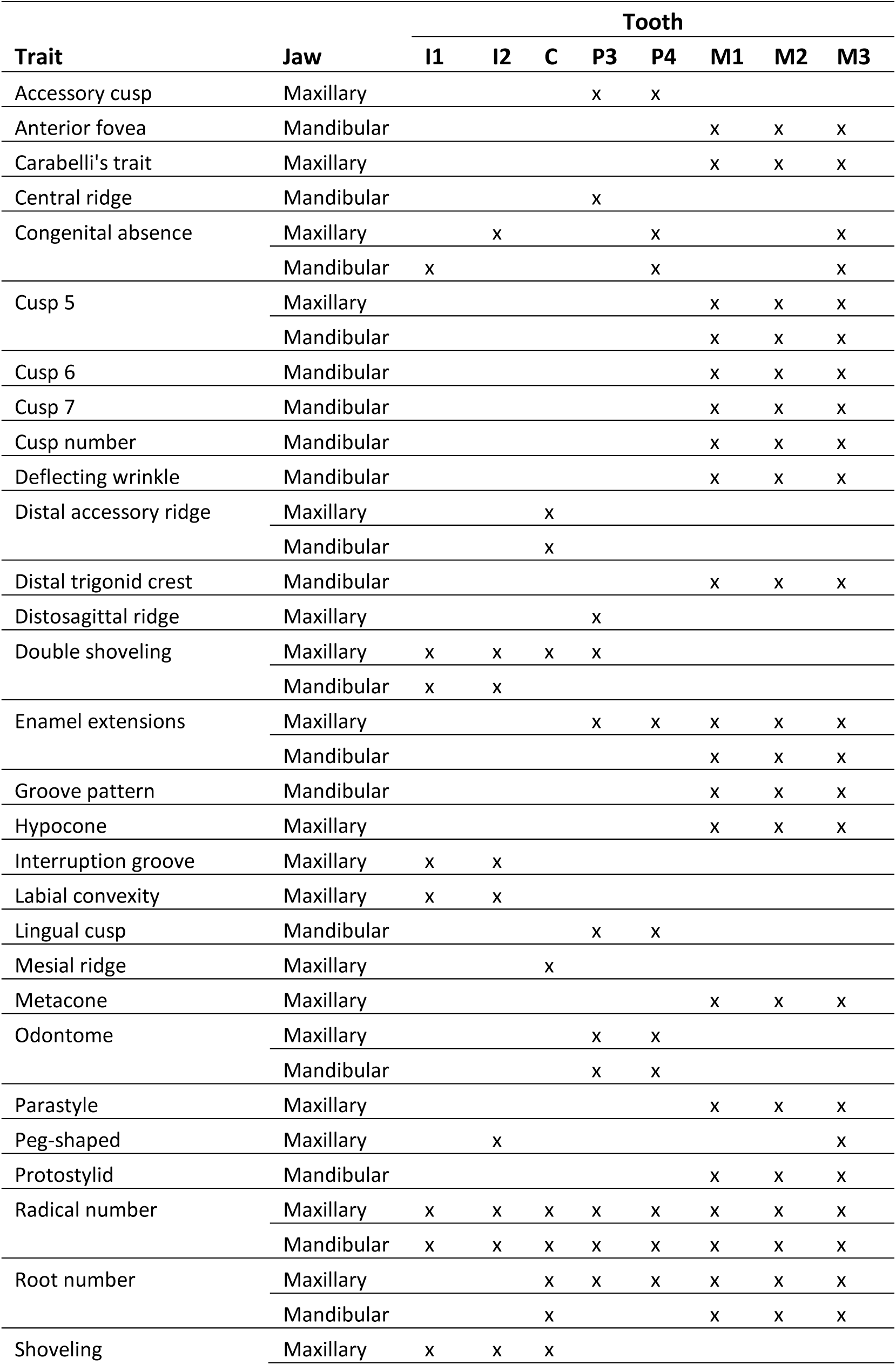
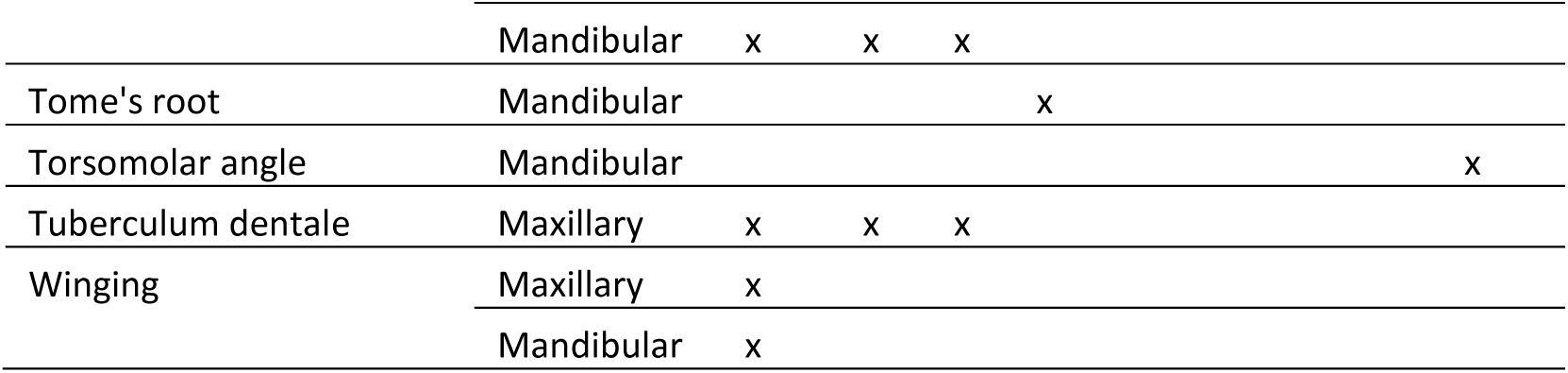
List of morphological traits in the DeMoDa Scoring System.

A key distinction between Turner et al. (1991) and Hanihara (2008) is in the level of detail captured by each scoring system. Turner et al. (1991) often used more expression categories for a trait compared to Hanihara (2008). Consequently, as the systems were combined, we converted the more-nuanced scoring system for a trait into the less-nuanced system for that same trait. This resulted in a loss of information, but at the same time, enabled a major increase in sample size and geographical representation. Some research questions will benefit more from the greater degree of specificity of trait expression, while other questions are better addressed with less specificity but a larger sample size. Therefore, we present two versions of DeMoDa Dataset v1. The first presents the combined data through DeMoDa Scoring System 1, with a total of 6,838 individuals. The second presents the data combined through DeMoDa Scoring System 2, with a total of 17,308 individuals.

## XVII. Guidance on the use of DeMoDa Dataset version 1

In the spirit of open access to data and the FAIR principles, DeMoDa Dataset verson 1 is designed so that it can be shared freely. We realize that we cannot control how the data are used nor the questions that are asked. However, we strongly encourage researchers to consider implementing an ethical framework for research that stems from these data and apply the CARE principles in tandem (Carroll et al., 2021, 2020). We advocate using these data to test hypotheses that help us understand human biological variation without racializing or marginalizing populations. Here we provide a series of questions that may help to guide hypothesis formation and interpretation of the results:

- Is the scientific answer to my research question going to provide more benefit to humanity than harm to any specific community?

○ *If yes, proceed*.
- Does this research approach the data from a typological perspective?

○ *If you answer yes to this question, consider re-framing your question/hypothesis*.
- Does this research have the potential to harm or specifically offend any of the descendant communities that may be represented in the analyses?

○ *If you answer yes to this question, consider re-framing your question/hypothesis*.
- Is there an opportunity to involve descendant communities in collaboration or consultation?

○ *If you answer yes to this question, consider taking the time and making the effort to make those contacts*.

## XVII. Accessing and citing DeMoDa Dataset version 1

As the DeMoDa Dataset version 1 is published on a public repository, it has been assigned a DOI and should be cited as:

Hlusko, L.J., Martínez de Pinillos González, M., Álvarez Fernández, A., Delgado Esteban, B., Delgado, M., Dern, L., Irish, J.D., Kondo, O., Martinón Torres, M., Scott, G.R., Thiebaut, A., Paul, K.S., Pilloud, M.A., Rathmann, H., Reyes-Centeno, H., Vlemincq-Mendieta, T., 2026. Global dataset of human morphological dental variation categorized into 32 geographic regions [Dataset].

*This dataset will not be released until this manuscript has been peer reviewed and accepted for publication*.

We also ask that users of the dataset also please reference the concept of the Dental Morphological Database and the reasoning that underlies the design of DeMoDa Dataset version 1 by citing this publication.

## XIX. Facilitating public engagement with human variation

In addition to the DeMoDa Dataset version 1 published on the open access repository Dryad, we also built an interactive web map that enables anyone with access to the website (https://demoda.cenieh.es) to explore for themselves how scores for 15 of the more well-known traits vary. The scores are shown in generalized geo-referenced pie charts with no population names. As the map is interactive and presents a small subset of the data, and because the data cannot be downloaded, the geographic locations in some instances are more detailed than what is published in DeMoDa Dataset version 1.

The data used for this public, interactive map feature overlap with the data that serve as the foundation for the forensic tool rASUDAS (https://osteomics.com/rASUDAS2/ (Scott et al., 2018a), but presents the data in a very different format. This combination provides a useful teaching tool for students learning about population histories and inter- and intra-populational variation (Paul et al., in prep).

## XX. Inviting conversation and critique of DeMoDa’s approach to managing these legacy Data

For the time being, the main database of DeMoDa will serve as a secure repository for the contextual data for each of the 17,308 individuals represented in DeMoDa Dataset v1. The SAB will continue to assess and discuss this project and the publication of the DeMoDa Dataset v1’s impact in both scientific and societal realms. We welcome and appreciate feedback on DeMoDa Dataset v1 and the thought processes that we used to decide on the design for this dataset and the larger DeMoDa project. The SAB’s future work will be adjusted as the scientific community, descendant communities, communities of care, and other stakeholders respond to this first step. While we fully anticipate that some colleagues will disagree with aspects of our logic and the structure of DemoDa, we hope that our approach sparks productive discussions on the reuse and management of non-biological legacy data from human skeletal remains.

## Acknowledgements

This research was funded by the European Research Council within the European Union’s Horizon Europe (ERC-2021-ADG, Tied2Teeth, project number 101054659). Views and opinions expressed are however those of the author(s) only and do not necessarily reflect those of the European Union or the European Research Council. Neither the European Union nor the granting authority can be held responsible for them.

The authors extend immense gratitude to the numerous colleagues who engaged in conversation about DeMoDa and helped to shape our perspectives, with specific mention of Lumila Ménendez and the American Association of Biological Anthropologists’ Science Policy Committee on Legacy Data from Human Skeletal Remains: Noriko Seguchi, Marien Beguelin, Healther Edgar, Erika L’Abbe, Mirsha Quinto Sánchez, Michael Rivera, and Teresa Wilson; and also Sheela Athreya, Ben Auerbach, Graciela Cabana, Rebecca Kormos, Nicholas Passalacqua, Arielle Pastore, Sabrina Sholts, Ian Towle, and Trudy Turner.

## Author contributions

**Marina Martínez de Pinillos González:** created the DeMoDa scoring systems; served as lead for the development of the database structure, data compilation and curation, and website; member of the DeMoDa Scientific Advisory Board; edited the manuscript

**Ana Álvarez Fernández:** assisted with development of the database structure, data compilation and curation, and website; provided Scientific Advisory Board support; edited the manuscript

**Beatriz Delgado Esteban:** assisted with development of the database structure, data compilation and curation, and website; provided Scientific Advisory Board support; edited the manuscript

**Miguel Delgado:** member of the DeMoDa Scientific Advisory Board; edited the manuscript

**Laresa L. Dern:** assisted with data compilation and curation; edited the manuscript

**Joel D. Irish:** member of the DeMoDa Scientific Advisory Board; assisted with data compilation and curation; edited the manuscript

**Osamo Kondo:** member of the DeMoDa Scientific Advisory Board; edited the manuscript

**María Martinón Torres:** member of the DeMoDa Scientific Advisory Board; edited the manuscript

**Mario Modesto-Mata:** co-lead for the development of the database structure; member of the DeMoDa Scientific Advisory Board; edited the manuscript

**G. Richard Scott:** co-lead on project concept; member of the DeMoDa Scientific Advisory Board; co-lead on the creation of the DeMoDa scoring systems; assisted with data compilation and curation; edited the manuscript

**Arthur Thiebaut:** provided Scientific Advisory Board support; edited the manuscript

**Kathleen Paul:** member of the DeMoDa Scientific Advisory Board; provided liaison with the Dental Anthropology Association; edited the manuscript

**Marin Pilloud:** co-lead on project concept; member of the DeMoDa Scientific Advisory Board; assisted with data compilation and curation; provided liaison with the Dental Anthropology Association; edited the manuscript

**Hannes Rathmann:** member of the DeMoDa Scientific Advisory Board; assisted with data compilation and curation; edited the manuscript

**Hugo Reyes-Centeno:** member of the DeMoDa Scientific Advisory Board; edited the manuscript

**Tatiana Vlemincq:** assisted with data compilation and curation; edited the manuscript

**Leslea J. Hlusko:** co-lead on project concept; lead for fundraising; lead on project management; Chair of the DeMoDa Scientific Advisory Board; assisted with development of the database structure, data compilation and curation, and website; wrote the manuscript

## Notes

### Competing Interest Statement

The authors have declared no competing interest.

### Summary of Updates

A few typos have been corrected and the link to the dataset revised.

## References

Adel, M., Hunt, K.J., Lau, D., Hartsfield, J.K., Reyes-Centeno, H., Beeman, C.S., Elshebiny, T., Sharab, L., 2025. Precision Assessment of Facial Asymmetry Using 3D Imaging and Artificial Intelligence. J. Clin. Med. 14, 7172. 10.3390/jcm14207172

Adhikari, K., Fontanil, T., Cal, S., Mendoza-Revilla, J., Fuentes-Guajardo, M., Chacón-Duque, J.-C., Al-Saadi, F., Johansson, J.A., Quinto-Sanchez, M., Acuña-Alonzo, V., Jaramillo, C., Arias, W., Barquera Lozano, R., Macín Pérez, G., Gómez-Valdés, J., Villamil-Ramírez, H., Hunemeier, T., Ramallo, V., Silva de Cerqueira, C.C., Hurtado, M., Villegas, V., Granja, V., Gallo, C., Poletti, G., Schuler-Faccini, L., Salzano, F.M., Bortolini, M.-C., Canizales-Quinteros, S., Rothhammer, F., Bedoya, G., Gonzalez-José, R., Headon, D., López-Otín, C., Tobin, D.J., Balding, D., Ruiz-Linares, A., 2016. A genome-wide association scan in admixed Latin Americans identifies loci influencing facial and scalp hair features. Nat. Commun. 7, 10815. 10.1038/ncomms10815

Agarwal, S.C., 2024. The bioethics of skeletal anatomy collections from India. Nat. Commun. 15, 1692. 10.1038/s41467-024-45738-6

Aguado, S., Hernández, C.L., 2026. Databases, Biobanks and International Consortia: Major Resources for Human Population Genomics and Biological Anthropology. Evol. Anthropol. Issues News Rev. 35, e70025. 10.1002/evan.70025

Aicardi, C., Del Savio, L., Dove, E.S., Lucivero, F., Tempini, N., Prainsack, B., 2016. Emerging ethical issues regarding digital health data. On the World Medical Association Draft Declaration on Ethical Considerations Regarding Health Databases and Biobanks. Croat. Med. J. 57, 207–213. 10.3325/cmj.2016.57.207

American Anthropological Association, 2024. Commission on the Ethical Treatment of Human Remains Issues Final Report.

Annas, G.J., 2000. Rules for Research on Human Genetic Variation — Lessons from Iceland. N. Engl. J. Med. 342, 1830–1833. 10.1056/NEJM200006153422412

Arcos-Burgos, M., Muenke, M., 2002. Genetics of population isolates. Clin. Genet. 61, 233–247. 10.1034/j.1399-0004.2002.610401.x

Árnason, E., Andersen, B., 2013. deCODE and Iceland: A Critique, in: Encyclopedia of Life Sciences. Wiley. 10.1002/9780470015902.a0005180.pub2

Auerbach, B.M., Jackson, F.L.C., Berry, S.D., Blakey, M.L., Caldwell, J., Clinton, C., Graves, J.L., Jones, J.B., Lofaro, E.M., Malhi, R.S., Mosley, C.V., Stubblefield, P.R., 2026. AABA Task Force on the Ethical Study of Human Remains Recommendations: Proposal for the Management and Oversight of Community Partnership and Ethical Stewardship of Human Remains. Am. J. Biol. Anthropol. 189, e70213. 10.1002/ajpa.70213

Bader, A.C., Carbaugh, A.E., Davis, J.L., Krupa, K.L., Malhi, R.S., 2023. Biological samples taken from Native American Ancestors are human remains under NAGPRA. Am. J. Biol. Anthropol. 181, 527–534. 10.1002/ajpa.24726

Barber, A., Brock, T., 2024. A Past Forgotten: A Look at Governmental Efforts to Recover and Restore Historic African American Cemeteries. Wake For. Law Rev. 59.

Beguelin, M., Menéndez, L., 2025. The Pucciarelli Craniofunctional LegacyDatabase: an evaluation of its usage in light of Open Data ideas. Am. J. Biol. Anthropol. 186, 13–14. 10.1002/ajpa.70031

Benn Torres, J., 2020. Anthropological perspectives on genomic data, genetic ancestry, and race. Am. J. Phys. Anthropol. 171, 74–86. 10.1002/ajpa.23979

Blakey, M.L., 2010. African Burial Ground Project: paradigm for cooperation? Mus. Int. 62, 61–68. 10.1111/j.1468-0033.2010.01716.x

Boast, R., Enote, J., 2013. Virtual Repatriation: It Is Neither Virtual nor Repatriation, in: Heritage in the Context of Globalization, SpringerBriefs in Archaeology. Springer New York, New York, NY, pp. 103–113. 10.1007/978-1-4614-6077-0_13

Borgelt, T.S., Goliath, J.R., Waxenbaum, E.B., 2025. Overcoming Systemic Barriers in US Forensic Anthropology Education: Considering Underlying Barriers to Diversity, Equity, Inclusion, and Belonging in Forensic Anthropology. Am. J. Biol. Anthropol. 187, e70069. 10.1002/ajpa.70069

Boulton, G., 2025. Science as a global public good: A necessary responsibility for science, in: Truth Unveiled. Elsevier, pp. 3–19. 10.1016/B978-0-443-23655-6.00001-0

Boycott, K.M., Parboosingh, J.S., Chodirker, B.N., Lowry, R.B., McLeod, D.R., Morris, J., Greenberg, C.R., Chudley, A.E., Bernier, F.P., Midgley, J., Møller, L.B., Innes, A.M., 2008. Clinical genetics and the Hutterite population: A review of Mendelian disorders. Am. J. Med. Genet. A. 146A, 1088–1098. 10.1002/ajmg.a.32245

Boyer, D.M., Jahnke, L.M., Mulligan, C.J., Turner, T., 2019 Workshop on Data Sharing in Biological Anthropology, 2020. Response to letters to the editor concerning AJPA commentary on “data sharing in biological anthropology: Guiding principles and best practices.” Am. J. Phys. Anthropol. 172, 344–346. 10.1002/ajpa.24065

Capili, B., Anastasi, J.K., 2024. Ethical Research and the Institutional Review Board: An Introduction. AJN Am. J. Nurs. 124, 50–54. 10.1097/01.NAJ.0001008420.28033.e8

Capocasa, M., Battaggia, C., Anagnostou, P., Montinaro, F., Boschi, I., Ferri, G., Alù, M., Coia, V., Crivellaro, F., Bisol, G.D., 2013. Detecting Genetic Isolation in Human Populations: A Study of European Language Minorities. PLoS ONE 8, e56371. 10.1371/journal.pone.0056371

Carroll, S.R., Garba, I., Figueroa-Rodríguez, O.L., Holbrook, J., Lovett, R., Materechera, S., Parsons, M., Raseroka, K., Rodriguez-Lonebear, D., Rowe, R., Sara, R., Walker, J.D., Anderson, J., Hudson, M., 2020. The CARE Principles for Indigenous Data Governance. Data Sci. J. 19, 43. 10.5334/dsj-2020-043

Carroll, S.R., Herczog, E., Hudson, M., Russell, K., Stall, S., 2021. Operationalizing the CARE and FAIR Principles for Indigenous data futures. Sci. Data 8, 108. 10.1038/s41597-021-00892-0

Cavalli-Sforza, L.L., 2005. The Human Genome Diversity Project: past, present and future. Nat. Rev. Genet. 6, 333–340. 10.1038/nrg1596

Claw, K.G., Anderson, M.Z., Begay, R.L., Tsosie, K.S., Fox, K., Garrison, N.A., Summer internship for INdigenous peoples in Genomics (SING) Consortium, Bader, A.C., Bardill, J., Bolnick, D.A., Brooks, J., Cordova, A., Malhi, R.S., Nakatsuka, N., Neller, A., Raff, J.A., Singson, J., TallBear, K., Vargas, T., Yracheta, J.M., 2018. A framework for enhancing ethical genomic research with Indigenous communities. Nat. Commun. 9, 2957. 10.1038/s41467-018-05188-3

Clever, I., Jong, L., 2025. Old Bones in New Databases: Historical Insights Into Race, Statistics, and Ancestry Estimation in Anthropology. Am. Anthropol. 127, 566–580. 10.1111/aman.28095

Clinton, C.K., Jackson, F.L.C., 2021. Historical overview, current research, and emerging bioethical guidelines in researching the New York African burial ground. Am. J. Phys. Anthropol. 175, 339–349. 10.1002/ajpa.24171

Cole, N.L., Kormann, E., Klebel, T., Apartis, S., Ross-Hellauer, T., 2024. The societal impact of Open Science: a scoping review. R. Soc. Open Sci. 11, 240286. 10.1098/rsos.240286

Coletta, D.K., Hlusko, L.J., Scott, G.R., Garcia, L.A., Vachon, C.M., Norman, A.D., Funk, J.L., Shaibi, G.Q., Hernandez, V., Filippis, E.D., Mandarino, L.J., 2021. Association of EDARV370A with breast density and metabolic syndrome in Latinos. PLOS ONE 16, e0258212. 10.1371/journal.pone.0258212

Comas, J., 1961. “Scientific” Racism Again? Curr. Anthropol. 2, 303–340. 10.1086/200208

Cornwall, J., Champney, T.H., De La Cova, C., Hall, D., Hildebrandt, S., Mussell, J.C., Winkelmann, A., DeLeon, V.B., 2024. American Association for Anatomy recommendations for the management of legacy anatomical collections. Anat. Rec. 307, 2787–2815. 10.1002/ar.25410

Dadich, A., Moore, L., Eapen, V., 2019. What does it mean to conduct participatory research with Indigenous peoples? A lexical review. BMC Public Health 19, 1388. 10.1186/s12889-019-7494-6

Delgado, M., Ramírez, L.M., Adhikari, K., Fuentes-Guajardo, M., Zanolli, C., Gonzalez-José, R., Canizales, S., Bortolini, M.-C., Poletti, G., Gallo, C., Rothhammer, F., Bedoya, G., Ruiz-Linares, A., 2019. Variation in dental morphology and inference of continental ancestry in admixed Latin Americans. Am. J. Phys. Anthropol. 168, 438–447. 10.1002/ajpa.23756

Delgado-Burbano, M.E., 2007. Population affinities of African Colombians to Sub-Saharan Africans based on dental morphology. HOMO 58, 329–356. 10.1016/j.jchb.2006.12.002

Department of Economic and Social Affairs, United Nations Secretariat, 2009. State of the World’s Indigenous Peoples. United Nationas, New York. Department of Health, Education, and Welfare, U.S., 1979. The Belmont Report.

Douglas-Jones, R., Walford, A., Seaver, N., 2021. Introduction: Towards an anthropology of data. J. R. Anthropol. Inst. 27, 9–25. 10.1111/1467-9655.13477

Echo-Hawk, R., 2010. Working together on race. SAA Archaeol. Rec. 10, 6–9.

Edgar, H., Berry, S., 2025. Ethics of new legacy data: balancing community interests in the use of medicolegal records from contemporary populations. Am. J. Biol. Anthropol. 186, 46. 10.1002/ajpa.70031

ERC Scientific Council, 2022. Open research data and data management plans: Information for ERC grantees, version 4.1. European Research Council, European Comission.

Fasale, M., Rao, D., Panwar, S., 2024. Interceptive correction of winged incisors using a novel 2 × 4 appliance: A case series. Int. J. Oral Health Sci. 14, 82. 10.4103/ijohs.ijohs_33_22

Fforde, C., 2020. Vermillion Accord on Human Remains (1989) (Indigenous Archaeology), in: Smith, C. (Ed.), Encyclopedia of Global Archaeology. Springer International Publishing, Cham, pp. 11016–11019. 10.1007/978-3-030-30018-0_23

Fforde, C., McKeown, C.T., Keeler, H. (Eds.), 2020. The Routledge Companion to Indigenous Repatriation: Return, Reconcile, Renew, 1st ed. Routledge. 10.4324/9780203730966

Fitzpatrick, E.F.M., Martiniuk, A.L.C., D’Antoine, H., Oscar, J., Carter, M., Elliott, E.J., 2016. Seeking consent for research with indigenous communities: a systematic review. BMC Med. Ethics 17, 65. 10.1186/s12910-016-0139-8

Font-Porterias, N., McNelis, M.G., Comas, D., Hlusko, L.J., 2022. Evidence of Selection in the Ectodysplasin Pathway among Endangered Aquatic Mammals. Integr. Org. Biol. 4, obac018. 10.1093/iob/obac018

Francis, E., Asker, C., Tischler, V., 2025. Testing ethical disagreement on ancestral human remains in museums. Int. J. Cult. Prop. 32, 355–364. 10.1017/S094073912610023X

Fuentes, A., 2010. The new biological anthropology: Bringing Washburn’s new physical anthropology into 2010 and beyond-The 2008 AAPA luncheon lecture. Am. J. Phys. Anthropol. 143, 2–12. 10.1002/ajpa.21438

Fumagalli, M., Moltke, I., Grarup, N., Racimo, F., Bjerregaard, P., Jørgensen, M.E., Korneliussen, T.S., Gerbault, P., Skotte, L., Linneberg, A., Christensen, C., Brandslund, I., Jørgensen, T., Huerta-Sánchez, E., Schmidt, E.B., Pedersen, O., Hansen, T., Albrechtsen, A., Nielsen, R., 2015. Greenlandic Inuit show genetic signatures of diet and climate adaptation. Science 349, 1343–1347. 10.1126/science.aab2319

Glantz, L.H., Annas, G.J., Grodin, M.A., Mariner, W.K., 1998. Research in Developing Countries: Taking “Benefit” Seriously. Hastings Cent. Rep. 28, 38. 10.2307/3528268

Godard, B., Schmidtke, J., Cassiman, J.-J., Aymé, S., 2003. Data storage and DNA banking for biomedical research: informed consent, confidentiality, quality issues, ownership, return of benefits. A professional perspective. Eur. J. Hum. Genet. 11, S88–S122. 10.1038/sj.ejhg.5201114

Gould, S.J., 1981. The mismeasure of man. Norton, New York.

Gould, S.J., 1978. Morton’s Ranking of Races by Cranial Capacity: Unconscious Manipulation of Data May Be a Scientific Norm. Science 200, 503–509. 10.1126/science.347573

Graves, J.L., 2021. Human biological variation and the “normal.” Am. J. Hum. Biol. 33, e23658. 10.1002/ajhb.23658

Green, C., 2023. Big Data in Archaeology, in: Handbook of Archaeological Sciences. John Wiley & Sons, Ltd, pp. 1249–1259. 10.1002/9781119592112.ch63

Grupe, G., Wahl, J., 2018. Changing Perceptions of Archaeological Human Remains in Germany, in: O’Donnabhain, B., Lozada, M.C. (Eds.), Archaeological Human Remains. Springer International Publishing, Cham, pp. 81–92. 10.1007/978-3-319-89984-8_6

Gulcher, J.R., Stefánsson, K., 2000. The Icelandic Healthcare Database and Informed Consent. N. Engl. J. Med. 342, 1827–1830. 10.1056/NEJM200006153422411

Halmai, N.B., Taitingfong, R., Jennings, L.L., Yracheta, J., Garba, I., Lund, J.R., Curley, C.A., Claw, K.G., Taualii, M., Garrison, N.A., Carroll, S.R., 2025. Indigenous Data Sovereignty in Genomics and Human Genetics: Genomic Equity and Justice for Indigenous Peoples. Annu. Rev. Genomics Hum. Genet. 26, 375–400. 10.1146/annurev-genom-022024-125543

Hanihara, T., 2008. Morphological variation of major human populations based on nonmetric dental traits. Am. J. Phys. Anthropol. 136, 169–182. 10.1002/ajpa.20792

Hanna, M.G., 2003. Old Bones, New Reality: A Review of Issues and Guidelines Pertaining to Repatriation. Can. J. Archaeol. J. Can. Archéologie 27, 234–257.

Haque, T., Anadon, E.M., Choy, K.W., Chu, E.Y., Heo, C.C., Koesbardiati, T., Lee, W.H., Liu, C., Murti, D.B., Nhoem, S., Rattanachet, P., Saraka, E.M.U., Tantuico, K.F.C., Tran, M., Villaluz, S.A., Yeo, W.X., Wangthongchaicharoen, N., Yukyi, N., Yuwono, P., Rivera, M., 2026. Emic–Etic Perspectives on Southeast Asian Cultural Attitudes Surrounding Human Remains. Int. J. Osteoarchaeol. 36, 150–157. 10.1002/oa.70065

Hlusko, L.J., Carlson, J.P., Chaplin, G., Elias, S.A., Hoffecker, J.F., Huffman, M., Jablonski, N.G., Monson, T.A., O’Rourke, D.H., Pilloud, M.A., Scott, G.R., 2018. Environmental selection during the last ice age on the mother-to-infant transmission of vitamin D and fatty acids through breast milk. Proc. Natl. Acad. Sci. 115. 10.1073/pnas.1711788115

Hlusko, L.J., Martínez de Pinillos González, M., Álvarez Fernández, A., Delgado Esteban, B., Delgado, M., Dern, L., Irish, J.D., Kondo, O., Martinón Torres, M., Scott, G.R., Thiebaut, A., Paul, K.S., Pilloud, M., Rathmann, H., Reyes-Centeno, H., Vlemincq-Mendieta, T., 2026. Global dataset of human morphological dental variation categorized into 32 geographic regions [Dataset]. 10.5061/dryad.76hdr7t9x

Hlusko, L.J., McNelis, M.G., 2022. Evolutionary adaptation highlights the interconnection of fatty acids, sunlight, inflammation and epithelial adhesion. Acta Paediatr. 111, 1313–1318. 10.1111/apa.16358

Howells, W.W., 1996. Howells’ craniometric data on the internet. Am. J. Phys. Anthropol. 101, 441–442. 10.1002/ajpa.1331010302

Hudson, M., Garrison, N.A., Sterling, R., Caron, N.R., Fox, K., Yracheta, J., Anderson, J., Wilcox, P., Arbour, L., Brown, A., Taualii, M., Kukutai, T., Haring, R., Te Aika, B., Baynam, G.S., Dearden, P.K., Chagné, D., Malhi, R.S., Garba, I., Tiffin, N., Bolnick, D., Stott, M., Rolleston, A.K., Ballantyne, L.L., Lovett, R., David-Chavez, D., Martinez, A., Sporle, A., Walter, M., Reading, J., Carroll, S.R., 2020. Rights, interests and expectations: Indigenous perspectives on unrestricted access to genomic data. Nat. Rev. Genet. 21, 377–384. 10.1038/s41576-020-0228-x

Irish, J.D., Morez Jacobs, A., Lea, J.M., Girdland Flink, L., Scott, G.R., 2026. Dental morphology and ancient human dispersals within and out of Africa. J. Hum. Evol. 10.1016/j.jhevol.2026.103855

Irish, J.D., Morez, A., Girdland Flink, L., Phillips, E.L.W., Scott, G.R., 2020. Do dental nonmetric traits actually work as proxies for neutral genomic data? Some answers from continental- and global-level analyses. Am. J. Phys. Anthropol. 172, 347–375. 10.1002/ajpa.24052

Jackson, F.L.C., Auerbach, B.M., 2026. Who Speaks for the Dead? Of Communities and Stewardship in Legacy Collections of Human Remains. Am. J. Biol. Anthropol. 189, e70216. 10.1002/ajpa.70216

Jones, S., 2012. Developments in Research Funder Data Policy. Int. J. Digit. Curation 7, 114–125. 10.2218/ijdc.v7i1.219

Kataoka, K., Fujita, H., Isa, M., Gotoh, S., Arasaki, A., Ishida, H., Kimura, R., 2021. The human EDAR 370V/A polymorphism affects tooth root morphology potentially through the modification of a reaction–diffusion system. Sci. Rep. 11, 5143. 10.1038/s41598-021-84653-4

Kimura, R., Yamaguchi, T., Takeda, M., Kondo, O., Toma, T., Haneji, K., Hanihara, T., Matsukusa, H., Kawamura, S., Maki, K., Osawa, M., Ishida, H., Oota, H., 2009. A Common Variation in *EDAR* Is a Genetic Determinant of Shovel-Shaped Incisors. Am. J. Hum. Genet. 85, 528–535. 10.1016/j.ajhg.2009.09.006

Klafehn, E., Pilloud, M., Yukyi, N., Gregoricka, L., Jenkins, T., 2025. Legacy data in bioarchaeological investigations and the need for proper curation of skeletal remains. Am. J. Biol. Anthropol. 186.

Klebel, T., Traag, V., Grypari, I., Stoy, L., Ross-Hellauer, T., 2025. The academic impact of Open Science: a scoping review. R. Soc. Open Sci. 12, 241248. 10.1098/rsos.241248

L’Abbe, E., Kruger, G., Liebenberg, L., Sapo, O., Ridel, A., 2025. Osteometric and morphological data collected from 20th century cadaveric skeletal collections in South Africa. Am. J. Biol. Anthropol. 186.

Lewis, J.E., DeGusta, D., Meyer, M.R., Monge, J.M., Mann, A.E., Holloway, R.L., 2011. The Mismeasure of Science: Stephen Jay Gould versus Samuel George Morton on Skulls and Bias. PLoS Biol. 9, e1001071. 10.1371/journal.pbio.1001071

Lewis, M., Pitt, R., Bohling, S., 2026. Lost or overused: legal, ethical and research imperatives for a centralised human-remains database in the UK. Antiquity 1–17. 10.15184/aqy.2026.10328

Li, Q., Faux, P., Winchester, E.W., Yang, G., Chen, Y., Ramírez, L.M., Fuentes-Guajardo, M., Poloni, L., Steimetz, E., Gonzalez-José, R., Acuña, V., Bortolini, M.-C., Poletti, G., Gallo, C., Rothhammer, F., Rojas, W., Zheng, Y., Cox, J.C., Patel, V., Hoffman, M.P., Ding, L., Peng, C., Cotney, J., Navarro, N., Cox, T.C., Delgado, M., Adhikari, K., Ruiz-Linares, A., 2025. PITX2 expression and Neanderthal introgression in HS3ST3A1 contribute to variation in tooth dimensions in modern humans. Curr. Biol. 35, 131–144.e6. 10.1016/j.cub.2024.11.027

Lindo, J., Rogers, M., Mallott, E.K., Petzelt, B., Mitchell, J., Archer, D., Cybulski, J.S., Malhi, R.S., DeGiorgio, M., 2018. Patterns of Genetic Coding Variation in a Native American Population before and after European Contact. Am. J. Hum. Genet. 102, 806–815. 10.1016/j.ajhg.2018.03.008

Mardini, M., Badawi, A., Zaven, T., Gergian, R., Nikita, E., 2023. Bioarchaeological perspectives to mobility in Roman Phoenicia: A biodistance study based on dental morphology. J. Archaeol. Sci. Rep. 47, 103759.

Márquez-Grant, N., Fibiger, L. (Eds.), 2011. The Routledge handbook of archaeological human remains and legislation: an international guide to laws and practice in the excavation and treatment of archaeological human remains, Routledge handbooks. Routledge, London ; New York.

McHenry, H.M., Delson, E., 2008. Obituary: William White Howells (1908–2005). Am. J. Phys. Anthropol. 135, 249–251. 10.1002/ajpa.20777

Menéndez, L.P., Urban, M., 2025. Tracing Human Diversity in South America’s Southern Cone: Linguistic, Morphometric, and Genetic Perspectives. Am. J. Biol. Anthropol. 187, e70077. 10.1002/ajpa.70077

Mitina, A., Wang, Y., Mair, W., Vanyushkina, A., Anikanov, N., Efimova, O., Guo, S., Mazin, P., Khaitovich, P., 2026. Comparative analysis of milk and brain fatty acids reveals human-specific signatures in brain development. Commun. Biol. 9, 631. 10.1038/s42003-025-09401-0

Morandini, A.C., Adeogun, O., Black, M., Holman, E., Collins, K., James, W., Lally, L., Fordyce, A., Dobbs, R., McDaniel, E., Putnam, H., Milano, M., 2025. Ectodermal dysplasia: a narrative review of the clinical and biological aspects relevant to oral health. Front. Pediatr. 13. 10.3389/fped.2025.1523313

Morton, S.G., 1844. Crania Aegyptiaca; or, observations on Egyptian ethnography, derived from anatomy, history, and the monuments. John Penington, Philadelphia.

Morton, S.G., 1839. Crania Americana; or, a comparative view of the skulls of various aboriginal nations of North and South America: to which is prefixed an essay on the varieties of the human species. J. Dobson, Philadelphia.

Mulligan, C.J., Boyer, D.M., Turner, T.R., Delson, E., Leonard, W.R., 2022. Data sharing in biological anthropology. Am. J. Biol. Anthropol. 178, 26–53. 10.1002/ajpa.24499

Mulligan, C.J., Hunley, K., Cole, S., Long, J.C., 2004. Population genetics, history, and health patterns in Native Americans. Annu. Rev. Genomics Hum. Genet. 5, 295–315. 10.1146/annurev.genom.5.061903.175920

Nash, S.E., Colwell, C., 2020. NAGPRA at 30: The Effects of Repatriation. Annu. Rev. Anthropol. 49, 225–239. 10.1146/annurev-anthro-010220-075435

National Research Council (US), 1997. Evaluating Human Genetic Diversity. National Academies Press, Washington, D.C. 10.17226/5955

Novacescu, D., Dumitru, C.S., Zara, F., Raica, M., Suciu, C.S., Barb, A.C., Rakitovan, M., Armega Anghelescu, A., Cindrea, A.C., Diana, S., Gaje, P.N., 2025. The Morphogenesis, Pathogenesis, and Molecular Regulation of Human Tooth Development—A Histological Review. Int. J. Mol. Sci. 26, 6209. 10.3390/ijms26136209

Office of the Secretary, Interior (United States), 2023. Native American Graves Protection and Repatriation Act Systematic Processes for Disposition or Repatriation of Native American Human Remains, Funerary Objects, Sacred Objects, and Objects of Cultural Patrimony (No. 2023- 27040 (88 FR 86452)). Federal Register.

Organisation for Economic Co-operation and Development (OECD), 2015. Making Open Science a Reality (OECD Science, Technology and Industry Policy Papers No. 25), OECD Science, Technology and Industry Policy Papers. 10.1787/5jrs2f963zs1-en

Paff, S., 2022. Anthropology by Data Science. Ann. Anthropol. Pract. 46, 7–18. 10.1111/napa.12169

Paschetta, C., González-José, R., Luis Lanata, J., 2017. De cómo cruzar fronteras en la ciencia: homenaje a Héctor M. Pucciarelli, IPCSH e IIDyPCa, CONICET. Puerto Madryn y San Carlos de Bariloche, Argentina.

Passalacqua, N.V., Bartelink, E., McQuade, W.E.P., Steadman, D., Boyd, D., Spradley, K., Sauerwein, K., Ho, R., 2025. The Development of Standards for the Ethical Use of Human Skeletal Remains for Education, Research, and Training in Forensic Anthropology. Am. J. Biol. Anthropol. 186, e70022. 10.1002/ajpa.70022

Passalacqua, N.V., Pilloud, M.A., Gruters, G.A., 2014. Letter to the Editor—Professionalism: Ethics and Scholarship in Forensic Science. J. Forensic Sci. 59, 573–575. 10.1111/1556-4029.12433

Pickering, M., 2020. A repatriation handbook: a guide to repatriating Australian Aboriginal and Torres Strait Islander Ancestral Remains. National Museum of Australia Press, Canberra.

Pilloud, M.A., Passalacqua, N.V., Bartelink, E.J., 2025. Forensic Anthropology as Practiced in the United States: Qualifications, Standards, and Ethical Practice. Am. J. Biol. Anthropol. 187, e70119. 10.1002/ajpa.70119

Pontzer, H., Wood, B.M., Raichlen, D.A., 2018. Hunter-gatherers as models in public health. Obes. Rev. 19, 24–35. 10.1111/obr.12785

Pucciarelli, H.M., Neves, W.A., González-José, R., Sardi, M.L., Rozzi, F.R., Struck, A., Bonilla, M.Y., 2006. East–West cranial differentiation in pre-Columbian human populations of South America. HOMO 57, 133–150. 10.1016/j.jchb.2005.12.003

Qaq, R., Franco, A., Manica, S., 2026. Global patterns of dental morphological variation: Revisiting ASUDAS trait frequencies. Morphologie 110, 101100. 10.1016/j.morpho.2025.101100

Quinto Sanchez, M., 2025. The custody and use of data from unidentified bodies for research and teaching in Mexico: the change of paradigm through forensic emergency. Am. J. Biol. Anthropol. 186. 10.1002/ajpa.7031

Rathmann, H., Kyle, B., Nikita, E., Harvati, K., Saltini Semerari, G., 2019. Population history of southern Italy during Greek colonization inferred from dental remains. Am. J. Phys. Anthropol. 170, 519–534. 10.1002/ajpa.23937

Rathmann, H., Lismann, S., Francken, M., Spatzier, A., 2023a. Estimating inter-individual Mahalanobis distances from mixed incomplete high-dimensional data: Application to human skeletal remains from 3rd to 1st millennia BC Southwest Germany. J. Archaeol. Sci. 156, 105802.

Rathmann, H., Perretti, S., Porcu, V., Hanihara, T., Scott, G.R., Irish, J.D., Reyes-Centeno, H., Ghirotto, S., Harvati, K., 2023b. Inferring human neutral genetic variation from craniodental phenotypes. PNAS Nexus 2, pgad217. 10.1093/pnasnexus/pgad217

Rathmann, H., Reyes-Centeno, H., 2020. Testing the utility of dental morphological trait combinations for inferring human neutral genetic variation. Proc. Natl. Acad. Sci. 117, 10769–10777. 10.1073/pnas.1914330117

Rathmann, H., Reyes-Centeno, H., Ghirotto, S., Creanza, N., Hanihara, T., Harvati, K., 2017. Reconstructing human population history from dental phenotypes. Sci. Rep. 7, 12495. 10.1038/s41598-017-12621-y

Rathmann, H., Stoyanov, R., Posamentir, R., 2022. Comparing individuals buried in flexed and extended positions at the Greek colony of Chersonesos (Crimea) using cranial metric, dental metric, and dental nonmetric traits. Int. J. Osteoarchaeol. 32, 49–63. 10.1002/oa.3043

Rivera, M., Savoldi, F., Tsoi, J., Hin Shin Lee, W., 2025. Tracing the history of Hong Kong’s skeletal collections and their use in bioanthropology and forensic anthropology. Am. J. Biol. Anthropol. 186, 139. 10.1002/ajpa.70031

Robbins Schug, G., Halcrow, S.E., De La Cova, C., 2025. They Are People Too: The Ethics of Curation and Use of Human Skeletal Remains for Teaching and Research. Am. J. Biol. Anthropol. 186, e70013. 10.1002/ajpa.70013

Scott, G.R., Irish, J.D., 2017. Human Tooth Crown and Root Morphology. Cambridge University Press.

Scott, G.R., Pilloud, M.A., Navega, D., Coelho, J. d’Oliveira, Cunha, E., Irish, J.D., 2018a. rASUDAS: A New Web-Based Application for Estimating Ancestry from Tooth Morphology. Forensic Anthropol. 1. 10.5744/fa.2018.0003

Scott, G.R., Turner II, C.G., 1997. The anthropology of modern human teeth: dental morphology and its variation in recent human populations. Cambridge University Press, Cambridge.

Scott, G.R., Turner II, C.G., Townsend, G.C., Martinón-Torres, M., 2018b. The anthropology of modern human teeth: dental morphology and its variation in recent and fossil Homo sapiens. Cambridge University Press.

Seguchi, N., Hayashi, A., 2025. Discussions of ethical considerations in the utilization of unpublished legacy data to benefit descendant communities: The case of Japan. Am. J. Biol. Anthropol. 186. 10.1002/ajpa.70031

Serwadda, D., Ndebele, P., Grabowski, M.K., Bajunirwe, F., Wanyenze, R.K., 2018. Open data sharing and the Global South—Who benefits? Science 359, 642–643. 10.1126/science.aap8395

Sholts, S.B., 2025. “To honor and remember”: An ethical awakening to African American remains in museums. Am. J. Biol. Anthropol. 186, e24943. 10.1002/ajpa.24943

Smith-Guzmán, N.E., Smid Núñez, J., Cybulski, J.D., Sánchez Herrera, L.A., 2025. Biodistance Analysis via Dental Phenotypic Diversity in Early Collective Burials at Cerro Juan Díaz, Panamá (30–650 CE) [Análisis de Biodistancia Mediante la Diversidad Fenotípica Dental en los Entierros Colectivos Tempranos del Cerro Juan Díaz, Panamá (30–650 EC)]. Am. J. Biol. Anthropol. 186, e25050. 10.1002/ajpa.25050

Stantis, C., De La Cova, C., Lippert, D., Sholts, S.B., 2023. Biological anthropology must reassess museum collections for a more ethical future. Nat. Ecol. Evol. 7, 786–789. 10.1038/s41559-023-02036-6

Stojanowski, C.M., Paul, K.S., Seidel, A.C., Duncan, W.N., Guatelli-Steinberg, D., 2019. Quantitative genetic analyses of postcanine morphological crown variation. Am. J. Phys. Anthropol. 168, 606–631. 10.1002/ajpa.23778

Suber, P., 2003. The taxpayer argument for open access. SPARC Open Access Newsl.

TallBear, K., 2013. Native American DNA: tribal belonging and the false promise of genetic science. University of Minnesota Press, Minneapolis, MN.

Taylor, L., 2016. The ethics of big data as a public good: which public? Whose good? Philos. Trans. R. Soc. Math. Phys. Eng. Sci. 374, 20160126. 10.1098/rsta.2016.0126

Templeton, A.R., 1999. Uses of Evolutionary Theory in the Human Genome Project. Annu. Rev. Ecol. Syst. 30, 23–49. 10.1146/annurev.ecolsys.30.1.23

Tikhonov, D., 2022. The role of adaptive variants of the EDAR gene in the development of breast cancer (Abstracts of the report at the scientific and practical conference “Molecular and biological mechanisms of human health formation in the North” on November 17–18, 2022, Yakutsk). Sib. Res. 8, 37–39. 10.33384/26587270.2022.08.02.06e

Tishkoff, S.A., Verrelli, B.C., 2003. Patterns of Human Genetic Diversity: Implications for Human Evolutionary History and Disease. Annu. Rev. Genomics Hum. Genet. 4, 293–340. 10.1146/annurev.genom.4.070802.110226

Tsipouri, L., Liarti, S., Vignetti, S., Grapengiesser, I.M., 2025. The economic impact of open science: a scoping review. R. Soc. Open Sci. 12, 250754. 10.1098/rsos.250754

Tsosie, K.S., Yracheta, J.M., Kolopenuk, J., Smith, R.W.A., 2021. Indigenous data sovereignties and data sharing in biological anthropology. Am. J. Phys. Anthropol. 174, 183–186. 10.1002/ajpa.24184

Turner, C.G., Nichol, C.R., Scott, G.R., 1991. Scoring procedures for key morphological traits of the permanent dentition: the Arizona State University Dental Anthropology System., in: Kelley, M., Larsen, C. (Eds.), Advances in Dental Anthropology. Wiley-Liss, New York, pp. 13–31.

Turner, T.R., Mulligan, C.J., 2019. Data sharing in biological anthropology: Guiding principles and best practices. Am. J. Phys. Anthropol. 170, 3–4. 10.1002/ajpa.23909

UNESCO, 2021. UNESCO Recommendation on Open Science. UNESCO. 10.54677/MNMH8546

Van Vaerenbergh, Y., Hazée, S., Zwienenberg, T.J., 2026. Open Science: A Review of Its Effectiveness and Implications for Service Research. J. Serv. Res. 29, 22–44. 10.1177/10946705251338461

Wali, A., Collins, R.K., 2023. Decolonizing Museums: Toward a Paradigm Shift. Annu. Rev. Anthropol. 52, 329–345. 10.1146/annurev-anthro-052721-040652

Wang, Xianwen, Liu, D., Ding, K., Wang, Xinran, 2012. Science funding and research output: a study on 10 countries. Scientometrics 91, 591–599. 10.1007/s11192-011-0576-6

Washburn, S.L., 1951. The new physical anthropology. Trans. N. Y. Acad. Sci. 13, 298–304. 10.1111/j.2164-0947.1951.tb01033.x

Washington, H.A., 2006. Medical apartheid: the dark history of medical experimentation on Black Americans from colonial times to the present, 1st ed. ed. Doubleday, New York.

Wilson, T., 2025. Preserving the Past: Digital Approaches to a Decade of Bioarchaeological Data from Northern Jordan. Am. J. Biol. Anthropol. 186. 10.1002/ajpa.70031

Wu, Y., Li, X., Chen, J., Yang, B., Yang, X., Hou, J., 2026. Genetic analysis of a Chinese family with non-syndromic tooth agenesis may reveal a potential multi-locus etiology. Gene 979, 149915. 10.1016/j.gene.2025.149915

Yang, G., Chen, Y., Li, Q., Benítez, D., Ramírez, L.M., Fuentes-Guajardo, M., Hanihara, T., Scott, G.R., Alonzo, V.A., Jose, R.G., Bortolini, M.C., Poletti, G., Gallo, C., Rothhammer, F., Rojas, W., Zanolli, C., Adhikari, K., Ruiz-Linares, A., Delgado, M., 2023. Dental size variation in admixed Latin Americans: Effects of age, sex and genomic ancestry. PLOS ONE 18, e0285264. 10.1371/journal.pone.0285264

Zhang, W., Vervoort, J., Pan, J., Gao, P., Zhu, H., Wang, X., Zhang, Y., Chen, B., Liu, Y., Li, Y., Pang, X., Zhang, S., Jiang, S., Lu, J., Lyu, J., 2022. Comparison of twelve human milk oligosaccharides in mature milk from different areas in China in the Chinese Human Milk Project (CHMP) study. Food Chem. 395, 133554. 10.1016/j.foodchem.2022.133554

