## Appendix_1 for "DeMoDa: A global open access database for comparative research in human dental morphological variation"

**De**ntal
**Mo**rphological
**Da**tabase
https://demoda.cenieh.es/

### Policies and Procedures

#### Developed June 2-4, 2025
#### Finalized February 27, 2026

#### DeMoDa Scientific Advisory Board

**Chair**, Leslea J. Hlusko, Centro Nacional de Investigación sobre la Evolución Humana

**Scientific Integrity Advisor**, G. Richard Scott, University of Nevada Reno

**Accessibility and Ethics Advisor**, Marin Pilloud, University of Nevada, Reno

**Dental Anthropology Association President**, Kathleen Paul, University of Arkansas

**Board Member and DeMoDa Administrator**, Marina Martínez de Pinillos González, Centro Nacional de Investigación sobre la Evolución Humana

**Board Member**, Miguel Delgado, Facultad de Ciencias Humanas y Sociales, Universidad del Cauca in Popayán, CONICET-Universidad Nacional de La Plata

**Board Member**, Joel D. Irish, Liverpool John Moores University

**Board Member**, Osamo Kondo, The University of Tokyo

**Board Member**, María Martinón Torres, Centro Nacional de Investigación sobre la Evolución Humana

**Board Member**, Mario Modesto Mata, Centro Nacional de Investigación sobre la Evolución Humana

**Board Member**, Hannes Rathmann, Senckenberg, University of Tübingen

**Board Member**, Hugo Reyes-Centeno, University of Kentucky

### Overview

#### Mission

The Dental Morphological Database (DeMoDa) is an online data repository for individual dental morphological trait scores collected by several prolific dental anthropologists. Combined, these scientists characterized the dental variation of more than 40,000 people from around the world, spanning millennia.

DeMoDa was developed in order to preserve these data and to provide a framework for making them available to researchers worldwide, thereby advancing the study of human dental variation.

#### Commitment to Open Science

DeMoDa was explicitly designed with a commitment to Open Science, in accordance with the FAIR Principles[1] and the Open Science initiative of the ERC Horizon Europe programme. As the data in DeMoDa were derived from human biological/skeletal remains, the DeMoDa Advisory Board is committed to balancing scientific value with concerns about how these human remains came to be in the various repositories from which the dental anthropological data were collected. This balance is elaborated on and explained in the Statement on Ethics section that follows.

#### Funding

DeMoDa is made possible by the financial support provided to the Tied2Teeth Project (directed by Leslea Hlusko), granted through the European Research Council within the European Union's Horizon Europe Framework Programme (ERC-2021-ADG, project number 101054659). Views and opinions expressed are, however, those of the author(s) only and do not necessarily reflect those of the European Union or the European Research Council. Neither the European Union nor the granting authority can be held responsible for them.

The CENIEH provides additional project support hosting the database website. CENIEH is supported by the Spanish Ministerio de Ciencia, Innovación y Universidades.

#### Acknowledgements

DeMoDa would not be possible without the assistance of the ERC Project Tied2Teeth's project members, especially Project Manager Inma Villamartín Santamaría and technicians Ana Álvarez and Beatriz Delgado. The dedicated assistance from Softeca is also essential for the development and maintenance of the database. We also thank Softeca for creating the name DeMoDa and the logo. The CENIEH has also provided essential assistance incorporating DeMoDa into the infrastructure; we especially express gratitude to Javier Valladolid Aguinaga, who oversees CENIEH's sistemas.

---

[1] Developed in 2016, the FAIR principles call for data to be Findable, Accessible, Interoperable, and Resusable. https://www.go-fair.org/fair-principles/

### Statement on Ethics

#### General Principles

The 2025 Scientific Advisory Board members represent eight countries across four continents, and therefore, bring to DeMoDa a range of scientific and societal perspectives. The ethical questions related to these legacy data were discussed in-depth during a board meeting on June 2-4, 2025.

The data in DeMoDa are based on observations of human skeletal material. These data were collected in a non-destructive manner. Variations in tooth shape were assessed visually and assigned numerical scores based on comparison with published standards. At the time of collection, all data were obtained in accordance with the applicable governmental laws and institutional policies and procedures in place at that time.

These data are typically based on the study of institutional/museum skeletal collections. The different histories of how human remains were incorporated into institutional collections raise a variety of ethical concerns. Some of the populations included in these institutional collections were incorporated as a consequence of serious inequities and injustices, ranging from societal neglect to racism and, in some instances, genocide. In many cases, institutions lack adequate information to ascertain the provenience context of their collections.

Various professional organizations have been engaging with the ethical issues surrounding the treatment, repatriation, and curation of human remains. These professional and legal entities across the globe are moving to respect the wishes of descendant communities and other communities of care (reflected in the 3rd principle of the Vermillion Accord on Human Remains adopted by the World Archaeological Congress in 1989[2]). There is also a recognition of respect for the potential scientific research value of skeletal remains (the 4th principle of the Vermillion accord). Both of these viewpoints are seen as legitimate and need to be balanced (5th and 6th principles of the Vermillion accord).

In contrast to the direct study of human remains, there has been less ethical engagement with the use of the legacy data collected from human remains, particularly data that has already been published either in full or in part (such as the dental morphological data in DeMoDa). Given this current situation, the Advisory Board recognizes that the decisions made and actions taken during their first meeting (June 2-

[2] https://worldarchaeologicalcongress.com/code-of-ethics/

4, 2025) will provide a point of debate and discussion for how legacy data are managed in the future.

The Board set the six principles of the World Archaeology Congress' Vermillion Accord (1989) as the foundation to which DeMoDa adheres, holding respect for both societal concerns and scientific value, noting that the application of the six principles for legacy data may differ in some ways from how they are applied to human remains. We also incorporate the points raised by the Global Indigenous Alliance's CARE principles, that data be shared following both FAIR and also respect Indigenous communities' through an evaluation of the Collective benefit, Authority to control, Responsibility, and Ethics.[3] The numerous academic and governmental organizations from around the world that have expanded on the Vermillion principles in regards to human remains provide essential perspectives that the DeMoDa Scientific Advisory Board brought into consideration.

The Board will continue to assess and discuss DeMoDa's impact in both scientific and societal realms, recognizing the dynamic nature of laws and policies across the world (that cannot be fully represented by any one committee). We welcome and appreciate feedback on our chosen approach (described below). We envision that the decisions and policies of our Board will be adjusted as the scientific community, descendant communities, communities of care, and other stakeholders have the opportunity to respond to this first step. We hope that our approach facilitates discussions and provides an opportunity to reimagine ethical futures of research and data collection from human remains.

[3] https://www.gida-global.org/care

#### Putting the Ethical Principles into Action

The study of human variation is historically rooted in efforts to identify similarities and differences between human populations - sometimes with the intent of using those differences to justify mistreatment, a discipline referred to as scientific racism, framed into colonial practices. Although the academic discipline has largely shifted to more diverse and biologically grounded motivations for studying human variation, the legacy of harm still lingers. For many of the communities harmed by scientific racism, the wounds remain raw.

##### The potential benefit

The DeMoDa Scientific Advisory Board recognizes that the information preserved in DeMoDa provides data that can serve to counter the historical typological studies that led to injustices. The data in DeMoDa preserve a much more nuanced record of continuous human dental variation, not only in terms of the scope of the dataset, but also in its reporting of raw data for individuals.

The DeMoDa Scientific Advisory Board notes that these historical data provide a unique and important opportunity to modernize studies of human dental variation at the species level, with relevance to biological, biomedical, and evolutionary research. The potential benefits to society ranges from:

- the basic biological research from which new insights about human physiology are derived, to
- a more realistic view of human variation that is continual and complex, and can serve to counter the racialized analyses of the past, as well as
- enabling descendant groups to access data of ancestral communities and become active stewards and researchers.

##### The potential for harm

The challenge that the DeMoDa Scientific Advisory Board discussed at-length is how to balance the value of the scientific insights with the negative impacts that may be experienced by some communities and populations whose dental variation is included in the database. The Advisory Board deeply respects that the wishes of descendant communities should have weight in how the dental variation of their ancestors is used to study the histories of their communities. Neglecting to do this type of consultation prior to the analysis of a range of biological data has led to scientific studies that are at odds with the values and beliefs of some communities, furthering the harm inflicted by scientific racism.

The dental morphological scores in DeMoDa are from hundreds of repositories across the globe, representing many hundreds of descendant communities. While the DeMoDa Scientific Advisory Board respects and appreciates the intentions behind the calls for individual community consultation, in compiling the DeMoDa data, it was immediately clear that this type of population-by-population consultation would not be feasible. It is not always possible to re-trace the data-collection steps of each researcher as the depth of detail that would be necessary was not recorded or no longer exists.

One option to address this situation is to develop a standardized approach for all populations. However, from a global perspective, community concerns are so variable that there is no one policy that would be appropriate for the full range of populations represented in DeMoDa.

Another option the Scientific Advisory Board considered is to not share any data until this challenge is addressed. This option could well render these historical data moot, despite their potential for societal benefit, and rendering past dental anthropology research irreproducible.

##### A balanced step forward

In order to balance the value and potential harm embedded in the database, a curated dataset of worldwide human dental morphological variation is being released in order to achieve the goal of promoting the study of human variation at the global scale. These comprehensive dental data are made public with less granular information about the populations and the individuals within each population. The Scientific Advisory Board hopes that this combination of data-availability with population and individual de-identification will adequately address the most serious ethical concerns.

Two steps were taken to de-sensitize the contextual data for the dental morphological scores published in this curated global dataset. Rather than using names of populations, or even providing detailed geographical locations, we compiled data into geographical regions so that no one particular population can be identified. We also removed the individual repository identification numbers for the human remains, so that it is not possible to recover this identifying information. In an effort towards transparency, we provide a separate list of the populations included in the database to be available for descendant communities and communities of care to evaluate.

For the time being, DeMoDa will serve as a secure repository for the more-detailed contextual data (individual and population identifications) until communities with a vested concern have had a chance to provide feedback and an appropriate next-step is determined.

### The Scientific Advisory Board

#### Charge

The DeMoDa Scientific Advisory Board is charged with oversight of the database, and determining an evolving policy for access, maintenance, and elaboration. Members of the Board are also responsible for nominating, voting for, and appointing other members as needed. All members have voting privileges.

#### Function

The work of the Scientific Advisory Board is generally done by consensus. When contentious issues arise, Robert's Rules of Order provide the structure for resolution, and a voting process is used.

Board votes are required for the appointment of Board Members, DeMoDa Technicians and Administrator users, major structural changes to the database, and changes to the policies and procedures described herein.

While DeMoDa is fully supported by the ERC Tied2Teeth project, the Chair of the Board (and Principal Investigator of the Tied2Teeth Project) will have the power to make executive decisions. When DeMoDa is supported by a more diverse range of funding, the Chair will make decisions based on majority vote by the Board.

#### Structure

The DeMoDa Scientific Advisory Board consists of the Chair, the Scientific Integrity Advisor, the Accessibility and Ethics Advisor, current President of the Dental Anthropology Association, and between 6 to 10 Board Members. At least one Board Member will serve as an "administrator" for the database. While the DeMoDa website is hosted at CENIEH, at least one of the Regular Members needs to be part of the scientific staff at CENIEH. The representation of the membership should aim to be diverse in terms of geographic and disciplinary representation. In total, the Board will have between 10 and 14 members.

At least one full meeting of the Scientific Advisory Board takes place each calendar year, either in person or by video conference. A quorum is defined as two-thirds of the Board.

##### Chair

**Appointment:** The DeMoDa Scientific Advisory Board Chair will be appointed by a 75% approval vote of the DeMoDa Scientific Advisory Board. Nominations for this position are provided by current and past members of the Advisory Board.
**Expectations and Responsibilities:**

- Serves as the primary contact for DeMoDa
- Provides general oversight of the database and website, ensuring that the policies and procedures approved by the Advisory Board are followed
- Is responsible for communicating with the Advisory Board on at least a quarterly basis, and convening at least one live meeting a year, either in-person or via video call
- Is responsible for keeping track of Advisory Board service terms and ensures that positions are filled as terms come to a close

**Term limit:** Five years with possibility to be reappointed without limit, pending 75% approval of the Advisory Board. The Accessibility and Ethics Advisor will be responsible for managing a re-appointment vote.

#### Scientific Integrity Advisor

**Appointment:** The scientific integrity advisor will be appointed by a simple majority vote of the Advisory Board. Nominations will be provided by current and past members of the Advisory Board.
**Expectations and Responsibilities:**
- Respond to queries about the composition of the data in the database
- Raise concerns as needed
- Guide decisions on data quality

**Term limit:** Five years with possibility to be reappointed without limit, pending simple majority approval of the Advisory Board.

#### Accessibility and Ethics Advisor

**Appointment:** The scientific integrity advisor will be appointed by a simple majority vote of the Advisory Board. Nominations will be provided by current and past members of the Advisory Board.
**Expectations and Responsibilities:**
- Guide oversight of ethical issues (e.g., the ethics surrounding data sharing, data accessibility and distribution, and treatment of human remains and legacy data)
- Monitor how DeMoDa is being perceived by professional, descendant, and other relevant communities

**Term limit:** Five years with possibility to be reappointed without limit, pending simple majority approval of the Advisory Board.

#### President of the Dental Anthropology Association

**Appointment:** The President of the Dental Anthropology Association will be appointed as per the Bylaws and Constitution of the Dental Anthropology Association.
**Expectations and Responsibilities:**
- Serves as the primary liaison between the DAA membership and DeMoDa board
- Facilitates the promotion of DeMoDa
- Works with DeMoDa board to ensure development and sustainability

**Term limit:** Three years as per the bylaws of the DAA

#### Board Members

**Appointment:** There shall be at least 6 and no more than 10 Board members. The qualifications and experiences of nominees should be related to the mission of DeMoDa. Board Members will be appointed by a simple majority vote of the Advisory Board. While the website is hosted by CENIEH, at least one member should be a staff member of CENIEH. At least one Board member will serve as an "administrator" of the database. The Advisory Board will endeavor to ensure diversity in terms of global and scientific sub-disciplinary representation.

**Expectations and Responsibilities:**

- Attend regular meetings of the Advisory Board and provide feedback to communications sent by email
- Be aware of the mission, policies, and procedures of DeMoDa and provide guidance to make sure that these are being followed
- Serve as voting members of the advisory board
- Be engaged by bringing concerns and ideas to the full Board

**Term limit:** Board Members hold a 3-year appointment on the Advisory Board with the potential to serve two consecutive terms. Members can rejoin the Board after being away for a period of at least 6 years, with two Board members being replaced (or re-appointed) approximately every 2 years.

### Technical Structure and Support

#### URL hosting

The Centro Nacional de Investigación sobre la Evolución Human (CENIEH), the Spanish National Center for Research on Human Evolution hosts DeMoDa's URL.

#### Website technical support

Currently provided by Softeca (https://www.softeca.es/).

#### Financial support

The Tied2Teeth project will financially and logistically support the database through October 2027.

***Proposal**: The Dental Anthropology Association will provide limited financial support to cover the cost of the Softeca yearly maintenance contract. This will be brought to the DAA Executive Committee for discussion.*

### Types of DeMoDa Users

The current structure of DeMoDa provides for four types of users: public access, Downloaders, Technicians, and Administrators. These are described in detail below.

#### Landing-page, public access

In addition to being a resource for scientific researchers, DeMoDa has a public-facing component. Through the landing page (https://demoda.cenieh.es), anyone on the internet has the ability to access a specific range of dental morphological traits for publicly available populations and view these trait frequencies by population plotted on a world map. These data are not downloadable and are not visible as specific populations or by individual data.

#### Downloaders

The DeMoDa Advisory Board has paused the establishment of this role, as the data in DeMoDa are to be kept private for the time being to satisfy ethical concerns. The published dataset will serve as the fulfillment of the mission of DeMoDa.

#### Technician

Users who have Technician status will have access to enter data into DeMoDa individual-by-individual using the data-entry template.

##### Requirements

Approval by the Advisory Board or the designated proxy

##### Appointment process

Technicians are appointed by the Chair of DeMoDa or their proxy (administrators). All technicians need to be approved by the DeMoDa Advisory Board by simple majority vote.

##### Responsibilities

- Follow the policies and procedures of DeMoDa
- Maintain confidentiality of the data
- Enter data with good faith to ensure data integrity
- Bring concerns to the attention of the Chair of the DeMoDa Advisory Board, or if the concerns involve the Chair, to another member of the Board

##### Cause for revocation of access

Technician access to DeMoDa can be revoked at any time for any reason deemed of sufficient concern by a majority of the Advisory Board. The primary reasons for revocation of access will be based primarily on use or treatment of data that does not follow the policies, procedures, and/or intent of DeMoDa. Any Technician who operates outside of DeMoDa's regulations and ethical expectations  will lose access immediately until a resolution is reached by the Advisory Board.

#### Administrator

Users who have Administrator status will have access similar to a technician. In addition, they can approve and oversee changes and additions to the database.

##### Requirements

Approval by the Advisory Board or the designated proxy

##### Appointment process

Administrators are appointed by the Chair of DeMoDa or their proxy. All administrators need to be approved by the DeMoDa Advisory Board by simple majority vote.

##### Responsibilities

- Follow the policies and procedures of DeMoDa
- Maintain confidentiality of the data
- Liaise with the technical support
- Enter data with good faith to ensure data integrity
- Approve changes to the database with good faith and to ensure data integrity
- Respond to user access requests
- Bring concerns to the attention of the Chair of the DeMoDa Advisory Board, or if the concerns involve the Chair, to another member of the Board

##### Cause for revocation of access

Administrator access to DeMoDa can be revoked at any time for any reason deemed of sufficient concern by a majority of the Advisory Board. The primary reasons for revocation of access will be based primarily on use or treatment of data that does not follow the policies, procedures, and/or intent of DeMoDa. Any Administrator who operates outside of DeMoDa's regulations and ethical expectations will lose access immediately until a resolution is reached by the Advisory Board.

### Use of the Data

#### Ethical Approaches

In the spirit of open access of data and the principles of FAIR, these edited data (as described in the ethical statement) are shared freely. The Board realizes that, also in the spirit of academic freedom, we cannot control how the data are used, nor of the questions that are asked. However, we encourage researchers to consider implementing an ethical framework for research that stems from these data and apply CARE principles in tandem.[4] We advocate that these data be used to pursue hypotheses that serve to understand our shared human past and evolution as well as biology, in a way that does not racialize or marginalize populations. Here we provide a series of questions that may help to guide hypothesis-formation and interpretation of the results:

- Is the scientific answer to my research question going to provide more benefit to humanity than harm to any specific community?
    - *If yes, proceed.*
- Does this research approach the data from a typological perspective?
    - *If you answer yes to this question, consider re-framing your question/hypothesis.*
- Does this research have the potential to harm any of the descendant communities that may be represented in the analyses?
    - *If you answer yes to this question, consider re-framing your question/hypothesis.*
- Is there an opportunity to involve descendant communities in collaboration or consultation?
    - *If you answer yes to this question, consider taking the time and making the effort to make those contacts.*

#### Citation

When the DeMoDa curated dataset is used in academic research, please reference the concept of the Dental Morphological Database described in this peer-reviewed publication:

*Forthcoming…*

As well as the actual dataset:

*Citation to the DeMoDa curated dataset that will be published on Dryad (following the Data Management plan that the Tied2Teeth project has with the ERC)*.

---

[4] https://www.nature.com/articles/s41597-021-00892-0
