## Appendix_2 for "DeMoDa: A global open access database for comparative research in human dental morphological variation"

**Appendix 2. Populations combined into the 32 geographic regions of DeMoDa Dataset v1**

| Geographic region | Population name or geographic location |
|---|---|
| Arctic | Chukchi [Tchoukotka; Sedentaire du village Tschetschine / Chukchis] |
| | Eskimo Alaska |
| | Southern Alaska] |
| | Bay, Wahle Sound; Lancaster Sound; Melville Peninsula / Canadian Eskimos] |
| | Peninsula / Canadian Eskimos] |
| | Wolstenbolme Sound; Upernavic; Sepulchre; Issungvak / Greenland] |
| | Upernavic / Greenland Eskimos] |
| | Eskimo Siberia [Indian Point; Port Providence, Plover Bay : East Siberia / Siberian Eskimos] |
| | EU Lapp [Lapps / North Europe] |
| | Karel I |
| | Karel II |
| | Lapps |
| | Mortensuoes, East Finmark, Norway / Lapps |
| | Nivkh [Nivkh (Giliakh) / NE Siberia] |
| | Point Hope |
| | Saint Lawrence |
| | Yukagir [Yukagir / NE Siberia] |
| Australia | Australia North |
| | Australia Murray River [Roonka site / Murray river basin] |
| | Australia North Queensland |
| | Woollahra; Cronulla; Lake Illawarra; La Perouse / New South Wales] |
| | Australia NSW [Sand Dunes, N. Cronulla; Bondi; Hnters Bay; Woolwish Dock, Hunters Hill; Lake Illawarra; Grey's Point; Quibray Bay; Forty Baskets Bay; near Sydney / New South Wales] |
| | Australia NT [Herbert River / Northern Territory] |
| | Australia NT [Millinginbi Island, Arnhem Land; Port Darwin / Northern Territory, Australia] |
| | Australia Queensland [Port Essington; Moreton Bay / Queensland, Australia] |
| | : near Adelaide / South Australia] |
| | Dry Creek; Snowtown; Largs Bay : Near Adelaide / South Australia] |
| | Australia Tasmania [Tasmania / Australia] |
| | Australia Unknown [Australia / Locality Unknown] |
| | Australia Victoria [Hexham, Victoria / Australia] |
| | Australia Victoria [Koondrook; Loddon River; Woorndoo; Mortlake / Victoria, Australia] |
| | Australia WA [Shark's Bay; Murchison River / West Australia] |
| | Australia WA [Western Australia] |
| | Australian Northern Territory NT |
| | Bartolombo Bay, North Australia |
| | Murchison River, South Australia |
| | Murray River, South Australia |
| | New South Wales, South Australia |
| | Point Darwin Tribe, North Australia |
| | South Australia |
| | Swan Reach, South Australia |
| | White Cliffs, New South Wales, South Australia |
| Central America | Carib / Nicaragua |
| | Carib [Santo Domingo; Landings bay / Carib] |

| | |
|---|---|
| | Guatuoso Indian / Costa Rica |
| | Jamaica [Carib, Jamaica] |
| | Jamaica [Utilla Bay Island / Jamaica / Central America] |
| | Jarasco; Bolson de Mapiani; Yucatan; Chihuahua; Milta Chareas, San Luis Potosi; Cora; Sierra Madre of |
| | Panama 1 |
| | Panama 2 |
| | Tlatelolco |
| | Venado Beach / Panama |
| Central Asia | Afghanistan |
| | Kafiristan; Ghazni; Peshawar / Pakistan |
| | Kazaks |
| | Panshir Valley; Helmand Valley; Cabul / Afghanistan |
| | Turkmen |
| | Uzbek 1 |
| | Uzbek 2 |
| | Uzbek 3 |
| Central Europe | Hungary |
| | Berlin / Germany |
| | Bludau, Machlenburh, Berlin, Heidenheim, Stuttgart / Germany |
| | Berlin, Heidenheim, Stuttgart / Germany |
| | Berne / Switzerland |
| | Buda-Pesth; Stuhlweissenberg; Pressburg; Csakovar; Nagy-Sap; Demko-Hegy / Hungary |
| | Charvaty, near Olomouc, Central Moravia / Czecho |
| | Charvaty, nr Olomouc, Central Moravia / Czecho |
| | Lenz Canton, Grisons / Switzerland |
| | Russniak, Pelabreyk from Galicia; Tarnapol / Poland |
| | Vienna; Modling, near ViennaRaach Gloggnitz; Malta Carinthia; Paach / Austria |
| | Vienna; Tyrol; Modling, near ViennaRaach Gloggnitz; Malta Carinthia; Paach / Austria |
| | Warsaw; Masure / Poland |
| Eastern Africa | Abyssinia, near Sepape / Ethiopia |
| | Adrian Battiste, Bantu / Mozambique |
| | Dhol Banta tribe, Berbera; Ogaden, Darod Kuhar; Burao Dist.; Erigavo Dist.; Kismayu / Somalia |
| | Dohlbanta tribe; Ef Der Erigavo District; Ogaden Somali, Parod Kuhar II; Derod or Hawiya / Somalia |
| | From a cave near Marandellas, Salisbury, South Rhodesia / Zimbabwe |
| | Haya |
| | Inchila Plain, nr Mt. Mlanje, Nyassaland / Malawi |
| | Kikuyu, Kaurite to Fort Hall; Nairobi; Elementeites, Naishi Valley; Turkana; Haya tribe / Kenya |
| | Kikuyu, Nairobi / Kenya |
| | Livingstonia / Zambia |
| | Maranda, Loangwe River / Zambia |
| | Mhuma / Ethiopia |
| | Mtussi; Mhutu / Rwanda |
| | Naivasha; Mt. Elgon, Basia Native / Uganda |
| | NW coast of Lake Nyassa, Lyassaland; Albert Edward Nyanza Lake / Malawi |
| | Pare, Gonja; Haya tribe, Lake Victoria's Island; Angoni, Nyasaland / Tanzania |
| | Pare, Gonja; Haya tribe; M'bena, Wabena, Ulanga Valley / Tanzania |
| | Teita |

| | |
|---|---|
| | Teso tribe, East Province; Ankole; Wakmia, Chif Ruakirara / Uganda |
| Eastern Mediterranean/Levant | 26th - 3Ath Dynasty, 664 - 343BC, Gizeh / Egypt |
| | Ain el Turba, El Baguat, Kharga / After Christian Period / Egypt |
| | Aintab; Adalia : Mohammedan, Armenian, Kurd; Scutari; Silenlen / Turkey |
| | Bosporus / Turkey |
| | Aleppo; Damascus / Syria |
| | Ancient Cyprus, Roman and Hellenismic / Cyprus |
| | Arab, Syria / Bedouins |
| | Baghdad, Karbala, Najaf / Iraq, Arab |
| | Cyprus Island / Middle and Late Bronze age, Lapethos |
| | Druze, Beirut; Maronite, Abech / Lebanon |
| | Early Christian and Christian periods / Nubia |
| | Early Christian Period / Egypt |
| | Early Christian period / Nubia |
|  | Hellenismic Period; Lapethos / Cyprus |
| | Kerma / Nubia |
| | Kish, ca 4,AAA years BP, early Dynasty / Iraq |
| | Near Persepolis / Iran |
| | P2-P3 Dynasty, Kerma / Nubia |
| | Pre-Dynastic period, Badari / Egypt |
| | Pre-Dynastic period, Naqada / Egypt |
| | Samarita / Palestine |
| | Sesebi / Nubia |
| | Sesebi, recent Nubia |
| | Tell Cuweir (Lachish), Bronze and Iron age / Palestine, Israel |
| | XIIth Dynasty / Egypt |
| | XX - XXVth Dynasty / Egypt |
| | XX-XXVth Dynasty, including a few samples from XVIII-XXI Dynasty / Egypt |
| | XX-XXVth Dynasty, including a few samples from XVIII-XXIth Dynasty / Egypt |
| Eastern South America | Buracao |
| | Cerca Grande (Site I) |
| | Corondó |
| | Gentio Cave II |
| | Lagoa Santa 1 |
| | Lagoa Santa 2 |
| | Lagoa Santa 3 |
| | Lapa da Gruta das Boleiras |
| | Sambaqui (South) |
| | Sambaqui Boguassi I |
| | Sambaqui Boguassi II |
| | Sambaqui de Piacaguera |
| | Tenorio |
| Eastern Siberia | Troitskoye |
| First Island Chain (Taiwan to Sakhalin) | Ainu from Hidaka, Nibutani, Shizunai, etc. / Hanihara Collection, Univ. of Tokyo |
| | Atayal / Taiwan |
| | Gornozavodsk and Nevelish / Sakhalin |
| | Gornozavodsk and Nevelisk / Sakhalin Ainu // Sakhalin Ainu |
| | Miyako Island, Hirayoshi Junior High School / University of the Ryukyus |

| | |
|---|---|
| | Neolithic Jomon / Shell mound site |
| | Recent Japanese |
| Iberian Peninsula | Neolithic - Bronze Sapin |
| | Maderia; Oporto / Portugal |
| | Zaraus, Province of Guipuscoa; Genista Cave, Gibraltar / Spain |
| Inner Asia | Evenki / NE Siberia |
| | Late Broze early Iron age, Chandman site, West Mongolia |
| | Mongol 2 |
| | Mongol 3 |
| | Nr Kiachta; Troiskosavsk / Bargoutine, Dali; Sables de l'Angara; Tounka; Oungwa / Buryats |
| | Nr Kiachta; Troiskosavsk / Northeast Siberia / Buryats |
| | Tuva |
| | Tuvinci |
| | Ulaanbaatar (Urga) / Mongolians |
| | Ulaanbaatar (Urga); Eulojke-Sou Gho; Route du hord d'Ourga; Environs d'Ourga; Ouzga / Mongol |
| | Urga |
| Mainland East Asia | An-Yang |
| | Northern Chinese |
| | Shanghai / Southern China |
| | South Koreans |
| | Tungku; Tientsin / Northern China |
| Malay Archipelago | Aeta tribe, Luzon / Negritos, Philippines |
| | Aeta; Camaunes sud; Dezmosa, Luzon; Bicol de Daraga / Negritos, Philippines |
| | Amboinee |
| | Amboinee, Lesser Sunda |
| | Atjak-Becar; Jambi, Rio Malays; Island of Nias; Mentawei Islands; Billiton Island / Sumatra, Indonesia |
| | Bali |
| | Balinese, Brahmin / Bali, Lesser Sunda |
| | Bisaya; Tagalog; Luzon, Manila; Mindanao; Negros; Panay Island / Philippines |
| | Borneo |
| | Boughis de Sidearing; Amboinais / Celebes |
| | Calatagan, Philippines |
| | Fagal; Bugi; Tagis; Batavia; Madura; Banda / Java |
| | Jalais de Padang; Pontjotsitn de Patjitan: Batavia; Madura; Magelang; Tagis / Java |
| | Jarawa tribe; Great Andaman Island; Little Andaman Island; Port Blair |
| | Java |
| | Koeboe; Nias Island; Billiton Island; Mentawei Islands / Sumatra, Indonesia |
| | Kupang / Timor Islands |
| | Maccassar |
| | Malacca |
| | Malais de Pahang / Malacca |
| | Malay |
| | Malay (Borneo) |
| | Malay Archipelago |
| | Malay Peninsula |
| | Malay Peninsula - Penang |
| | Malay Peninsula, Dullah, Pulo Penana Is. |
| | Malay Peninsula, Johorei |

| | |
|---|---|
| | Maya |
| | Mentawa Is. |
| | Minanese from Island of Sumbawa |
| | Great Andaman Is. |
| | Moro, Island of Sulu |
| | Murut of Batu, Lawi Int, Sarawak / Borneo |
| | Paga Sakai, Gua Cha / Early Malay (Mesolithic to Neolithic) |
| | Pagi-Pagi Islands / Molucca |
| | Pagi-Pagi Islands; Talauer Island; Amboyna Island / Molucca |
| | Philippines 2 |
| | Rebella, Tue e Prei-Drok, Pres de Kampot / Cambodia |
| | Sarawak, Borneo |
| | Selangore |
| | Semang Negritos / Malay, SE Asia |
| | Singapore; Banca Is; Timor Laut; Quedah Fort; Perak; Cebu Is. / Malay |
| | Malaysia / Malay |
| | Solor / Flores island |
| | Sumatra |
| | Philippines |
| | Teressa Island / Nicobar Islands |
| | Timor Island / Lesser Sunda |
| | Toradja / Celebes |
| New Zealand | Chatham Islands Moriori [Moriori / Chatham Islands] |
| | New Zealand Maori [Pihantea, N Island; Whangarei District; Opura / New Zealand, Maori] |
| | Waikato / Maori, New Zealand] |
| North Africa | Guanche, Cave at Lapaz in Puerto Orotava; Puerto de la Madera, Teneriffe / Canary Islands |
| | Libya |
| | Tenerife, Guanche, Canary Islands / Morocco |
| North America | Aleut [Aleutian Island Chain] |
| | CA British Columbia [Klicksiwi River, SE Fort Rupert; Eburne; Nimpkish; Port Hammond; Nootka; Lytton; Bella Bella; Nicola; Namaino; Vancouver Island, Columbia river / British Columbia] |
| | Hammond; Nootka; Lytton; Clayoquot, Vancouver Is.; Bella Bella; Nicola; Nanaimo; Vancouver Island; |
| | CA Canadian Indian [Dene, Forr McPharson; Hare Indians / Canada] |
| | CA Canadian Indian [Ft Rae / Canadian Indians] |
| | CA Northwest Coast [Northwest Coast of Canada] |
| | CA Northwest Territory [Mackenzie Delta, near Fort Mcpherson / Northwest Territory] |
| | Toronto / Ontario] |
| | Simcoe County, Ontario; Lake Huron / Ontario] |
| | CA Tlingit [Admiralty Island; W Coast of Prince of Wales Islands / Tlingit, SE Alaska] |
| | Tlingit / Southeast Alaska] |
| | Eskimo Unknown [Eskimos Locality Unknown] |
| | NA_NatAm [Sioux, Fort Rice / North Dakota] |
| | NEA Chukchi [Chukchis] |
| | Pima |
| | Shell Banks Nr Mobile; Mason Island, Limestone Co; Pine Island Marshall Co; Vackson Co; Baugh's |
| | US Alabama [Shell banks Nr Mobile; Mason Island, Limestone Co; Pine Island, Marshall Co / Alabama] |
| | Indians] |
| | River / Alaska Indians] |

| |
|---|
| US Alaska Indian [Sitka; Yukon River basin / Alaska Indians] |
| US Arch Lake Skull [New Mexico, ca. PA,AAA years BP] |
| Blackfalls; Canyon del Muerto; Canyon de Chelly / Arizona] |
| US Arizona [Nr Allantown, Pueblos; Tonto Apache; Navaho, Canyon Del Muerte; Pima San Carlos, Apache; Cottonwood, Hodbrook; Black Falls; Canyon del Muerto; Canyon de Chelly / Arizona] |
| Miller Co; Nr Snowball, Searcey Co; Pecan Point / Arkansas] |
| Miller Co; Searcey Co; Poinsetle Co / Arkansas] |
| US California [California] |
| US California [North and South California] |
| US California [Santa Clara Co. / California] |
| US Colorado [Ute; Pueblo / Colorado] |
| US Dakota North [Fort Stevenson, Fort Yates : Sioux / North Dakota] |
| US Dakota North [Sioux, Fort Rice / North Dakota] |
| US Dakota North [Sioux, Heart River, Fort Stevenson / North Dakota] |
| US Dakota South [Arikara Indians / South Dakota] |
| US Dakota South [Frot Wadsworth, Mobridge : Arikara Indians / South Dakota] |
| US Dakota South2 [Fort Pierre, Sioux tribe; Mandan Indian, Hart and Knife River / South Dakota] |
| US Dakota SouthP [Arikara; Fort Wadsworth, Mobridge / South Dakota] |
| US Delaware [Lewes / Delaware] |
| US Delaware [Lewes; Fort Resolution Great Slave Lake / Delaware] |
| US Florida [Belle Glade Mound site, Palm Beach County / Florida] |
| US Florida [Old Salt Work; N Symara; Pensacola / Florida] |
| US Georgia [Irene Mound site, Chatham County / Georgia] |
| US Georgia [Irene Mound, Chathan County / Georgia] |
| US Horn Shelter #2 [Texas, ca. 9,AAA years BP] |
| US Illinois [Jersey Co.; Calhoun Co.; St Claire Co.; Henderson Co.; Randolph Co. / Illinois] |
| US Illinois [Jersey Co.; Randolph Co.; Aurora; Calhoun / Illinois] |
| US Illinois [Jersey County / Illinois] |
| US Iowa [Iowa] |
| US Kansas [Coniphan & Riley Co; Wichita / Kansas] |
| US Kansas [Wichita / Kansas] |
| US Kentucky [Indian Knoll, Ohio County / Kentucky] |
| US Kentucky [Ohio County; Union County / Kentucky] |
| US Kentucky [The Indian Knoll, Green River, Ohio County; Union County / Kentucky] |
| US Kodiak [Chief's Point; Kiavak; Old Karluk; Uyak Bay / Kodiak Island] |
| US Kodiak [Shiridan Bay; Uyak Bay; Koniag, Kiavak; Old Kurluk / Kodiak Island] |
| US Kodiak [Uyak Bak, Kodiak Island] |
| US Louisiana [Louisiana] |
| US Louisiana [Monroe / Louisiana] |
| George's Co; Hughes site Montgomery Co; Port Tabacco / Maryland] |
| US Massachusetts [Massachusetts] |
| US Michigan [Chippewa / Michigan] |
| US Mississippi [Chickasaw Indians, Tupelo / Mississippi] |
| US Missouri [Miller's Cave / Missouri] |
| US Missouri [Yankton Sioux Indian, Missouri Valley; Minekaree Indian (Crow Indians) / Missouri] |
| US Montana [Montana] |
| US Montana [Nr Blackfoot / Montana] |
| US Nebraska [Brule Sioux, Bordeaux Creek / Nebraska] |

| | |
|---|---|
| | US Nevada [Pah Ute / Nevada] |
| | US New Jersey [Delaware Indians / New Jersey] |
| | US New Jersey [Delaware Indians, Opposite Minnisink Island / New Jersey] |
| | Caliente; McKinley Co, Zuni; Rincon Del Camino; Kiminioli Valley, Chaco / New Mexico] |
| | US New Mexico [Sandoval Co.; Navaho, Fort Mcrae; Apache, Ojo Caliente; Hawikuh, Mckinley Co.; Pueblo, Puye; McKinley Co, Zuni; Rincon Del Camino,; Kiminioli Valley, Chaco / New Mexico] |
| | US New York [Ulster County; Croton on Hudson / New York] |
| | US Ohio [Hamilton County / Ohio] |
| | US Ohio [Ohio / USA] |
| | US Oklahoma [Ozage / Oklahoma] |
| | US Oregon [Coos Bay / Oregon] |
| | US Oregon [Elk City, Yaguina River, Chetco Indian / Oregon] |
| | US Pennsylvania [Martin's Creek, Northampton County / Pennsylvania] |
| | US Rhode Island [Narragansett / Rhode Island] |
| | US Tennessee [Nr Nashville, Lauderdale County; Cherokee Indian, Pllalchian stock / Tennessee] |
| | US Tennessee [Nr Nashville; Sevierville / Tennessee] |
| | US Texas [Comanche, Ft Conco; Toncaway, Ft. Cobb; Lipan / Texas] |
| | US Texas [Kichopoo; Nr Jackboro; Nr Aat-toh Mountain; Nr Ft Stauton; Ft Concho; Ft Griffin / Texas] |
| | US Utah [Bradshaw Mound, Beaver, Ute; Grand Gulch; Red Canyon / Utah] |
| | US Utah [Ute, Pah-Ute; Grand Gulch; White Canyon / Utah] |
| | Southampton Co.; Fisher site Potomac River, Loudoun Co.; Lynch Station, Campbell Co.; Potomac |
| | US Virginia West [Herriott site / West Virginia] |
| | US Washington [Old Fort, Walla Wallo; Fort Colvilla, Columbia River / Washington State] |
| | US Wisconsin [Ottigamie "Fox Indian" / Wisconsin] |
| | US Wisconsin [Wisconsin] |
| | US Wyoming [Bannock / Wyoming] |
| North of Black Sea | Mesolithic (Fatma-Koba) |
| | Mesolithic (Murzak-Koba) |
| | Uzbekistan; Cossack; Odessa; Lett, Riga; Caucasas, Erivan / Russia |
| | Ukraine Neolithic |
| Northern and Northeastern Europe | Bergen, Trondhjem / Norway |
| | Danish Neolithic |
| | Early Estonia |
| | Insane, Dane / Denmark |
| | Kaberla |
| | Komi |
| | Ladoga Finns |
| | Mangup-Kale, Crimea; Pskoff, Estate Sapolia; Lett, Riga; Caucasas, Erivan / Russia |
| | Reindeer Island |
| | Russians |
| | Saarijarivi, Birkala / Finland |
| | Stockholm / Sweden |
| | Swedish, Iron Age Vallhagar, Gottand |
| | Vallhagar, Gottand / Sweden |
| Northern India | India [Bhdshee, Benares, Agra and Ouah / Northwest Province] |
| | India Assam [Ninu, a Konyak Naga Villege / Assam] |
| | Manipur State / Assam, India] |
| | India Bihar [Ruamgath, Dhalbum; Hyderabad; Patna Province / Behar, India] |

| | |
|---|---|
| | India Northwest [Suttee, Bombay; Delhi / West and Northwest India] |
| | India Northwest Province [Bhdshee, Benares, Agra and Ouah / Northwest Province] |
| Northern South America | Akawoi Indian; Carbouger of Surinam; Macusi Indian; Wapisiana / Guyana |
| | Akawoi Indian; Carbouger, Coronie ; Taruma girl; Arawak tribe; Taruma / Guyana |
| | Bogota; Muizca, Tunjuelo; Muizca: Carare, Choachi / Colombia |
| | Maracaibo / Venezuela |
| | Muizca, Tunjuelo, near Bogola / Colombia |
| | Piaroa Indian, Orinoco River / Venezuela |
| Pacific Islands | Banks Islnad; Royalty Islands; Near Oubatch, Ouebia / New Caledonia |
| | Cape Kujoi, near Rabaul; Gazelle Peninsulla; Ralum; Matupi; Branche / New Britain |
| | Cook Islands [Manuai; Ainu; Mangaia / Cook Islands] |
| | Cook Islands [Manuai; Atiu; Titiaroa; Mangaia / Cook Islands] |
| | Easter Islands [Hotu Itu; Miru Clan; Motu Nui; Orongo / Easter Islands] |
| | Easter Islands [Orongo; Riitoparanu; Miru Clan; Hotu Itu; Motu Nui; Rapa Nui / Easter Islands] |
| | Farawintale, Taravoa, Tou, Papao Fate, Tahiti / Society Islands |
| | Jacquinot Bay / New Britain |
| | Tanga Island, Admiralty / Bismarck Archipelago |
| | Hawaii [Oahu from Mokapu site (precontact), Kauai, Lanai / Hawaii] |
| | Hawaii [Oahu, Kanaka / Hawaii] |
| | Hawaii [Oahu, Lanai, Kauai, Maui, Hawaii, Molokai / Hawaii] |
| | Toriulu : Tonga / Opoulou; Samoa |
| | Levuka, Ovalan; Libouka, Ue Obaiaou, Bourretas tribe; Sebouka; Rotumak Is. / Fiji |
| | Lifuka, Haapai group; Caloline Islands, Oolean, Iouli or Oulleary / Tonga : Samoa |
| | Lihir Island; Admiralty Island, Baluan; St Matthias / New Ireland |
| | Malakula Island, SW Bay, Hahai; Ambrym; Tanna Island; Mallicollo / New Hebrides |
| | Island / New Hebrides |
| | Mapatoni, Tahuata; Hatuatua, Pehipaka, Temoea'oko, Hatiheu, Nuku Hiva; Ua Huka / Marquesas |
| | Mer Island, Hammond Island; Thursday Isl.; Darnley Isl.; Prince of Wales Isl. / Torres Strait |
| | MIC Mariana [Epau / Guam] |
| | MIC Mariana [Guam; Saipan; Tinian / Mariana Islands] |
| | Mokapu |
| | near Oubatche, Ouebia / New Caledonia |
| | New Britain |
| | New Britain (Gazelle Peninsula) |
| | New Britain (Ralum) |
| | New Britain, Child |
| | New Guinea |
| | New Guinea Gulf |
| | New Guinea Gulf Province |
| | New Hebrides |
| | New Hebrides Vanuatu |
| | Vanua Balavu; Rotumak Is.; Ovalau; Nausori, Rewa River, Viti levu; Narocivo, / Fiji |
| | Solomon |
| | Taravoa, Pora, Haapape, Tahiti; Moorea / Society Islands |
| | Tatua, Tabar Island; Simberi; Namatanai; St Matthias site / New Ireland |
| | Temoea'oko, Hatiheu, Fatu Hiva; Tahu Ata; Nuku Hiva; Ua Huka / Marquesas |
| | Torres Strait |
| | Trevanion Island / Santa Cruz Islands |

| | |
|---|---|
| | Uafuka / Marquesas |
| | Malaita Is.; Hammond Isl; Ronbiana, New Georgia / Solomon Islands |
| | W & E Sepik Province, Madang Province, Purari River delta; Fly River Delta / Papua New Guinea |
| | Wallis Is.; Erubor, darnley Is.; Mabuag Is.; Tarnley Is. / Torres Strait |
| | Yap; Palau; Mortlock; Magngahan; Pire Mangohan; Ponape; Lougounor; Fais / Caroline Islands |
| Peninsular India | Bengal district / Northeast India |
| | Bombay / India |
| | Derhampre, Ganjam dist. |
| | Dhalbhum: Bihar / India |
| | Ghazipur, Calcutta; Lascar; Shampooker; Jaunbazar / Bengal, India |
| | Gulf of Cutch, Thug; Dhanko; Rewa Kantha / Bombay, India |
| | locality unknown / India |
| | Malabar / India |
| | Malabar Coast / Southwest India |
| | Santal Chutia, Nagpur / India |
| | Uriah; Akal, Singhbhum / Orissa, India |
| South India | Andhra Pradesh; Vakkaliga / Mysore, India |
| | Ceylon (Malahyalam, Moorman, Singhalese, Tamil, Veddah) |
| | Ceylon, Sri Lanka / Veddah |
| | Mysore / India |
| | Nilgiri, Lahada / Mahalls / India |
| | Sri Lanka / Veddah tribe |
| | Tamil, Colombo / Ceylon / India |
| | Trichinopoly; Trichu, Cochin State; Kandda, Bellary, Dravidian / Madras, India |
| Southeast Asia mainland | Abri-sous-roche de Tam-Tang-Anh / Neolithic Laos |
| | Arracan Hills; Rangoon; Pegu / Myanmar |
| | Bangkok / Thailand |
| | Bronze and Iron age Thailand / Ban Chiang site |
| | Cambodia, Vietnam, Laos, Myanmar, Thailand |
| | Canton; Vic of Ichang; Yunan Prov, Meng-Ting; Shanghai; Hong Kong; Macao / China, South |
| | Dong Dong; Tonkin; Mekong / Vietnam |
| | Kha / Laos |
| | Luang Dra Bang; Kha / Laos |
| | Myanmar |
| | Pho-Binh-Gia, Tonkin / Neolithic Vietnam / including Early Thai from Ban Chiang, Bronze and Iron age |
| | Thailand |
| | Vietnam |
| Southeastern Europe | Croatian, Eso Island, Dalmatia; Serbia; Tuzla, Bosnia / Yugoslavia |
| | Danubian Provinces, Mediaeval Cemetery, Crivoscia (S. Dalmatia) / Herzegovina |
| | EProvince of Otranto; Syracuse, Palermo, Sicily; Rome; Balsorano / Italy |
| | Gipsy, Bucharest / Rumania |
| | Greece |
| | Paterno; Lecce; Popoli; Capistrello Luco; Pontecorvo / Italy |
| | Serbians; Adrianople; Tcirpan / Bulgaria |
| | Skenderum / Albania |
| | Slovenia, Slav; Southern Slav, Dalmatia / Yugoslavia |
| | Tcirpan / Bulgaria |
| | Transylvania; Ploesci, Wallachia; Armenian; Bucharest / Rumania |

| | |
|---|---|
| | Peloponnesus, Kreta, Candia / Greece |
| Southern Africa | Basuto / Lesotho |
| | Bechuanaland, Botswana, Koranna: Hottentot / South Africa |
| | Great Karroo, Beaufort West, South Africa / Bushman |
| | Hottentot, South Africa |
| | Kaffir tribe : Basuto, Mantatee; Tulu; Tambuki / South Africa |
| | Natal Zulu; Nr Rorke's Drift / Zulu, South Africa |
| | South Africa |
| | Pedi |
| | Sands near Wynbeerg; Kalahari / Bushman / South Africa |
| | Mashona; Amexosa / South Africa |
| | Zulu, Natal / South Africa |
| Southern South America | Alacaluf, Port Humbre & Dawson; Cura Cautin; Cllacatuf, Dawson Island; Huilliche, Osorno; Araucanian; Coquimbo; Alacaluf, Fatal Bay, Juan Stuven Island; Quiani, Arica; Calita, Vitor; Wellington Island; Solinas Moud, Arica; Punta Pichalo / Chile/ Chile |
| | Cuchipuy |
| | La Herradura |
| | Paleo-Indian |
| | Patagonia |
| | Punta Teatinos |
| | Beadle Channel, Navarin Island, Yahgan : Tierra del Fuego |
| | Island, Yahgan : Tierra del Fuego |
| Tibetan Plateau and Siwalik Hills | Goorkha from Kalamandu / Nepal |
| | Kashmir / India |
| | Lepcha, Darjeeling, Himalayan slope / Sikkim, India |
| | Lepcha, Darjeeling; Bhotia or Bodpa / Sikkim |
| | Nepal |
| | Pathan; Lahore, Protob Sing; Bajput; Afridi; Jat / Punjab, India |
| | Pokra; Himalaya Slope / Nepal |
| | Province of Kham; Symbunath, Valley of Nepal; Sankhmol / Tibet |
| | Punjab / NW India |
| | Tibetans |
| | Yasinese, Garkuch / Kashmir, India |
| Western Africa | Abuakwa; Dagomba; Kjebi / Ashanti, West Africa |
| | Vai; Kroo tribe / Liberia |
| | Abomey, Dahomey |
| | Abuakwa, Ashanti |
| | Ashanti |
| | Atakpa, Calabar |
| | Bamboo forest, Kigeri Buanda / Uganda : Pygmy / Congo : Akka tribe, Monbuttu, Central Africa / Bamburi tribe, Congo Forest |
| | Calabar |
| | Calabar, Cross River |
| | Coastal West Africa; Sossu; Appa; Lagos; Barconka; Maka; Kounga; Calabar |
| | Coucouleur tribe, Fonta / Senegal |
| | Creek Town, Calabar |
| | Cross River |
| | Dagomba |

| | |
|---|---|
| | Dahomey |
| | Cameroon |
| | Fan, Ogove River; Fernand Vaz River / Gabon |
| | Faradje; Wakema; Tshimbulu, Kasai District; N. Viengo Falls, Partugese / Congo |
| | Fernan Vaz; Loango; Upper Mobanyki; Pygmy; Upongwi County, Nr Selte Cama; Bahuana, Luano / Congo |
| | Free Town / Sierra Leone |
| | Gambia |
| | Guajah / French Guinea, West Africa |
| | Ibea; Mandingo; Boki; Anyang; Duala; Kumabembe; Mabea; Bawa, Yaunde / Cameroon |
| | Ibo |
| | Ibo tirbe / Southern Nigeria |
| | Kalsima Ala, Nigeria, Munshi tribe / Nigeria |
| | Kjebi Ashanti |
| | Kossa or Kasu tribe; Papas or Mahai; Calabar / West Africa |
| | North of Ivory Coast |
| | Ogaja; Kadura; Ayu; Gannawarri from baban Gidda, Kagoro tribe from Modakia / North Nigeria |
| | SSKjebi; Abuakwa; Dagomba; Cannibal from Ononguna / Ashanti, Northern Ghana |
| | Tim Maris Tribe / Sierra Leone |
| | Tolah or Mandingo, Bathurst / Gambia |
| | Whydah, Dahomey |
| Western Europe | Belgium |
| | Dorestad de Heul |
| | Ensay site / Mediaeval Scotland |
| | Merovingian times / France |
| | Lent |
| | Mediaeval England / Repton site |
| | Mediaeval Scotland / Ensay site |
| | Mediaeval Southwest England / Poundbury site |
| | Mediaeval UK / Repton site |
| | Paris; Merovingian to Gallo-Roman Periods / France |
| | Prov. of Gelderland; Amsterdam; Utrecht / Holland |
| | Province of Groningen; Friesland; Province of Gelderland; Amsterdam / Holland |
| | Skenpleby; Barton Hill, Streatley; Long Barrow; Littleton Drew; West Kemmet / Neolithic UK |
| | Spitalfields collection |
| | Rodmarton; Imber, Wilts; Kemmet Wilts; W. Monkton, N. Wilts; Long Barrow Norton Bavant, Wilts; |
| Western South America | Ancon |
| | Asia |
| | Ayalan |
| | Bolivia |
| | Buena Vista |
| | Cabezas Largas |
| | Chanduy |
| | Chavina, Acari Valley |
| | Chicama |
| | Chicama2 |
| | Chilca |
| | Chilca; San Damian; Cerro del Oro; Arica; Aymara; Chincha; Quichua; Balley of the Rimac / Peru |

| |
|---|
| Chimborozo / Ecuador |
| Chincha |
| Cinco Cerros |
| Cinco Cerros / Peru |
| Cinco Cerros; Chilca; Ancon; Cajamarquilla; Lomas; Huarochiri; Chicama; San Damian; Cerro del Oro; Ancom; Quichua Pachacamac; Yunka Crand, Chimee; Canate Valley; Huaco, Callao / Peru |
| Cotocollao |
| Coyungo/a |
| Culebras |
| Ecuador |
| Huaca Prieta |
| Huacho |
| Ica |
| La Cubuya |
| La Paloma |
| Lomas |
| Matucana |
| Nasca region |
| Near Huarochiri |
| Near Santa Lucia |
| Quintai, near Huacho |
| San Damian |
| Santa Elena |
| Sica Sica region, Tama Tam Chullpa; Chelen Chullpa, Churkoni group / Bolivia |
