## Appendix_3 for "DeMoDa: A global open access database for comparative research in human dental morphological variation"

### **Appendix 3. The Dental Morphological Database Scoring Systems** (Martínez de Pinillos González et al., 2026 in preparation)

| TYPE | TOOTH * | SCORING / DEFINITION (Turner et al., 1991) | SCORING / DEFINITION (Scott and Irish, 2017) | Formula for DeMoDa Scoring System 1 | DeMoDa Scoring System 1 (Turner + Scott and Irish) | SCORING / DEFINITION (Hanihara, 2008) | Formula for DeMoDa Scoring System 2 | DeMoDa Scoring System 2 (Turner + Scott and Irish + Hanihara) |
|---|---|---|---|---|---|---|---|---|
| **ACCESSORY CUSPS (Premolar mesial and distal accessory cusps)**<br><br>ACC_CUSP | **$P^3$ $P^4$ (T)**<br>**$P^3$ $P^4$ (S)**<br>**$P^3$ $P^4$ (H)** | 0. No accessory cusps occur.<br>1. Mesial and/or distal accessory cusps are present.<br>(NOTE: for some datasets more detailed morphology was denoted as "M", "D", or "MD". We converted these to Scott and Irish, 2017 scores "1", "2" and "3" respectively, since the DeMoDa committee decided to keep S&I scorings only for this trait.) | 0: accessory cusp absent<br>1: well-defined mesial accessory cusp with palpable cusp tip<br>2: well-defined distal accessory cusp with palpable cusp tip<br>3: well-defined mesial and distal accessory cusps with palpable cusp tips | 0 = 0T / 0S<br>1 = 1T/1S<br>2 = 2S<br>3 = 3S | 0: accessory cusp absent<br>1: well-defined mesial accessory cusp with palpable cusp tip<br>2: well-defined distal accessory cusp with palpable cusp tip<br>3: well-defined mesial and distal accessory cusps with palpable cusp tips | 0: grades 0 of Turner et al., (1991)<br>1: grades 1 of Turner et al., (1991)<br><br>INFO PARA $P^3$ Y $P^4$:<br>*This trait is recorded following the ASU system.* | 0 = 0T / 0S / 0H<br>1 = 1T / 1,2,3S / 1H | 0. Absence: grades 0 of DeMoDa 1<br>1. Present: grades 1 of DeMoDa 1 |
| **ANTERIOR FOVEA**<br><br>AF | **$M_1$ (T)**<br>**$M_1$ $M_2$ $M_3$ (S)** | 0. Anterior fovea is absent. The sulcus between cusps 1 and 2 continues without interruption from the center of the occlusal surface to the mesial border.<br>1. A weak ridge connects the mesial aspects of cusps 1 and 2 producing a faint groove.<br>2. The connecting ridge is larger and the resulting groove deeper than grade 1.<br>3. Groove is longer than in grade 2.<br>4. Groove is very long and mesial ridge is robust. | 0: absence<br>1: trace, with slight development of mesial marginal ridge<br>2: essential ridges on trigonid better developed, as is marginal ridge<br>3: essential ridges pronounced and marginal ridge well developed, producing a distinctive fovea on the anterior portion of the trigonid<br>4: pronounced essential ridges and marginal ridge produce a well-defined fovea | 0 = 0T / 0S<br>1 = 1T / 1S<br>2 = 2T / 2S<br>3 = 3T / 3S<br>4 = 4T / 4S | 0. Anterior fovea is absent.<br>1. A weak ridge connects the mesial aspects of cusps 1 and 2 producing a faint groove.<br>2. The connecting ridge is larger and the resulting groove deeper than grade 1.<br>3. Groove is longer than in grade 2.<br>4. Groove is very long and mesial ridge is robust. | | 0 = 0T / 0S<br>1 = 1T / 1S<br>2 = 2T / 2S<br>3 = 3T / 3S<br>4 = 4T / 4S | 0. Anterior fovea is absent. The sulcus between cusps 1 and 2 continues without interruption from the center of the occlusal surface to the mesial border.<br>1. A weak ridge connects the mesial aspects of cusps 1 and 2 producing a faint groove.<br>2. The connecting ridge is larger and the resulting groove deeper than grade 1.<br>3. Groove is longer than in grade 2.<br>4. Groove is very long and mesial ridge is robust. |
| **CARABELLI'S TRAIT**<br><br>CARA | **$M^2$ $M^3$(T)**<br>**$M^2$ $M^3$ (S)** | 0. The mesiolingual aspect of cusp 1 (protocone) is smooth.<br>1. A groove is present.<br>2. A pit is present.<br>3. A small Y-shaped depression is present.<br>4. A large Y-shaped depression is present.<br>5. A small cusp occurs but it lacks a free apex and its distal border does not contact the lingual groove separating cusps 1 and 4.<br>6. A medium-sized attached-apex cusp is present that makes contact with the medial lingual groove.<br>7. A large free cusp is present.<br>(NOTE: sometimes Turner scored this trait with a number and asterisk. When this happens, means that there is a pit in the lingual groove) | 0: mesiolingual cusp does not exhibit any grooves or pits on the lingual surface<br>1: a vertical groove separates the protocone from the mesial marginal ridge complex; grade 1 expression occurs when there is a slight eminence that deflects distally from this groove<br>2: when expression goes beyond a slight groove or eminence and takes the form of a pit<br>3: expression is still slight but takes on a more distinct form than shown by grades 1 and 2<br>4: the most pronounced expression of Carabelli's trait that does not involve a tubercle with a free apex; grade 4 takes the classic bird-wing form<br>5: small tubercle with a free apex<br>6: moderate tubercle with a free apex<br>7: pronounced tubercle with a free apex | 0 = 0T / 0S<br>1 = 1T / 1S<br>2 = 2T / 2S<br>3 = 3T / 3S<br>4 = 4T / 4S<br>5 = 5T / 5S<br>6 = 6T / 6S<br>7 = 7T / 7S | 0. The mesiolingual aspect of cusp 1 (protocone) is smooth.<br>1. A groove is present.<br>2. A pit is present.<br>3. A small Y-shaped depression is present.<br>4. A large Y-shaped depression is present.<br>5. A small cusp occurs but it lacks a free apex and its distal border does not contact the lingual groove separating cusps 1 and 4.<br>6. A medium-sized attached-apex cusp is present that makes contact with the medial lingual groove.<br>7. A large free cusp is present. | | 0 = 0T / 0S<br>1 = 1T / 1S<br>2 = 2T / 2S<br>3 = 3T / 3S<br>4 = 4T / 4S<br>5 = 5T / 5S<br>6 = 6T / 6S<br>7 = 7T / 7S | 0. The mesiolingual aspect of cusp 1 (protocone) is smooth.<br>1. A groove is present.<br>2. A pit is present.<br>3. A small Y-shaped depression is present.<br>4. A large Y-shaped depression is present.<br>5. A small cusp occurs but it lacks a free apex and its distal border does not contact the lingual groove separating cusps 1 and 4.<br>6. A medium-sized attached-apex cusp is present that makes contact with the medial lingual groove.<br>7. A large free cusp is present. |
| | **$M^1$ (T)**<br>**$M^1$ (S)**<br>**$M^1$ (H)** | 0. The mesiolingual aspect of cusp 1 (protocone) is smooth.<br>1. A groove is present.<br>2. A pit is present.<br>3. A small Y-shaped depression is present.<br>4. A large Y-shaped depression is present.<br>5. A small cusp occurs but it lacks a free apex and its distal border does not contact the lingual groove separating cusps 1 and 4.<br>6. A medium-sized attached-apex cusp is present that makes contact with the medial lingual groove.<br>7. A large free cusp is present.<br>(NOTE: sometimes Turner scored this trait with a number and asterisk. When this happens, means that there is a pit in the lingual groove) | 0: mesiolingual cusp does not exhibit any grooves or pits on the lingual surface<br>1: a vertical groove separates the protocone from the mesial marginal ridge complex; grade 1 expression occurs when there is a slight eminence that deflects distally from this groove<br>2: when expression goes beyond a slight groove or eminence and takes the form of a pit<br>3: expression is still slight but takes on a more distinct form than shown by grades 1 and 2<br>4: the most pronounced expression of Carabelli's trait that does not involve a tubercle with a free apex; grade 4 takes the classic bird-wing form<br>5: small tubercle with a free apex<br>6: moderate tubercle with a free apex<br>7: pronounced tubercle with a free apex | 0 = 0T / 0S<br>1 = 1T / 1S<br>2 = 2T / 2S<br>3 = 3T / 3S<br>4 = 4T / 4S<br>5 = 5T / 5S<br>6 = 6T / 6S<br>7 = 7T / 7S | 0. The mesiolingual aspect of cusp 1 (protocone) is smooth.<br>1. A groove is present.<br>2. A pit is present.<br>3. A small Y-shaped depression is present.<br>4. A large Y-shaped depression is present.<br>5. A small cusp occurs but it lacks a free apex and its distal border does not contact the lingual groove separating cusps 1 and 4.<br>6. A medium-sized attached-apex cusp is present that makes contact with the medial lingual groove.<br>7. A large free cusp is present. | 0: grades 0-2 of Turner et al., (1991)<br>1: grades 3-7 of Turner et al., (1991)<br><br>INFO PARA $M^1$:<br>*Based on Dahlberg's (1963) criteria, presence (Dahlberg's d‑g) is distinguished from absence (Dahlberg's a‑c). The grade a‑c is equivalent to 0‑2 of the ASU system.* | 0 = 0,1,2T / 0,1,2S / 0H<br>1 = 3,4,5,6,7T / 3,4,5,6,7S / 1H | 0. Absence: grades 0-2 of DeMoDa 1<br>1. Present: grades 3-7 of DeMoDa 1 |
| **CENTRAL RIDGE**<br><br>CR | **$P_3$ (H)** | | | | | 0: the sagittal fissure is continuous.<br>1: the sagittal fissure is interrupted by a crest.<br><br>INFO PARA $P_3$:<br>*Deutero-proto relationship. The scoring procedure followed is one by Higa et al. (2003).* | 0 = 0H<br>1 = 1H | 0. Absence: the sagittal fissure is continuous.<br>1. Present: the sagittal fissure is interrupted by a crest. |
| **CONGENITAL ABSENCE**<br><br>CONG_AB | **$I^2$ $P^4$ $M^3$ (T)**<br>**$I_1$ $P_4$ $M_3$ (T)** | 0. Tooth is present. Any degree of visible impaction is considered as present.<br>1. Tooth is congenitally absent. No sign of tooth.<br>(NOTE: Turner often used a different scoring system that included the "+" sign. We have left them in the database, but | | 0 = 0T<br>1 = 1T / 2S peg-shaped incisor (upper I2 variants) / 3S peg-shaped molar (upper M3) | 0. Tooth is present. Any degree of visible impaction is considered as present.<br>1. Tooth is congenitally absent. No sign of tooth. | | 0 = 0T<br>1 = 1T / 2S peg-shaped incisor (upper I2 variants) / 3S peg-shaped molar (upper M3) | 0. Tooth is present. Any degree of visible impaction is considered as present.<br>1. Tooth is congenitally absent. No sign of tooth. |

| | | | | | | | | |
|---|---|---|---|---|---|---|---|---|
| | | we don't know what they mean, although they probably refer to "1" as he did not use that number on the templates) | | | | | | |
| **CUSP 5 (Metaconule)**<br><br>C5M | **$M^1 M^2 M^3$ (T)**<br>**$M^1 M^2 M^3$ (S)** | 0. Site of cusp 5 is smooth, there being only a single distal groove present separating cusp 3 and 4.<br>1. Faint cuspule present.<br>2. Trace cuspule present.<br>3. Small cuspule present.<br>4. Small cusp present.<br>5. Medium-sized cusp present. | 0: trait is absent, only one vertical groove on distal surface of upper molar between hypocone and metacone<br>1: slight conule<br>2: trace conule<br>3: small cuspule<br>4: small cusp<br>5: medium cusp | 0 = 0T / 0S<br>1 = 1T / 1S<br>2 = 2T / 2S<br>3 = 3T / 3S<br>4 = 4T / 4S<br>5 = 5T / 5S | 0. Site of cusp 5 is smooth, there being only a single distal groove present separating cusps 3 and 4.<br>1. Faint cuspule present.<br>2. Trace cuspule present.<br>3. Small cuspule present.<br>4. Small cusp present.<br>5. Medium-sized cusp present. | | 0 = 0T / 0S<br>1 = 1T / 1S<br>2 = 2T / 2S<br>3 = 3T / 3S<br>4 = 4T / 4S<br>5 = 5T / 5S | 0. Site of cusp 5 is smooth, there being only a single distal groove present separating cusps 3 and 4.<br>1. Faint cuspule present.<br>2. Trace cuspule present.<br>3. Small cuspule present.<br>4. Small cusp present.<br>5. Medium-sized cusp present. |
| **CUSP 5 (Hypoconulid)**<br><br>C5H | **$M_1 M_3$ (T)**<br>**$M_1 M_3$ (S)** | 0. No occurrence of cusp 5. The molar has only four cusps (cusps 1-4).<br>1. Cusp 5 is present and very small.<br>2. Cusp 5 is small.<br>3. Cusp 5 is medium-sized.<br>4. Cusp 5 is large.<br>5. Cusp 5 is very large. | 0: hypoconulid is absent (four-cusped tooth)<br>1: trace expression<br>2: slight<br>3: moderate<br>4: strong<br>5: pronounced | 0 = 0T / 0S<br>1 = 1T / 1S<br>2 = 2T / 2S<br>3 = 3T / 3S<br>4 = 4T / 4S<br>5 = 5T / 5S | 0. No occurrence of cusp 5.<br>1. Cusp 5 is present and very small.<br>2. Cusp 5 is small.<br>3. Cusp 5 is medium-sized.<br>4. Cusp 5 is large.<br>5. Cusp 5 is very large. | | 0 = 0T / 0S<br>1 = 1T / 1S<br>2 = 2T / 2S<br>3 = 3T / 3S<br>4 = 4T / 4S<br>5 = 5T / 5S | 0. No occurrence of cusp 5.<br>1. Cusp 5 is present and very small.<br>2. Cusp 5 is small.<br>3. Cusp 5 is medium-sized.<br>4. Cusp 5 is large.<br>5. Cusp 5 is very large. |
| | **$M_2$ (T)**<br>**$M_2$ (S)**<br>**$M_2$ (H)** | 0. No occurrence of cusp 5. The molar has only four cusps (cusps 1-4).<br>1. Cusp 5 is present and very small.<br>2. Cusp 5 is small.<br>3. Cusp 5 is medium-sized.<br>4. Cusp 5 is large.<br>5. Cusp 5 is very large. | 0: hypoconulid is absent (four-cusped tooth)<br>1: trace expression<br>2: slight<br>3: moderate<br>4: strong<br>5: pronounced | 0 = 0T / 0S<br>1 = 1T / 1S<br>2 = 2T / 2S<br>3 = 3T / 3S<br>4 = 4T / 4S<br>5 = 5T / 5S | 0. No occurrence of cusp 5. The molar has only four cusps (cusps 1-4).<br>1. Cusp 5 is present and very small.<br>2. Cusp 5 is small.<br>3. Cusp 5 is medium-sized.<br>4. Cusp 5 is large.<br>5. Cusp 5 is very large. | 0: grade 0 of Turner et al., (1991)<br>1: grades 1-5 of Turner et al., (1991)<br><br>INFO PARA $M_2$:<br>*The counting method is the same as that of the ASU system.* | 0 = 0T / 0S /0H<br>1 = 1,2,3,4,5T / 1,2,3,4,5S / 1H | 0. Absence: grades 0 of DeMoDa 1<br>1. Present: grades 1-5 of DeMoDa 1 |
| **CUSP 6**<br><br>C6 | **$M_3$ (T)**<br>**$M_3$ (S)** | 0. Cusp 6 is absent.<br>1. Cusp 6 is much smaller than cusp 5.<br>2. Cusp 6 is smaller than cusp 5.<br>3. Cusp 6 is equal in size to cusp 5.<br>4. Cusp 6 is larger than cusp 5.<br>5. Cusp 6 is much larger than cusp 6. | 0: absence of cusp 6<br>1: cusp 5 is more than twice the size of cusp 6<br>2: cusp 5 is about twice as large as cusp 6<br>3: cusps 5 and 6 are about equal in size<br>4: cusp 6 is slightly larger than cusp 5<br>5: cusp 6 is markedly larger than cusp 5 | 0 = 0T / 0S<br>1 = 1T / 1S<br>2 = 2T / 2S<br>3 = 3T / 3S<br>4 = 4T / 4S<br>5 = 5T / 5S | 0. Cusp 6 is absent.<br>1. Cusp 6 is much smaller than cusp 5.<br>2. Cusp 6 is smaller than cusp 5.<br>3. Cusp 6 is equal in size to cusp 5.<br>4. Cusp 6 is larger than cusp 5.<br>5. Cusp 6 is much larger than cusp 6. | | 0 = 0T / 0S<br>1 = 1T / 1S<br>2 = 2T / 2S<br>3 = 3T / 3S<br>4 = 4T / 4S<br>5 = 5T / 5S | 0. Cusp 6 is absent.<br>1. Cusp 6 is much smaller than cusp 5.<br>2. Cusp 6 is smaller than cusp 5.<br>3. Cusp 6 is equal in size to cusp 5.<br>4. Cusp 6 is larger than cusp 5.<br>5. Cusp 6 is much larger than cusp 6. |
| | **$M_1 M_2$ (T)**<br>**$M_1 M_2$ (S)**<br>**$M_1 M_2$ (H)** | 0. Cusp 6 is absent.<br>1. Cusp 6 is much smaller than cusp 5.<br>2. Cusp 6 is smaller than cusp 5.<br>3. Cusp 6 is equal in size to cusp 5.<br>4. Cusp 6 is larger than cusp 5.<br>5. Cusp 6 is much larger than cusp 6. | 0: absence of cusp 6<br>1: cusp 5 is more than twice the size of cusp 6<br>2: cusp 5 is about twice as large as cusp 6<br>3: cusps 5 and 6 are about equal in size<br>4: cusp 6 is slightly larger than cusp 5<br>5: cusp 6 is markedly larger than cusp 5 | 0 = 0T / 0S<br>1 = 1T / 1S<br>2 = 2T / 2S<br>3 = 3T / 3S<br>4 = 4T / 4S<br>5 = 5T / 5S | 0. Cusp 6 is absent.<br>1. Cusp 6 is much smaller than cusp 5.<br>2. Cusp 6 is smaller than cusp 5.<br>3. Cusp 6 is equal in size to cusp 5.<br>4. Cusp 6 is larger than cusp 5.<br>5. Cusp 6 is much larger than cusp 6. | 0: grade 0 of Turner et al., (1991)<br>1: grades 1-5 of Turner et al., (1991)<br><br>INFO PARA $M_1$:<br>*The counting method of this trait is the same as that of the ASU system. Scored cusp as present regardless of the size.*<br><br>INFO PARA $M_2$:<br>*Follows the ASU system.* | 0 = 0T / 0S /0H<br>1 = 1,2,3,4,5T / 1,2,3,4,5S / 1H | 0. Absence: grades 0 of DeMoDa 1<br>1. Present: grades 1-5 of DeMoDa 1 |
| **CUSP 7**<br><br>C7 | **$M_2 M_3$ (T)**<br>**$M_2 M_3$ (S)** | 0. No occurrence of cusp 7.<br>1. Faint cusp is present. There are two weak lingual grooves also present instead of one.<br>1A. A faint tipless cusp 7 occurs displaced as a bulge on the lingual surface of cusp.<br>2. Cusp 7 is small.<br>3. Cusp 7 is medium-sized.<br>4. Cusp 7 is large.<br>(NOTE: some Turner datasheets scored this feature with ">4". We don't know what they mean, but they probably indicate the cusps could be bigger than large) | 0: no accessory cusp between cusps 2 and 4<br>1: small, wedge-shaped cusp between cusps 2 and 4<br>1A: this expression does not assume the typical wedge-shaped form of a cusp 7 but is marked by a groove on the lingual surface of the metaconid<br>2: distinct but small cusp<br>3: moderate cusp<br>4: large cusp | 0 = 0T / 0S<br>1 = 1T / 1S<br>1A = 1A T / 1A S<br>2 = 2T / 2S<br>3 = 3T / 3S<br>4 = 4T / 4S | 0. No occurrence of cusp 7.<br>1. Faint cusp is present. There are two weak lingual grooves also present instead of one.<br>1A. A faint tipless cusp 7 occurs displaced as a bulge on the lingual surface of cusp.<br>2. Cusp 7 is small.<br>3. Cusp 7 is medium-sized.<br>4. Cusp 7 is large. | | 0 = 0T / 0S<br>1 = 1T / 1S<br>1A = 1A T / 1A S<br>2 = 2T / 2S<br>3 = 3T / 3S<br>4 = 4T / 4S | 0. No occurrence of cusp 7.<br>1. Faint cusp is present. There are two weak lingual grooves also present instead of one.<br>1A. A faint tipless cusp 7 occurs displaced as a bulge on the lingual surface of cusp.<br>2. Cusp 7 is small.<br>3. Cusp 7 is medium-sized.<br>4. Cusp 7 is large. |
| | **$M_1$ (T)**<br>**$M_1$ (S)**<br>**$M_1$ (H)** | 0. No occurrence of cusp 7.<br>1. Faint cusp is present. There are two weak lingual grooves also present instead of one.<br>1A. A faint tipless cusp 7 occurs displaced as a bulge on the lingual surface of cusp.<br>2. Cusp 7 is small.<br>3. Cusp 7 is medium-sized.<br>4. Cusp 7 is large.<br>(NOTE: some Turner datasheets scored this feature with ">4". We don't know what they mean, but they probably indicate the cusps could be bigger than large) | 0: no accessory cusp between cusps 2 and 4<br>1: small, wedge-shaped cusp between cusps 2 and 4<br>1A: this expression does not assume the typical wedge-shaped form of a cusp 7 but is marked by a groove on the lingual surface of the metaconid<br>2: distinct but small cusp<br>3: moderate cusp<br>4: large cusp | 0 = 0T / 0S<br>1 = 1T / 1S<br>1A = 1A T / 1A S<br>2 = 2T / 2S<br>3 = 3T / 3S<br>4 = 4T / 4S | 0. No occurrence of cusp 7.<br>1. Faint cusp is present. There are two weak lingual grooves also present instead of one.<br>1A. A faint tipless cusp 7 occurs displaced as a bulge on the lingual surface of cusp.<br>2. Cusp 7 is small.<br>3. Cusp 7 is medium-sized.<br>4. Cusp 7 is large. | 0: grade 0 of Turner et al., (1991)<br>1: grades 1-4 of Turner et al., (1991)<br><br>INFO PARA $M_1$:<br>*The counting method of this trait is the same as that of the ASU system. Scored cusp as present regardless of the size.* | 0 = 0T / 0S /0H<br>1 = 1,1A,2,3,4T / 1,1A,2,3,4S / 1H | 0. Absence: grades 0 of DeMoDa 1<br>1. Present: grades 1-4 of DeMoDa 1 |
| **CUSP NUMBER**<br><br>CUSPN | **$M_1 M_2 M_3$ (T)**<br>**$M_1 M_2 M_3$ (S)** | 4. Cusps 1-4 are present (1, protoconid; 2, metaconid: 3, hypoconid; 4, entoconid).<br>5. Cusps 1-5 are present (5, hypoconulid).<br>6. Cusps 1-6 are present (6, entoconulid).<br>(NOTE: some Turner datasheets scored this feature with "3" and ">4". We don't know what they mean, but they probably indicate the number of cusps in those particular individuals) | 4. Cusps 1-4 are present (1, protoconid; 2, metaconid: 3, hypoconid; 4, entoconid).<br>5. Cusps 1-5 are present (5, hypoconulid).<br>6. Cusps 1-6 are present (6, entoconulid). | 4 = 4T / 4S<br>5 = 5T / 5S<br>6 = 6T / 6S | 4. Cusps 1-4 are present (1, protoconid; 2, metaconid: 3, hypoconid; 4, entoconid).<br>5. Cusps 1-5 are present (5, hypoconulid).<br>6. Cusps 1-6 are present (6, entoconulid). | | 4 = 4T / 4S<br>5 = 5T / 5S<br>6 = 6T / 6S | 4. Cusps 1-4 are present (1, protoconid; 2, metaconid: 3, hypoconid; 4, entoconid).<br>5. Cusps 1-5 are present (5, hypoconulid).<br>6. Cusps 1-6 are present (6, entoconulid). |

| | | | | | | | | |
|---|---|---|---|---|---|---|---|---|
| **DEFLECTING WRINKLE**<br><br>DW | **$M_2$ $M_3$ (T)**<br>**$M_2$ $M_3$ (S)** | 0. Deflecting wrinkle is absent. Medial ridge of cusp 2 is straight.<br>1. Cusp 2 medial ridge is straight but shows a midpoint constriction.<br>2. Medial ridge is deflected distalward, but does not make contact with cusp 4.<br>3. Medial ridge is strongly deflected distalward, forming an L-shaped ridge. The medial ridge contacts cusp 4.<br>(NOTE: although this trait seldom occurs on $M_2$ and $M_3$, we have decided to include it for those teeth) | 0: deflecting wrinkle absent; essential ridge of metaconid runs a straight course from cusp tip to central occlusal fossa<br>1: essential ridge is straight but with midpoint constriction<br>2: essential ridge deflects at halfway point toward central occlusal fossa but does not contact hypoconid<br>3: essential ridge shows strong deflection at midpoint and does contact hypoconid | 0 = 0T / 0S<br>1 = 1T / 1S<br>2 = 2T / 2S<br>3 = 3T / 3S | 0: deflecting wrinkle absent; essential ridge of metaconid runs a straight course from cusp tip to central occlusal fossa<br>1: essential ridge is straight but with midpoint constriction<br>2: essential ridge deflects at halfway point toward central occlusal fossa but does not contact hypoconid<br>3: essential ridge shows strong deflection at midpoint and does contact hypoconid | | 0 = 0T / 0S<br>1 = 1T / 1S<br>2 = 2T / 2S<br>3 = 3T / 3S | 0: deflecting wrinkle absent; essential ridge of metaconid runs a straight course from cusp tip to central occlusal fossa<br>1: essential ridge is straight but with midpoint constriction<br>2: essential ridge deflects at halfway point toward central occlusal fossa but does not contact hypoconid<br>3: essential ridge shows strong deflection at midpoint and does contact hypoconid |
| | **$M_1$ (T)**<br>**$M_1$ (S)**<br>**$M_1$ (H)** | 0. Deflecting wrinkle is absent. Medial ridge of cusp 2 is straight.<br>1. Cusp 2 medial ridge is straight but shows a midpoint constriction.<br>2. Medial ridge is deflected distalward, but does not make contact with cusp 4.<br>3. Medial ridge is strongly deflected distalward, forming an L-shaped ridge. The medial ridge contacts cusp 4. | 0: deflecting wrinkle absent; essential ridge of metaconid runs a straight course from cusp tip to central occlusal fossa<br>1: essential ridge is straight but with midpoint constriction<br>2: essential ridge deflects at halfway point toward central occlusal fossa but does not contact hypoconid<br>3: essential ridge shows strong deflection at midpoint and does contact hypoconid | 0 = 0T / 0S<br>1 = 1T / 1S<br>2 = 2T / 2S<br>3 = 3T / 3S | 0: deflecting wrinkle absent; essential ridge of metaconid runs a straight course from cusp tip to central occlusal fossa<br>1: essential ridge is straight but with midpoint constriction<br>2: essential ridge deflects at halfway point toward central occlusal fossa but does not contact hypoconid<br>3: essential ridge shows strong deflection at midpoint and does contact hypoconid | 0: grades 0-1 of Turner et al., (1991)<br>1: grades 2-3 of Turner et al., (1991)<br><br>INFO PARA $M_1$:<br>*Following Weidenreich (1937), if the medial ridge of metaconid deflected distally, the trait is scored as present. Presence corresponds to grade 2‑3 of the ASU system.* | 0 = 0-1T / 0-1S / 0H<br>1 = 2,3T / 2,3S / 1H | 0. Absence: grades 0-1 of DeMoDa 1<br>1. Present: grades 2-3 of DeMoDa 1 |
| **DISTAL ACCESSORY RIDGE (Canine distal accessory ridge) DAR**<br><br>DAR | **C’ (T)**<br>**C, (T)**<br>**C’ (S)**<br>**C, (S)** | 0. Distal accessory ridge is absent.<br>1. Distal accessory ridge is very faint. No example is provided on plaque.<br>2. Distal accessory ridge is weakly developed.<br>3. Distal accessory ridge is moderately developed.<br>4. Distal accessory ridge is strongly developed.<br>5. Distal accessory ridge is very pronounced. | 0: trait absence<br>1: faint expression (not shown on UC DAR plaque)<br>2: slight expression<br>3: moderate development<br>4: strongly developed<br>5: pronounced expression | 0 = 0T / 0S<br>1 = 1T / 1S<br>2 = 2T / 2S<br>3 = 3T / 3S<br>4 = 4T / 4S<br>5 = 5T / 5S | 0: absent<br>1: faint expression (not shown on UC DAR plaque)<br>2: slight expression<br>3: moderate development<br>4: strongly developed<br>5: pronounced expression | | 0 = 0T / 0S<br>1 = 1T / 1S<br>2 = 2T / 2S<br>3 = 3T / 3S<br>4 = 4T / 4S<br>5 = 5T / 5S | 0: absent<br>1: faint expression (not shown on UC DAR plaque)<br>2: slight expression<br>3: moderate development<br>4: strongly developed<br>5: pronounced expression |
| **DISTAL TRIGONID CREST**<br><br>DTC | **$M_2$ $M_3$ (T)**<br>**$M_2$ $M_3$ (S)** | 0. Absent. Distal borders of cusps 1 and 2 are not attached by a crest or loph.<br>1. Present. Distal borders are connected by a ridge.<br>(NOTE: Turner often used a different scoring system that included the “+” sign. We have left them in the database, but we don’t know what they mean, although they probably refer to “1” as he did not use that number on the templates) | 0: distal trigonid crest absent<br>1: distal trigonid crest present | 0 = 0T / 0S<br>1 = 1T / 1S | 0: distal trigonid crest absent<br>1: distal trigonid crest present | | 0 = 0T / 0S<br>1 = 1T / 1S | 0: distal trigonid crest absent<br>1: distal trigonid crest present |
| | **$M_1$ (T)**<br>**$M_1$ (S)**<br>**$M_1$ (H)** | 0. Absent. Distal borders of cusps 1 and 2 are not attached by a crest or loph.<br>1. Present. Distal borders are connected by a ridge.<br>(NOTE: Turner often used a different scoring system that included the “+” sign. We have left them in the database, but we don’t know what they mean, although they probably refer to “1” as he did not use that number on the templates) | 0: distal trigonid crest absent<br>1: distal trigonid crest present | 0 = 0T / 0S<br>1 = 1T / 1S | 0: distal trigonid crest absent<br>1: distal trigonid crest present | 0: grade 0 of Turner et al., (1991)<br>1: grade 1 of Turner et al., (1991)<br><br>INFO PARA $M_1$:<br>*Following the original description by Weidenreich (1937), the crest connecting the tip of the metaconid with the distal accessory ridge of the protoconid without interruption is scored as present.* | 0 = 0T / 0S / 0H<br>1 = 1T / 1S / 1H | 0. Absence: grades 0 of DeMoDa 1<br>1. Present: grades 1 of DeMoDa 1 |
| **DISTOSAGITTAL RIDGE (Uto-Aztecan premolar)**<br><br>DSR | **$P^3$ (T)**<br>**$P^3$ (S)** | 0. Normal premolar form occurs.<br>1. Distosagittal ridge is present. | Grade 0: absent<br>Grade 1: present | 0 = 0T / 0S<br>1 = 1T / 1S | 0: absent<br>1: present | | 0 = 0T / 0S<br>1 = 1T / 1S | 0: absent<br>1: present |
| **DOUBLE-SHOVELING**<br><br>DSHOV | **$I^2$ C’ $P^3$ (T)**<br>**$I_1$ $I_2$ (T)**<br>**$I^2$ (S)** | 0. Labial surface is smooth.<br>1. Faint mesial and distal ridging can be seen in strong contrasting light. Distal ridge maybe absent in this and stronger grades.<br>2. Trace ridging.<br>3. Semi-double-shovel. Ridging can be readily palpated.<br>4. Double-shovel. Ridging is pronounced on at least 1/2 of total crown length.<br>5. Pronounced double-shovel. Ridging is very prominent and may occur from the occlusal surface to the crown-root junction.<br>6. Double-shoveling also occurs on the upper lateral incisors, canines, first premolars and lower incisors. | 0 (absence): no labial marginal ridges present; surface is smooth<br>1 (faint): very faint labial ridging, more evident on mesial than distal margin<br>2 (trace): ridge more distinct than faint expression of grade 1 but still slight<br>3 (slight): ridges distinct enough to be palpated<br>4 (moderate): ridging clearly evident along at least one half of crown height<br>5 (pronounced): very distinct ridging expressed from incisal edge to crown root junction<br>6: (very pronounced): extreme double-shoveling with well-developed ridges along both the mesial and distal labial margins | 0 = 0T / 0S<br>1 = 1T / 1S<br>2 = 2T / 2S<br>3 = 3T / 3S<br>4 = 4T / 4S<br>5 = 5T / 5S<br>6 = 6T / 6S | 0 (absence): no labial marginal ridges present; surface is smooth<br>1 (faint): very faint labial ridging, more evident on mesial than distal margin<br>2 (trace): ridge more distinct than faint expression of grade 1 but still slight<br>3 (slight): ridges distinct enough to be palpated<br>4 (moderate): ridging clearly evident along at least one half of crown height<br>5 (pronounced): very distinct ridging expressed from incisal edge to crown root junction<br>6: (very pronounced): extreme double-shoveling with well-developed ridges along both the mesial and distal labial margins | | 0 = 0T / 0S<br>1 = 1T / 1S<br>2 = 2T / 2S<br>3 = 3T / 3S<br>4 = 4T / 4S<br>5 = 5T / 5S<br>6 = 6T / 6S | 0 (absence): no labial marginal ridges present; surface is smooth<br>1 (faint): very faint labial ridging, more evident on mesial than distal margin<br>2 (trace): ridge more distinct than faint expression of grade 1 but still slight<br>3 (slight): ridges distinct enough to be palpated<br>4 (moderate): ridging clearly evident along at least one half of crown height<br>5 (pronounced): very distinct ridging expressed from incisal edge to crown root junction<br>6: (very pronounced): extreme double-shoveling with well-developed ridges along both the mesial and distal labial margins |
| | **$I^1$ (T)**<br>**$I^1$ (S)**<br>**$I^1$ (H)** | 0. Labial surface is smooth.<br>1. Faint mesial and distal ridging can be seen in strong contrasting light. Distal ridge maybe absent in this and stronger grades.<br>2. Trace ridging.<br>3. Semi-double-shovel. Ridging can be readily palpated. | 0 (absence): no labial marginal ridges present; surface is smooth<br>1 (faint): very faint labial ridging, more evident on mesial than distal margin<br>2 (trace): ridge more distinct than faint expression of grade 1 but still slight<br>3 (slight): ridges distinct enough to be palpated | 0 = 0T / 0S<br>1 = 1T / 1S<br>2 = 2T / 2S<br>3 = 3T / 3S<br>4 = 4T / 4S<br>5 = 5T / 5S<br>6 = 6T / 6S | 0 (absence): no labial marginal ridges present; surface is smooth<br>1 (faint): very faint labial ridging, more evident on mesial than distal margin<br>2 (trace): ridge more distinct than faint expression of grade 1 but still slight<br>3 (slight): ridges distinct enough to be palpated | 0: grades 0-2 of Turner et al., (1991)<br>1: grades 3-6 of Turner et al., (1991)<br><br>INFO PARA $I^1$:<br>*Following the ASU system, the grade 3‑6 is scored as present.* | 0 = 0,1,2T / 0,1,2S / 0H<br>1 = 3,4,5,6T / 3,4,5,6S / 1H | 0. Absence: grades 0-2 of DeMoDa 1<br>1. Present: grades 3-6 of DeMoDa 1 |

| | | | | | | | | |
|---|---|---|---|---|---|---|---|---|
| | | 4. Double-shovel. Ridging is pronounced on at least 1/2 of total crown length.<br>5. Pronounced double-shovel. Ridging is very prominent and may occur from the occlusal surface to the crown-root junction.<br>6. Double-shoveling also occurs on the upper lateral incisors, canines, first premolars and lower incisors. | 4 (moderate): ridging clearly evident along at least one half of crown height<br>5 (pronounced): very distinct ridging expressed from incisal edge to crown root junction<br>6: (very pronounced): extreme double-shoveling with well-developed ridges along both the mesial and distal labial margins | | 4 (moderate): ridging clearly evident along at least one half of crown height<br>5 (pronounced): very distinct ridging expressed from incisal edge to crown root junction<br>6: (very pronounced): extreme double-shoveling with well-developed ridges along both the mesial and distal labial margins | | | |
| **ENAMEL EXTENSIONS**<br><br>EE | **$P^3$ $P^4$ (T)**<br>**$M^1$ $M^2$ $M^3$ (T)**<br>**$M^1$ $M^2$ $M^3$ (S)**<br>**$M_1$ $M_2$ $M_3$ (S)** | 0. Enamel border is straight or rarely curved towards the crown. Score any extension not attached to crown as absent.<br>1. A faint, approximately 1.0 mm long enamel extension projects towards and along the root.<br>2. A medium-sized, approximately 2.0 mm long enamel extension.<br>3. A lengthy extension, generally greater than 4.0 mm in length is present. It may extend all the way to the root bifurcation on molar teeth. | 0: cervical enamel line is horizontal<br>1: enamel line extends about 1 mm toward root bifurcation<br>2: enamel line extends about 2 mm toward root bifurcation<br>3: enamel line extends 4 mm or more toward root bifurcation | 0 = 0T / 0S<br>1 = 1T / 1S<br>2 = 2T / 2S<br>3 = 3T / 3S | 0: cervical enamel line is horizontal<br>1: enamel line extends about 1 mm toward root bifurcation<br>2: enamel line extends about 2 mm toward root bifurcation<br>3: enamel line extends 4 mm or more toward root bifurcation | | 0 = 0T / 0S<br>1 = 1T / 1S<br>2 = 2T / 2S<br>3 = 3T / 3S | 0: cervical enamel line is horizontal<br>1: enamel line extends about 1 mm toward root bifurcation<br>2: enamel line extends about 2 mm toward root bifurcation<br>3: enamel line extends 4 mm or more toward root bifurcation |
| **GROOVE PATTERN**<br><br>GP | **$M_1$ $M_2$ $M_3$ (T)**<br>**$M_1$ $M_2$ $M_3$ (S)** | Y. Cusps 2 and 3 are in contact.<br>X. Cusps 1 and 4 are in contact.<br>+. Cusps 1 - 4 are in contact. | Y pattern: contact between cusps 2 and 3<br>X pattern: contact between cusps 1 and 4<br>+ pattern: contact between cusps 1, 2, 3, and 4 at central sulcus | Y = YT / YS<br>X = XT / XS<br>+ = +T / +S | Y. Cusps 2 and 3 are in contact.<br>X. Cusps 1 and 4 are in contact.<br>+. Cusps 1- 4 are in contact. | | Y = YT / YS<br>X = XT / XS<br>+ = +T / +S | Y. Cusps 2 and 3 are in contact.<br>X. Cusps 1 and 4 are in contact.<br>+. Cusps 1- 4 are in contact. |
| **HYPOCONE**<br><br>HYPO | **$M^1$ $M^3$ (T)**<br>**$M^1$ $M^3$ (S)** | 0. No hypocone (cusp 4). Site is smooth.<br>1. Faint ridging present.<br>2. Faint cuspule present<br>3. Small cusp present.<br>3.5. Moderate-sized cusp present.<br>4. Large cusp present.<br>5. Very large cusp present. | 0: no hypocone expression of any form; a true three-cusped tooth.<br>1: for this grade, there is a low-level expression of the hypocone, often expressed as no more than an outline on the distolingual aspect of the trigon. In Dahlberg's original classification, this would be scored as a three-cusped upper molar along with grade 0.<br>2: in the Dahlberg classification, 3+ was equivalent to a small conical hypocone on the distolingual border of the trigon; grade 2 reflects this phenotype, where there is basically a conical cusp, or tubercle, with a free apex.<br>3: the hypocone is reduced in size but assumes a normal ovate shape along with a distinct free apex.<br>4: this grade would be equivalent to 3.5 on the modified hypocone plaque; the hypocone is reduced in size but is moderate rather than slight in expression.<br>5: hypocone is well developed, a step beyond grade 4.<br>6: pronounced expression of the hypocone; often equals or exceeds the size of the major cusps of the trigon. | 0 = 0T / 0S<br>1 = 1T / 1S<br>2 = 2T / 2S<br>3 = 3T / 3S<br>4 = 3.5T / 4S<br>5 = 4T / 5S<br>6 = 5T / 6S | 0. No hypocone.<br>1. Faint ridging present.<br>2. Faint cuspule present.<br>3. Small cusp present.<br>4. Moderate-sized cusp present.<br>5. Large cusp present.<br>6. Very large cusp present. | | 0 = 0T / 0S<br>1 = 1T / 1S<br>2 = 2T / 2S<br>3 = 3T / 3S<br>4 = 3.5T / 4S<br>5 = 4T / 5S<br>6 = 5T / 6S | 0. No hypocone.<br>1. Faint ridging present.<br>2. Faint cuspule present<br>3. Small cusp present.<br>4. Moderate-sized cusp present.<br>5. Large cusp present.<br>6. Very large cusp present. |
| | **$M^2$ (T)**<br>**$M^2$ (S)**<br>**$M^2$ (H)** | 0. No hypocone (cusp 4). Site is smooth.<br>1. Faint ridging present.<br>2. Faint cuspule present<br>3. Small cusp present.<br>3.5. Moderate-sized cusp present.<br>4. Large cusp present.<br>5. Very large cusp present. | 0: no hypocone expression of any form; a true three-cusped tooth.<br>1: for this grade, there is a low-level expression of the hypocone, often expressed as no more than an outline on the distolingual aspect of the trigon. In Dahlberg's original classification, this would be scored as a three-cusped upper molar along with grade 0.<br>2: in the Dahlberg classification, 3+ was equivalent to a small conical hypocone on the distolingual border of the trigon; grade 2 reflects this phenotype, where there is basically a conical cusp, or tubercle, with a free apex.<br>3: the hypocone is reduced in size but assumes a normal ovate shape along with a distinct free apex.<br>4: this grade would be equivalent to 3.5 on the modified hypocone plaque; the hypocone is reduced in size but is moderate rather than slight in expression.<br>5: hypocone is well developed, a step beyond grade 4.<br>6: pronounced expression of the hypocone; often equals or exceeds the size of the major cusps of the trigon. | 0 = 0T / 0S<br>1 = 1T / 1S<br>2 = 2T / 2S<br>3 = 3T / 3S<br>4 = 3.5T / 4S<br>5 = 4T / 5S<br>6 = 5T / 6S | 0. No hypocone.<br>1. Faint ridging present.<br>2. Faint cuspule present<br>3. Small cusp present.<br>4. Moderate-sized cusp present.<br>5. Large cusp present.<br>6. Very large cusp present. | 0: grades 0-2 of Turner et al., (1991)<br>1: grades 3-5 of Turner et al., (1991)<br><br>INFO PARA $M^2$:<br>*Presence corresponds to 4- and 4+ of Plaque P9, Dahlberg (1949) and grade 3 -5 of the ASU system.* | 0 = 0,1,2T / 0,1,2S / 0H<br>1 = 3,3.5,4,5T / 3,4,5,6S / 1H | 0. Absence: grades 0-2 of DeMoDa 1<br>1. Present: grades 3-6 of DeMoDa 1 |
| **INTERRUPTION GROOVE**<br><br>IG | **$I^1$ $I^2$ (T)**<br>**$I^1$ $I^2$ (S)** | 0. None. Mesial, distal, and medial areas of incisor lingual surface are smooth, continuous, and not disrupted by any vertical to near-horizontal groove.<br>M. An interruption groove occurs on the mesiolingual border.<br>D. An interruption groove occurs on the distolingual border.<br>MD. Grooves occur on both the mesio- and distolingual borders.<br>Med. A groove occurs in the medial area of the cingulum. | 0 = absence of grooves on lingual marginal ridges and basal cingula.<br>M = groove on mesiolingual marginal ridge.<br>D = groove on distolingual marginal ridge.<br>MD = grooves on both mesiolingual and distolingual marginal ridges.<br>med = groove on medial aspect of basal cingulum, sometimes extending onto root. | 0 = 0T / 0S<br>M = MT / MS<br>D = DT / DS<br>MD = MDT / MDS<br>Med = MedT / MedS | 0 = absence of grooves on lingual marginal ridges and basal cingula.<br>M = groove on mesiolingual marginal ridge.<br>D = groove on distolingual marginal ridge.<br>MD = grooves on both mesiolingual and distolingual marginal ridges.<br>med = groove on medial aspect of basal cingulum, sometimes extending onto root. | | 0 = 0T / 0S<br>M = MT / MS<br>D = DT / DS<br>MD = MDT / MDS<br>Med = MedT / MedS | 0 = absence of grooves on lingual marginal ridges and basal cingula<br>M = groove on mesiolingual marginal ridge<br>D = groove on distolingual marginal ridge<br>MD = grooves on both mesiolingual and distolingual marginal ridges<br>med = groove on medial aspect of basal cingulum, sometimes extending onto root |
| **LABIAL CONVEXITY**<br><br>LAB_CONV | **$I^1$ $I^2$ (T)**<br>**$I^1$ (S)** | 0. Labial surface is flat. There is no convexity at location approximately 1/3 from unworn occlusal surface, or 2/3 distance from crown-root junction.<br>1. Labial surface exhibits trace curvature.<br>2. Labial surface exhibits weak curvature.<br>3. Labial surface exhibits moderate curvature. | 0. Labial surface is flat.<br>1. Labial surface exhibits trace convexity.<br>2. Labial surface exhibits weak convexity.<br>3. Labial surface exhibits moderate convexity.<br>4. Labial surface exhibits pronounced convexity.<br>5. Labial surface exhibits very strong convexity. | 0 = 0T / 0S<br>1 = 1T / 1S<br>2 = 2T / 2S<br>3 = 3T / 3S<br>4 = 4T / 4,5S | 0. Labial surface is flat.<br>1. Labial surface exhibits trace convexity.<br>2. Labial surface exhibits weak convexity.<br>3. Labial surface exhibits moderate convexity.<br>4. Labial surface exhibits pronounced convexity. | | 0 = 0T / 0S<br>1 = 1T / 1S<br>2 = 2T / 2S<br>3 = 3T / 3S<br>4 = 4T / 4,5S | 0. Labial surface is flat.<br>1. Labial surface exhibits trace convexity.<br>2. Labial surface exhibits weak convexity.<br>3. Labial surface exhibits moderate convexity.<br>4. Labial surface exhibits pronounced convexity. |

| | | | | | | | | |
|---|---|---|---|---|---|---|---|---|
| | | 4. Labial surface exhibits strong curvature. | | | | | | |
| **LINGUAL CUSP (Premolar lingual cusp variation)**<br><br>LING_CUSP | **$P_3$ $P_4$ (T)**<br>**$P_3$ $P_4$ (S)** | A. No lingual cusp. A ridge may be present that suggests a much reduced structure without a free tip, but it is scored as cusp absent. Grade A was added after plaque production began when it was realized that lingual cusps can be absent.<br>0. One lingual cusp. Size and form may vary a great deal but tip can be seen.<br>1. One or two lingual cusps. This indecisive class should not be used for worn teeth. It is better to score such teeth as missing data.<br>2. Two lingual cusps. Mesial cusp is much larger than distal cusp.<br>3. Two lingual cusps. Mesial cusp is larger than lingual cusp.<br>4. Two lingual cusps. Mesial and distal cusps are equal in size.<br>5. Two lingual cusps. Distal cusp is larger than mesial cusp.<br>6. Two lingual cusps. Distal cusp is much larger than mesial cusp.<br>7. Two lingual cusps. Distal cusp is very much larger than lingual cusp. With wear this class can be confused with grade 0. When in doubt score individual as missing data.<br>8. Three lingual cusps. Each is about the same size.<br>9. Three lingual cusps. Mesial cusp is much larger than medial and/or distal cusp. With wear grade 9 can be confused with grade 3. When in doubt score such an individual as missing data. | $P_3$<br>0: lingual cusp has no free apex<br>1: single lingual cusp (on plaque, grades 0–1)<br>2: two lingual cusps (on plaque, grades 2–7)<br>3: three lingual cusps (on plaque, grades 8–9)<br>$P_4$<br>0: lingual cusp has no free apex<br>1: single lingual cusp (on plaque, grades 0–1)<br>2: two lingual cusps (on plaque, grades 2–7)<br>3: three lingual cusps (on plaque, grades 8–9) | 0 = AT / 0S<br>1 = 0,1T / 1S<br>2 = 2,3,4,5,6,7T / 2S<br>3 = 8,9T / 3S | 0: lingual cusp has no free apex<br>1: single lingual cusp (on plaque, grades 0–1)<br>2: two lingual cusps (on plaque, grades 2–7)<br>3: three lingual cusps (on plaque, grades 8–9) | | 0 = AT / 0S<br>1 = 0,1T / 1S<br>2 = 2,3,4,5,6,7T / 2S<br>3 = 8,9T / 3S | 0: lingual cusp has no free apex<br>1: single lingual cusp (on plaque, grades 0–1)<br>2: two lingual cusps (on plaque, grades 2–7)<br>3: three lingual cusps (on plaque, grades 8–9) |
| **MESIAL RIDGE (Canine mesial ridge. Bushman canine)**<br><br>MR | **C' (T)**<br>**C' (S)** | 0. Mesial and distal lingual ridges are the same size. Neither is attached to tuberculum dentale if it is present.<br>1. Neither is attached to tuberculum dentale if it is present.<br>2. Mesiolingual ridge is larger than the distolingual ridge, and moderately attached to tuberculum dentale.<br>3. Mesiolingual ridge is much more pronounced in size than the distolingual ridge and fully incorporated into tuberculum dentale. | 0: mesial and distal lingual ridges are the same size. Neither is attached to the tuberculum dentale, if present.<br>1: mesiolingual ridge is larger than distolingual and is weakly attached to the tuberculum dentale.<br>2: mesiolingual ridge is larger than the distolingual and is moderately attached to the tuberculum dentale.<br>3: Morris's type form. Mesiolingual ridge is much larger than the distolingual and is fully incorporated into the tuberculum dentale. | 0 = 0T / 0S<br>1 = 1T / 1S<br>2 = 2T / 2S<br>3 = 3T / 3S | 0. Mesial and distal lingual ridges are the same size. Neither is attached to tuberculum dentale if it is present.<br>1. Mesiolingual ridge is larger than the distolingual ridge, and weakly attached to tuberculum dentale.<br>2. Mesiolingual ridge is larger than the distolingual ridge, and moderately attached to tuberculum dentale.<br>3. Mesiolingual ridge is much more pronounced in size than the distolingual ridge and fully incorporated into tuberculum dentale (Morris´s type form). | | 0 = 0T / 0S<br>1 = 1T / 1S<br>2 = 2T / 2S<br>3 = 3T / 3S | 0. Mesial and distal lingual ridges are the same size. Neither is attached to tuberculum dentale if it is present.<br>1. Mesiolingual ridge is larger than the distolingual ridge, and weakly attached to tuberculum dentale.<br>2. Mesiolingual ridge is larger than the distolingual ridge, and moderately attached to tuberculum dentale.<br>3. Mesiolingual ridge is much more pronounced in size than the distolingual ridge and fully incorporated into tuberculum dentale (Morris´s type form). |
| **METACONE**<br><br>META | **$M^1$ $M^2$ $M^3$ (T)**<br>**$M^1$ $M^2$ $M^3$ (S)** | 0. Metacone (cusp 3) is absent.<br>1. An attached ridge is present at the metacone site, but there is no free cusp apex.<br>2. A faint cuspule with a free apex is present.<br>3. Weak cusp is present.<br>3.5. An intermediate-sized cusp is present (not shown on plaque).<br>4. Metacone is large.<br>5. Metacone is very large (equal to a large M1 hypocone). | 0: metacone absent<br>1: there is a ridge at the metacone site but no free apex<br>2: metacone expressed as faint cuspule with a free apex<br>3: weak cusp<br>3.5: intermediate-sized cusp that falls between grades 3 and 4 (interpolation necessary)<br>4: metacone is large<br>5: metacone is pronounced, equal in size to a large UM1 hypocone | 0 = 0T / 0S<br>1 = 1T / 1S<br>2 = 2T / 2S<br>3 = 3T / 3S<br>3.5 = 3.5T / 3.5S<br>4 = 4T / 4S<br>5 = 5T / 5S | 0. Metacone (cusp 3) is absent.<br>1. An attached ridge is present at the metacone site, but there is no free cusp apex.<br>2. A faint cuspule with a free apex is present.<br>3. Weak cusp is present.<br>3.5. An intermediate-sized cusp is present (not shown on plaque).<br>4. Metacone is large.<br>5. Metacone is very large (equal to a large M1 hypocone). | | 0 = 0T / 0S<br>1 = 1T / 1S<br>2 = 2T / 2S<br>3 = 3T / 3S<br>3.5 = 3.5T / 3.5S<br>4 = 4T / 4S<br>5 = 5T / 5S | 0. Metacone (cusp 3) is absent.<br>1. An attached ridge is present at the metacone site, but there is no free cusp apex.<br>2. A faint cuspule with a free apex is present.<br>3. Weak cusp is present.<br>3.5. An intermediate-sized cusp is present (not shown on plaque).<br>4. Metacone is large.<br>5. Metacone is very large (equal to a large M1 hypocone). |
| **ODONTOME**<br><br>ODONT | **$P^3$ $P^4$ (T)**<br>**$P_3$ $P_4$ (T)**<br>**$P^3$ $P^4$ (S)**<br>**$P_3$ $P_4$ (S)** | 0. Odontome not present.<br>1. Odontome present.<br>(NOTE: Turner often used a different scoring system that included the "+" sign. We have left them in the database, but we don't know what they mean, although they probably refer to "1" as he did not use that number on the templates). | 0: absence<br>1: odontome present in central sulcus (score all eight premolars) | 0 = 0T / 0S<br>1 = 1T / 1S | 0. Absent.<br>1. Present. | | 0 = 0T / 0S<br>1 = 1T / 1S | 0. Absent.<br>1. Present. |
| **PARASTYLE**<br><br>PARA | **$M^1$ $M^2$ $M^3$ (T)**<br>**$M^1$ $M^2$ $M^3$ (S)** | 0. The buccal surfaces of cusps 2 and 3 (paracone and metacone) are smooth.<br>1. A pit is present in or near the buccal groove between cusps 2 and 3.<br>2. Small cusp present with attached apex, usually on cusp 2.<br>3. Medium-sized cusp present with free apex anywhere on the buccal surface.<br>4. Large cusp present with free apex anywhere on the buccal surface.<br>5. Very large cusp present with free apex anywhere on the buccal surface.<br>6. An effectively free peg-shaped crown is present, attached to the third molar | 0: buccal surfaces of cusps 2 and 3 are smooth<br>1: a small pit near the buccal groove between cusps 2 and 3<br>2: small cusp but no free apex<br>3: medium cusp with free apex<br>4: large cusp with free apex<br>5: very large cusp with free apex that may extend onto the surfaces of both cusps 2 and 3<br>6: peg-shaped crown attached to root of second or third molar. This classic form of Bolk's paramolar tubercle may represent a supernumerary tooth that is fused to the buccal surface of UM2 or UM3. Accessory cusps with all the characteristics of a paramolar tubercle have also been observed on LM2 and LM3, adding evidence to the possibility these are fused supernumerary teeth. | 0 = 0T / 0S<br>1 = 1T / 1S<br>2 = 2T / 2S<br>3 = 3T / 3S<br>4 = 4T / 4S<br>5 = 5T / 5S<br>6 = 6T / 6S | 0. The buccal surfaces of cusps 2 and 3 (paracone and metacone) are smooth.<br>1. A pit is present in or near the buccal groove between cusps 2 and 3.<br>2. Small cusp present with attached apex, usually on cusp 2.<br>3. Medium-sized cusp present with free apex anywhere on the buccal surface.<br>4. Large cusp present with free apex anywhere on the buccal surface.<br>5. Very large cusp present with free apex anywhere on the buccal surface.<br>6. An effectively free peg-shaped crown is present, attached to the third molar root (not | | 0 = 0T / 0S<br>1 = 1T / 1S<br>2 = 2T / 2S<br>3 = 3T / 3S<br>4 = 4T / 4S<br>5 = 5T / 5S<br>6 = 6T / 6S | 0. The buccal surfaces of cusps 2 and 3 (paracone and metacone) are smooth.<br>1. A pit is present in or near the buccal groove between cusps 2 and 3.<br>2. Small cusp present with attached apex, usually on cusp 2.<br>3. Medium-sized cusp present with free apex anywhere on the buccal surface.<br>4. Large cusp present with free apex anywhere on the buccal surface.<br>5. Very large cusp present with free apex anywhere on the buccal surface.<br>6. An effectively free peg-shaped crown is present, attached to the third molar root (not given on plaque). This condition is extremely rare. |

| | | | | | | | | |
|---|---|---|---|---|---|---|---|---|
| | | root (not given on plaque). This condition is extremely rare. | | | given on plaque). This condition is extremely rare. | | | |
| **PEG-SHAPED (Peg-shaped incisor) UPPER LATERAL INCISOR VARIANTS**<br><br>PEG | **$I^2$ (T)<br>$I^2$ (S)** | 0. Normal-sized incisor.<br>1. Incisor reduced in size, but crown form is normal.<br>2. Incisor is very reduced in size, lacks appropriate morphology, being instead peg-shaped.<br>(NOTE: some Turner datasheets scored this as "0", "R", "P". We translated this to "0", "1" and "2" respectively) | 0: UI2 normal in form and size.<br>1: UI2 normal in form but diminutive in size (less than ½ mesiodistal diameter of UI1).<br>2: congenital absence.<br>3: peg-shaped UI2, conical in form, often with no morphological features.<br>4: talon cusp (in same location but much more pronounced and distinctive than a tuberculum dentale).<br>5: triform UI2 with a large lingual structure that runs from basal cingulum to incisal edge.<br>6: unusual UI2 forms that do not fit any of the above categories. | 0 = 0T/ 0S<br>1 = 1T /1S<br>2 = 2T / 3,4,5,6S | 0. Normal-sized incisor.<br>1. Incisor reduced in size, but crown form is normal.<br>2. Incisor is very reduced in size, lacks appropriate morphology, being instead peg-shaped. | | 0 = 0T/ 0S<br>1 = 1T /1S<br>2 = 2T / 3,4,5,6S | 0. Normal-sized incisor.<br>1. Incisor reduced in size, but crown form is normal.<br>2. Incisor is very reduced in size, lacks appropriate morphology, being instead peg-shaped. |
| **PEG-SHAPED (Peg-shaped molar)**<br><br>PEG | **$M^3$ (T)<br>$M^3$ (S)** | 0. Full-sized crown with normal third molar morphology.<br>1. Molar reduced in size to 7 to 10 mm linguobuccal diameter. Form is near normal or somewhat shriveled.<br>2. Molar is less than 7 mm in linguobuccal diameter. Crown is peg or cone-shaped with rarely more than two rounded cusps lacking secondary morphology. Root is simple and single.<br>(NOTE: some Turner datasheets scored this as "0", "R", "P". We translated this to "0", "1" and "2" respectively). | 0: third molar present and normal.<br>1: third molar significantly reduced in size (ca. ½ normal size, with two or more cusps).<br>2: third molar peg-shaped (only a single cusp evident)<br>3: third molar congenitally absent. | 0 = 0T/ 0S<br>1 = 1T /1S<br>2 = 2T / 2S | 0. Full-sized crown with normal third molar morphology.<br>1. Molar reduced in size to 7 to 10 mm linguobuccal diameter. Form is near normal or somewhat shriveled.<br>2. Molar is less than 7 mm in linguobuccal diameter. Crown is peg or cone-shaped with rarely more than two rounded cusps lacking secondary morphology. Root is simple and single. | | 0 = 0T/ 0S<br>1 = 1T /1S<br>2 = 2T / 2S | 0. Full-sized crown with normal third molar morphology.<br>1. Molar reduced in size to 7 to 10 mm linguobuccal diameter. Form is near normal or somewhat shriveled.<br>2. Molar is less than 7 mm in linguobuccal diameter. Crown is peg or cone-shaped with rarely more than two rounded cusps lacking secondary morphology. Root is simple and single. |
| **PROTOSTYLID**<br><br>PROTO | **$M_2$ $M_3$ (T)<br>$M_2$ $M_3$ (S)** | 0. No expression of any sort. Buccal surface is smooth.<br>1. A pit occurs in the buccal groove separating cusps 1 and 3.<br>2. Buccal groove tends to curve distalward.<br>3. A faint groove occurs extending mesialward from the buccal groove.<br>4. Groove is slightly more pronounced.<br>5. Groove is stronger and can be easily seen.<br>6. Groove extends across most of the buccal surface of cusp 1. This is considered a weak or small cusp.<br>7. A cusp occurs with a free (unattached) cusp tip.<br>(NOTE: in one individual, Turner scored this trait with a "4*" and "5*". We don't know what they mean, but we assume that similar to Carabelli´s trait, there is probably a pit in the buccal groove) | 0: no pit or positive expression on buccal surface of lower molar<br>1: buccal pit (a pit of varying sizes, situated around the midpoint of the crown in the protoconid–hypoconid inter-lobal groove)<br>2: a very slight swelling and associated groove coursing mesially from buccal groove<br>3: slight positive expression on mesiobuccal cusp<br>4: moderate positive expression<br>5: strong positive expression<br>6: pronounced positive expression<br>7: most distinctive form of protostylid, expressed as tubercle | 0 = 0T / 0S<br>1 = 1T / 1S<br>2 = 2T / 2S<br>3 = 3T / 3S<br>4 = 4T / 4S<br>5 = 5T / 5S<br>6 = 6T / 6S<br>7 = 7T / 7S | 0. No expression of any sort. Buccal surface is smooth.<br>1. A pit occurs in the buccal groove separating cusps 1 and 3.<br>2. Buccal groove tends to curve distalward.<br>3. A faint groove occurs extending mesialward from the buccal groove.<br>4. Groove is slightly more pronounced.<br>5. Groove is stronger and can be easily seen.<br>6. Groove extends across most of the buccal surface of cusp 1. This is considered a weak or small cusp.<br>7. A cusp occurs with a free (unattached) cusp tip. | | 0 = 0T / 0S<br>1 = 1T / 1S<br>2 = 2T / 2S<br>3 = 3T / 3S<br>4 = 4T / 4S<br>5 = 5T / 5S<br>6 = 6T / 6S<br>7 = 7T / 7S | 0. No expression of any sort. Buccal surface is smooth.<br>1. A pit occurs in the buccal groove separating cusps 1 and 3.<br>2. Buccal groove tends to curve distalward.<br>3. A faint groove occurs extending mesialward from the buccal groove.<br>4. Groove is slightly more pronounced.<br>5. Groove is stronger and can be easily seen.<br>6. Groove extends across most of the buccal surface of cusp 1. This is considered a weak or small cusp.<br>7. A cusp occurs with a free (unattached) cusp tip. |
| | **$M_1$ (T)<br>$M_1$ (S)<br>$M_1$ (H)** | 0. No expression of any sort. Buccal surface is smooth.<br>1. A pit occurs in the buccal groove separating cusps 1 and 3.<br>2. Buccal groove tends to curve distalward.<br>3. A faint groove occurs extending mesialward from the buccal groove.<br>4. Groove is slightly more pronounced.<br>5. Groove is stronger and can be easily seen.<br>6. Groove extends across most of the buccal surface of cusp 1. This is considered a weak or small cusp.<br>7. A cusp occurs with a free (unattached) cusp tip.<br>(NOTE: in one individual, Turner scored this trait with a "4*" and "5*". We don't know what they mean, but we assume that similar to Carabelli´s trait, there is probably a pit in the buccal groove). | 0: no pit or positive expression on buccal surface of lower molar<br>1: buccal pit (a pit of varying sizes, situated around the midpoint of the crown in the protoconid–hypoconid inter-lobal groove)<br>2: a very slight swelling and associated groove coursing mesially from buccal groove<br>3: slight positive expression on mesiobuccal cusp<br>4: moderate positive expression<br>5: strong positive expression<br>6: pronounced positive expression<br>7: most distinctive form of protostylid, expressed as tubercle | 0 = 0T / 0S<br>1 = 1T / 1S<br>2 = 2T / 2S<br>3 = 3T / 3S<br>4 = 4T / 4S<br>5 = 5T / 5S<br>6 = 6T / 6S<br>7 = 7T / 7S | 0. No expression of any sort. Buccal surface is smooth.<br>1. A pit occurs in the buccal groove separating cusps 1 and 3.<br>2. Buccal groove tends to curve distalward.<br>3. A faint groove occurs extending mesialward from the buccal groove.<br>4. Groove is slightly more pronounced.<br>5. Groove is stronger and can be easily seen.<br>6. Groove extends across most of the buccal surface of cusp 1. This is considered a weak or small cusp.<br>7. A cusp occurs with a free (unattached) cusp tip. | 0: grades 0-1 of Turner et al., (1991)<br>1: grades 2-7 of Turner et al., (1991)<br><br>INFO PARA $M_1$:<br>*The ASU system ' s 2‑7 were scored as present.* | 0 = 0,1T / 0,1S / 0H<br>1 = 2,3,4,5,6,7T / 2,3,4,5,6,7S / 1H | 0. Absence: grades 0-1 of DeMoDa 1<br>1. Present: grades 2-7 of DeMoDa 1 |
| **RADICAL NUMBER**<br><br>RADN | **$I^1$ $I^2$ C' (T)<br>$P^3$ $P^4$ (T)<br>$M^1$ $M^2$ $M^3$ (T)<br><br>$I_1$ $I_2$ C, (T)<br>$P_3$ $P_4$ (T)<br>$M_1$ $M_2$ $M_3$ (T)** | 1. One radical. No developmental grooves.<br>2. Two radicals. Two developmental grooves or two round roots with no developmental grooves.<br>3. Three radicals. Three developmental grooves or one round root with no developmental grooves and one root with two developmental grooves.<br>4. Four radicals. Continuation of above with various root number and developmental groove combinations.<br>5. Five radicals. Continuation of above.<br>6. Six radicals. Continuation of above.<br>7. Seven radicals. Continuation of above.<br>8. Eight radicals. Continuation of above. | | 0 = 0T<br>1 = 1T<br>2 = 2T<br>3 = 3T<br>4 = 4T<br>5 = 5T<br>6 = 6T<br>7 = 7T<br>8 = 8T | 1. One radical. No developmental grooves.<br>2. Two radicals. Two developmental grooves or two round roots with no developmental grooves.<br>3. Three radicals. Three developmental grooves or one round root with no developmental grooves and one root with two developmental grooves.<br>4. Four radicals. Continuation of above with various root number and developmental groove combinations.<br>5. Five radicals. Continuation of above.<br>6. Six radicals. Continuation of above.<br>7. Seven radicals. Continuation of above.<br>8. Eight radicals. Continuation of above. | | 0 = 0T<br>1 = 1T<br>2 = 2T<br>3 = 3T<br>4 = 4T<br>5 = 5T<br>6 = 6T<br>7 = 7T<br>8 = 8T | 1. One radical. No developmental grooves.<br>2. Two radicals. Two developmental grooves or two round roots with no developmental grooves.<br>3. Three radicals. Three developmental grooves or one round root with no developmental grooves and one root with two developmental grooves.<br>4. Four radicals. Continuation of above with various root number and developmental groove combinations.<br>5. Five radicals. Continuation of above.<br>6. Six radicals. Continuation of above.<br>7. Seven radicals. Continuation of above.<br>8. Eight radicals. Continuation of above. |

| | | | | | | | | |
|---|---|---|---|---|---|---|---|---|
| **ROOT NUMBER**<br>**(Canine root number)**<br><br>ROOTN | **C, (T)**<br>**C, (S)** | 1. One root.<br>2. Two roots, free more than 1/4 to 1/3 total root length.<br>(NOTE: sometimes Turner scored this trait with 1* which means "bifid tip") | 0: one-rooted LC, with or without root grooves separating buccal and lingual cones<br>1: two-rooted LC, with inter-radicular projection separating buccal and lingual cones by at least ¼ to ⅓ of total root length | 1 = 1T / 0S<br>2 = 2T / 1S | 1. One root.<br>2. Two roots, free more than 1/4 to 1/3 total root length. | | 1 = 1T / 0S<br>2 = 2T / 1S | 1. One root.<br>2. Two roots, free more than 1/4 to 1/3 total root length. |
| **ROOT NUMBER**<br>**(Premolar root number)**<br><br>ROOTN | **$P^3 P^4$ (T)**<br>**$P^3 P^4$ (S)** | 1. One root. Tip may be bifurcated.<br>2. Two roots. Separate roots must be greater than 1/4 to 1/3 of total root length.<br>3. Three roots. Length defined as above.<br>(NOTE: sometimes Turner scored this trait with a number and asterisk. "1*" means "bifid tip" and "2*" means trifid tip) | 1: one-rooted (root grooves separate cones but no inter-radicular projection)<br>2: two-rooted (inter-radicular projection separates buccal and lingual root cones for ¼ to ⅓ of total root length)<br>3: three-rooted (there is an inter-radicular projection that separates the buccal root into two distinct roots, and another projection separating the two buccal roots from a single lingual root) | 0 = 0T / 0S<br>1 = 1T / 1S<br>2 = 2T / 2S<br>3 = 3T / 3S | 1. One root. Tip may be bifurcated.<br>2. Two roots. Separate roots must be greater than 1/4 to 1/3 of total root length.<br>3. Three roots. Length defined as above. | | 0 = 0T / 0S<br>1 = 1T / 1S<br>2 = 2T / 2S<br>3 = 3T / 3S | 1. One root. Tip may be bifurcated.<br>2. Two roots. Separate roots must be greater than 1/4 to 1/3 of total root length.<br>3. Three roots. Length defined as above. |
| **ROOT NUMBER**<br>**(Upper molar root number)**<br><br>ROOTN | **$M^1 M^2 M^3$ (T)**<br>**$M^1 M^2 M^3$ (S)** | 1. One root. Tip may be bifurcated with deeply inset developmental grooves.<br>2. Two roots. Separate roots are greater than 1/4 to 1/3 of total root length. Length determination should take into account bending which is common on third molars.<br>3. Three roots. Length defined as above.<br>4. Four roots. Length defined as above. | 1: one-rooted (root cones separated by grooves but there are no inter-radicular projections)<br>2: two-rooted (one inter-radicular projection separates one root from two fused roots)<br>3: three-rooted (three inter-radicular projections separate all three roots for at least ¼ to ⅓ of total root length) | 1 = 1T / 1S<br>2 = 2T / 2S<br>3 = 3,4T / 3S | 1. One root. Tip may be bifurcated with deeply inset developmental grooves.<br>2. Two roots. Separate roots are greater than 1/4 to 1/3 of total root length. Length determination should take into account bending which is common on third molars.<br>3. Three roots. Length defined as above. | | 1 = 1T / 1S<br>2 = 2T / 2S<br>3 = 3,4T / 3S | 1. One root. Tip may be bifurcated with deeply inset developmental grooves.<br>2. Two roots. Separate roots are greater than 1/4 to 1/3 of total root length. Length determination should take into account bending which is common on third molars.<br>3. Three roots. Length defined as above. |
| **ROOT NUMBER**<br>**(Lower molar root number)**<br><br>ROOTN | **$M_1 M_2 M_3$ (T)**<br>**$M_1 M_2 M_3$ (S)** | 1. One root. Root will usually be U-shaped in cross section with a deep developmental groove in the lingual surface. Root tip may be bifurcated. If tips are free more than ¼ to 1/3 of the total root length, score as two-rooted.<br>2. Two roots. Two separate roots exist for at least 1/4 to 1/3 of total root length. A strong distolingual radical is likely an unattached supernumerary third root.<br>3. Three roots. A third (supernumerary) root is present on the distolingual aspect. It may be very small, but usually is about 1/3 the size of the normal distal root. | 1: one-rooted lower molar (mesial and distal roots of lower molars can be fused on either buccal or lingual aspect or both)<br>2: inter-radicular structure produces clear separation of mesial and distal roots for at least ¼ to ⅓ of total root length | 1 = 1T / 0S<br>2 = 2T / 1S<br>3 = 3T | 1. One root. Root will usually be U-shaped in cross section with a deep developmental groove in the lingual surface. Root tip may be bifurcated. If tips are free more than ¼ to 1/3 of the total root length, score as two-rooted.<br>2. Two roots. Two separate roots exist for at least 1/4 to 1/3 of total root length. A strong distolingual radical is likely an unattached supernumerary third root.<br>3. Three roots. A third (supernumerary) root is present on the distolingual aspect. It may be very small, but usually is about 1/3 the size of the normal distal root. | | 1 = 1T / 0S<br>2 = 2T / 1S<br>3 = 3T | 1. One root. Root will usually be U-shaped in cross section with a deep developmental groove in the lingual surface. Root tip may be bifurcated. If tips are free more than ¼ to 1/3 of the total root length, score as two-rooted.<br>2. Two roots. Two separate roots exist for at least 1/4 to 1/3 of total root length. A strong distolingual radical is likely an unattached supernumerary third root.<br>3. Three roots. A third (supernumerary) root is present on the distolingual aspect. It may be very small, but usually is about 1/3 the size of the normal distal root. |
| | **C' (T)**<br>**$I_1 I_2$ (T)**<br>**C' (S)**<br>**$I_1 I_2$ C, (S)** | C'<br>0. None. Lingual surface is essentially flat.<br>1. Faint. Very slight elevations of mesial and distal aspects of lingual surface can be seen and palpated.<br>2. Trace. Elevations are easily seen. This grade probably considered minimal expression by most observers.<br>3. Semi-shovel. Stronger ridging is present and there is a tendency for ridge convergence at the cingulum.<br>4. Semi-shovel. Convergence and ridging are stronger than in grade 3.<br>5. Shovel. Strong development of ridges, which almost contact at cingulum.<br>6. Marked shovel. Strongest development. Mesial and distal lingual ridges are sometimes in contact at the cingulum.<br>$I_1 I_2$<br>0. None. Lingual surface is essentially flat.<br>1. Faint. Very slight elevations of mesial and distal aspects of lingual surface can be seen and palpated.<br>2. Trace. Elevations are easily seen. This grade probably considered minimal expression by most observers.<br>3. Semi-shovel. Stronger ridging is present and there is a tendency for ridge convergence at the cingulum.<br>(NOTE: Turner only scored the lower shoveling for the lower central incisors: $I_{1S}$) | C'<br>0. None. Lingual surface is essentially flat.<br>1. Faint. Very slight elevations of mesial and distal aspects of lingual surface can be seen and palpated.<br>2. Trace. Elevations are easily seen. This grade probably considered minimal expression by most observers.<br>3. Semi-shovel. Stronger ridging is present and there is a tendency for ridge convergence at the cingulum.<br>4. Semi-shovel. Convergence and ridging are stronger than in grade 3.<br>5. Shovel. Strong development of ridges, which almost contact at cingulum.<br>6. Marked shovel. Strongest development. Mesial and distal lingual ridges are sometimes in contact at the cingulum.<br>$I_1 I_2$ C,<br>0. None. Lingual surface is essentially flat.<br>1. Faint. Very slight elevations of mesial and distal aspects of lingual surface can be seen and palpated.<br>2. Trace. Elevations are easily seen. This grade probably considered minimal expression by most observers.<br>3. Semi-shovel. Stronger ridging is present and there is a tendency for ridge convergence at the cingulum. | 0 = 0T / 0S<br>1 = 1T / 1S<br>2 = 2T / 2S<br>3 = 3T / 3S<br>4 = 4T / 4S<br>5 = 5T / 5S<br>6 = 6T / 6S | 0. None. Lingual surface is essentially flat.<br>1. Faint. Very slight elevations of mesial and distal aspects of lingual surface can be seen and palpated.<br>2. Trace. Elevations are easily seen. This grade probably considered minimal expression by most observers.<br>3. Semi-shovel. Stronger ridging is present and there is a tendency for ridge convergence at the cingulum.<br>4. Semi-shovel. Convergence and ridging are stronger than in grade 3.<br>5. Shovel. Strong development of ridges, which almost contact at cingulum.<br>6. Marked shovel. Strongest development. Mesial and distal lingual ridges are sometimes in contact at the cingulum. | | 0 = 0T / 0S<br>1 = 1T / 1S<br>2 = 2T / 2S<br>3 = 3T / 3S<br>4 = 4T / 4S<br>5 = 5T / 5S<br>6 = 6T / 6S | 0. None. Lingual surface is essentially flat.<br>1. Faint. Very slight elevations of mesial and distal aspects of lingual surface can be seen and palpated.<br>2. Trace. Elevations are easily seen. This grade probably considered minimal expression by most observers.<br>3. Semi-shovel. Stronger ridging is present and there is a tendency for ridge convergence at the cingulum.<br>4. Semi-shovel. Convergence and ridging are stronger than in grade 3.<br>5. Shovel. Strong development of ridges, which almost contact at cingulum.<br>6. Marked shovel. Strongest development. Mesial and distal lingual ridges are sometimes in contact at the cingulum. |
| **SHOVELING** | **$I^1 I^2$ (T)**<br>**$I^1 I^2$ (S)**<br>**$I^1 I^2$ (H)** | 0. None. Lingual surface is essentially flat.<br>1. Faint. Very slight elevations of mesial and distal aspects of lingual surface can be seen and palpated.<br>2. Trace. Elevations are easily seen. This grade probably considered minimal expression by most observers.<br>3. Semi-shovel. Stronger ridging is present and there is a tendency for ridge convergence at the cingulum. | $I^1$<br>0 (absence): it is rare for UI1 to express the complete absence of marginal ridges (see Figure 4.1a for example). For this reason, grade 0 on the UI1 shoveling plaque actually shows very slight marginal ridge expression.<br>1 (trace): marginal ridges can be discerned, but expression is slight, with mesial marginal ridge not extending to the basal eminence. | 0 = 0T / 0S<br>1 = 1T / 1S<br>2 = 2T / 2S<br>3 = 3T / 3S<br>4 = 4T / 4S<br>5 = 5T / 5S<br>6 = 6T / 6S<br>7 = 7T / 7S | 0. None. Lingual surface is essentially flat.<br>1. Faint. Very slight elevations of mesial and distal aspects of lingual surface can be seen and palpated.<br>2. Trace. Elevations are easily seen. This grade probably considered minimal expression by most observers.<br>3. Semi-shovel. Stronger ridging is present and there is a tendency for ridge convergence at the cingulum. | 0: grades 0-2 of Turner et al., (1991)<br>1: grades 3-7 of Turner et al., (1991)<br><br>INFO PARA $I^1$:<br>*The depth of Lingual fossa deeper than 0.5 mm is scored as present. This classification corresponds to the categories of semi, moderate, and strong shovel employed by Hanihara et al. (1970), and grade 3‑7 of the Arizona State University Dental* | 0 = 0-1-2T / 0-1-2S / 0H<br>1 = 3-4-5-6-7T / 3-4-5-6-7S / 1H | 0. Absence: grades 0-2 of DeMoDa 1<br>1. Present: grades 3-7 of DeMoDa 1 |

| | | | | | | | | |
|---|---|---|---|---|---|---|---|---|
| SHOV | | 4. Semi-shovel. Convergence and ridging are stronger than in grade 3.<br>5. Shovel. Strong development of ridges, which almost contact at cingulum.<br>6. Marked shovel. Strongest development. Mesial and distal lingual ridges are sometimes in contact at the cingulum.<br>7. Barrel. Expression exceeds grade 6 (only $I^2$) | 2 (low moderate): ridges more pronounced, with mesial marginal ridge extending further down on basal eminence.<br>3 (high moderate): ridges more pronounced, almost coalescing at basal eminence.<br>4 (low pronounced): well developed ridges that converge at basal eminence.<br>5 (medium pronounced): more pronounced marginal ridges meeting at basal eminence.<br>6 (high pronounced): pronounced ridges that meet at basal eminence, almost folding around on themselves.<br>7 (extreme pronounced): any expression that exceeds grade 6 can be placed in grade 7. It is a rare expression, so much so that a good example for the plaque was never found. This grade would involve marginal ridges that folded around on themselves, similar to grade 6 on the UI2 shoveling plaque.<br>$I^2$<br>0 (absence): as for UI1, complete absence of marginal ridges is rare on UI2. There are slight marginal ridges on the plaque for grade 0.<br>1 (trace): characterized by the presence of faint mesial and distal marginal ridges.<br>2 (low moderate): moderate marginal ridges with little fossa formation.<br>3 (high moderate): distinct marginal ridges but only moderate lingual fossa.<br>4 (low pronounced): well developed marginal ridges that come in contact at the lingual base of the crown.<br>5 (medium pronounced): well developed marginal ridges, forming a distinct lingual fossa.<br>6 (semi-barreled): the marginal ridges wrap around and contact at a low point on the basal eminence.<br>7 (barreled): the marginal ridges are so pronounced that they contact at almost the incisal surface of the basal eminence, assuming a full barrel shape. | | 4. Semi-shovel. Convergence and ridging are stronger than in grade 3.<br>5. Shovel. Strong development of ridges, which almost contact at cingulum.<br>6. Marked shovel. Strongest development. Mesial and distal lingual ridges are sometimes in contact at the cingulum.<br>7. Barrel. Expression exceeds grade 6. | *Anthropology System (ASU system) (Turner et al., 1991; Scott and Turner, 1997).*<br><br>INFO PARA $I^2$:<br>*The teeth classified as the ASU system's grade 3-7 are scored as present.* | | |
| **TOME'S ROOT LOWER FIRST PREMOLAR ROOT NUMBER**<br><br>TOMES | **$P_3$ (T)**<br>**$P_3$ (S)** | 0. Developmental grooving is absent, or if present, shallow with rounded rather than V-shaped indentation.<br>1. Developmental groove is present and has a shallow V-shaped cross-section.<br>2. Developmental groove is present and has a moderately deep V-shaped cross-section.<br>3. Developmental groove is present, V-shaped, and deep. Groove extends at least 1/3 of total root length.<br>4. Developmental grooving is deeply invaginated on both the mesial and distal root surfaces.<br>5. Two free roots are present. Their length is at least 1/4 to 1/3 of the total root length. | 0: slight or no groove separating cones on mesial surface of LP1 root<br>1: slight V-shaped groove separating cones<br>2: deeper V-shaped groove separating cones<br>3: deep developmental groove separating root cones along at least ⅓ of root<br>4: deep grooving on both mesial and distal surfaces of root<br>5: inter-radicular projection present so LP1 has two roots, a large buccal root and a smaller mesial/lingual root | 0 = 0T / 0S<br>1 = 1T / 1S<br>2 = 2T / 2S<br>3 = 3T / 3S<br>4 = 4T / 4S<br>5 = 5T / 5S | 0: slight or no groove separating cones on mesial surface of LP1 root<br>1: slight V-shaped groove separating cones<br>2: deeper V-shaped groove separating cones<br>3: deep developmental groove separating root cones along at least ⅓ of root<br>4: deep grooving on both mesial and distal surfaces of root<br>5: inter-radicular projection present so LP1 has two roots, a large buccal root and a smaller mesial/lingual root | | 0 = 0T / 0S<br>1 = 1T / 1S<br>2 = 2T / 2S<br>3 = 3T / 3S<br>4 = 4T / 4S<br>5 = 5T / 5S | 0: slight or no groove separating cones on mesial surface of LP1 root<br>1: slight V-shaped groove separating cones<br>2: deeper V-shaped groove separating cones<br>3: deep developmental groove separating root cones along at least ⅓ of root<br>4: deep grooving on both mesial and distal surfaces of root<br>5: inter-radicular projection present so LP1 has two roots, a large buccal root and a smaller mesial/lingual root |
| **TORSOMOLAR ANGLE**<br><br>TORSO | **$M_3$ (T)**<br>**$M_3$ (S)** | When the third molar has rotated lingualward or buccalward relative to a line drawn through the middle of the first and second molars.<br>(NOTE: to simplify the angle measurements that Turner wrote on the templates, we converted these angles to the 0 and 1 method of Scott and Irish, 2017) | 0: absent (torsomolar angle <10°)<br>1: present (torsomolar angle ≥10° in either buccal or lingual direction) | 0 = 0S<br>1 = 1S | 0: absent (torsomolar angle <10°)<br>1: present (torsomolar angle ≥10° in either buccal or lingual direction) | | 0 = 0S<br>1 = 1S | 0: absent (torsomolar angle <10°)<br>1: present (torsomolar angle ≥10° in either buccal or lingual direction) |
| **TUBERCULUM DENTALE**<br><br>TD | **$I^1$ $I^2$ C' (T)**<br>**$I^1$ $I^2$ C' (S)** | 0. No expression. Cingular region of lingual surface is smooth. Ignore any shoveling presence.<br>1. Faint ridging. Matches grade 1 of ASU U11 t.d. plaque.<br>2. Trace ridging. Matches grade 2 of ASU U11 t.d. plaque.<br>3. Strong ridging. Matches grade 3 of ASU U11 t.d. plaque.<br>4. Pronounced ridging. Matches grade 4 of ASU U11 t.d. plaque.<br>5-. A weakly developed cuspule is attached to either the mesio- or distolingual ridge. Cuspule apex is not free.<br>5. Weakly developed cuspule with a free apex. Size corresponds approximately with ASU UC DAR plaque grade 4 tuberculum dentale.<br>6. Strong cusp with free apex. Size is equal to or greater than the ASU UC DAR plaque grade 5 tuberculum dentale. | $I^1$<br>0: no ridges on basal eminence<br>1: one slight ridge<br>2: two slight ridges, or one moderate ridge<br>3: one moderate ridge and one slight ridge, or one pronounced ridge<br>4: two moderate ridges, or one slight and one pronounced ridge<br>5: one moderate ridge and one pronounced ridge<br>6: two pronounced ridges, or three slight and/or moderate ridges<br>$I^2$<br>0: absence of ridge or tubercle formation<br>1: slight ridge<br>2: moderate ridge<br>3: small tubercle<br>4: moderate tubercle<br>5: large or double tubercle<br>6: welt on basal cingulum (distinct from talon cusp; see Trait 23)<br>C'<br>0: absence of any ridge or tubercle formation<br>1: very slight tubercle, characterized by a single groove | 0 = 0T / 0S<br>1 = 1T / 1S<br>2 = 2T / 2S<br>3 = 3T / 3S<br>4 = 4T / 4S<br>5 = 5-,5T / 5S<br>6 = 6T / 6,7S | 0. No expression. Cingular region of lingual surface is smooth.<br>1. Faint ridging.<br>2. Trace ridging.<br>3. Strong ridging.<br>4. Pronounced ridging.<br>5. Weakly developed cuspule with a free apex.<br>6. Strong cusp with free apex. | | 0 = 0T / 0S<br>1 = 1T / 1S<br>2 = 2T / 2S<br>3 = 3T / 3S<br>4 = 4T / 4S<br>5 = 5,5-T / 5S<br>6 = 6T / 6,7S | 0. No expression. Cingular region of lingual surface is smooth.<br>1. Faint ridging.<br>2. Trace ridging.<br>3. Strong ridging.<br>4. Pronounced ridging.<br>5. Weakly developed cuspule with a free apex.<br>6. Strong cusp with free apex. |

| | | | | | | | | |
|---|---|---|---|---|---|---|---|---|
| | | (NOTE: sometimes Turner scored this trait with “DAR4” and “DAR5”, but we converted these to grade 5 and grade 6 respectively. Moreover, Turner also scored this feature with “<1”. We don’t know what they mean, but they probably indicate a tuberculum dentale smaller than faint ridging) | 2: slight tubercle, outlined by two grooves<br>3: moderate tubercle<br>4: medium tubercle with no free apex<br>5: large tubercle with free apex<br>6: pronounced tubercle with free apex<br>7: hyper-pronounced tubercle with free apex | | | | | |
| **WINGING**<br><br>WING | **$I^1$ (T)**<br>**$I^1$ (S)**<br>**$I_1$ (S)** | 1. Bilateral winging. Central incisors are rotated mesiolingualward, giving a V-shaped appearance when viewed from the occlusal surface. When the angle formed is greater than 20 degrees, it is classed as 1A; when less than 20 degrees, 1B.<br>2. Unilateral winging. Only one of the incisors is rotated. The other is straight. No subclasses are recognized.<br>3. Straight. Both teeth form a straight labial surface, or follow the curvature of the dental arcade.<br>4. Counter-winging. One or both teeth are rotated distolingualward. | 0 (absence): if the line is parallel to the labial surfaces or if the distal margins fall below the line, winging is absent. Angle ≥180°.<br>1 (trace winging): the mesial margins of the upper incisors fall slightly below the line. Angle 160–180°.<br>2 (moderate winging): the mesial margins are more removed from the line. Angle 135–159°.<br>3 (pronounced winging): there is a distinct distance between the line and the mesial margins. Angle <135°. | 0 = 2,3,4T / 0S<br>1 = 1,1A T / 1S<br>2 = 1B T / 2,3S | 0: absent.<br>1: trace winging (angle 160–180°) the mesial margins of the upper incisors fall slightly below the line.<br>2: moderate (angle 135–159°) or pronounced (angle <135°) winging. | | 0 = 2,3,4T / 0S<br>1 = 1,1A T / 1S<br>2 = 1B T / 2,3S | 0: absent<br>1: trace winging (angle 160–180°) the mesial margins of the upper incisors fall slightly below the line.<br>2: moderate (angle 135–159°) or pronounced (angle <135°) winging. |

* (T) are the teeth scored according to Turner et al., 1991 system.
(S) are the teeth scored according to Scott and Irish, 2017 system.
(H) are the teeth scored according to Hanihara 2008 system.


**Hanihara, T., 2008.** Morphological variation of major human populations based on nonmetric dental traits. American Journal of Physical Anthropology. 136:169-182

**Scott, G.R., Irish, J.D., 2017.** Human Tooth Crown and Root Morphology: The Arizona State University Dental Anthropology System. Cambridge University Press.

**Turner, C.G., Nichol, C.R., Scott, G.R., 1991.** Scoring procedures for key morphological traits of the permanent dentition: the Arizona State University Dental Anthropology System. In: Kelley, M., Larsen, C. (Eds.), Advances in Dental Anthropology. Wiley-Liss, New York, pp. 13–31.
